## Supplemental Material for "Multi-Coloured Sequential Resonance Energy Transfer for Simultaneous Ligand Binding at G Protein-Coupled Receptors"

### Contents

**Supplemental Table 1** 2
**Supplemental Table 2** 4
**Supplemental Table 3** 5
**Supplemental Table 4** 6
**Supplemental Table 5** 7
**Supplemental Figure 1.** 8
**Supplemental Figure 2.** 9
**Supplemental Figure 3.** 10
**Supplemental Figure 4.** 11
**Supplemental Figure 5.** 12
**Supplemental Figure 6.** 14
**Supplemental Figure 7.** 13
**Supplemental Figure 8.** 15
**Supplemental Figure 9.** 16
**Supplemental Figure 10.** 17
**Supplemental Figure 11.** 18
**Supplemental Chemistry Methods** 19
**NMR Spectra and Analytical HPLC Traces** 61

***Supplemental Table 1.*** *FFA1 Site two Tracer and Precursor $Ca^{2+}$ Data*

| ID | Structure | $pEC_{50}$ | EMax[a] |
|---|---|---|---|
| T360 | 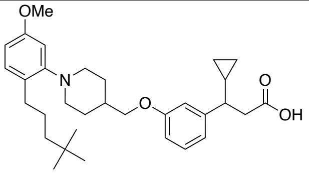 | 7.48±0.31 | 105±5 |
| Cpd 6h | 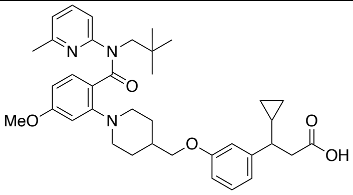 | 7.58±0.37 | 108±8 |
| TUG-2450 | 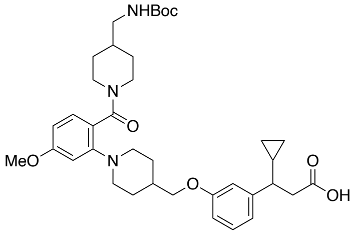 | 6.18±0.09 | 108±7 |
| TUG-2451 | 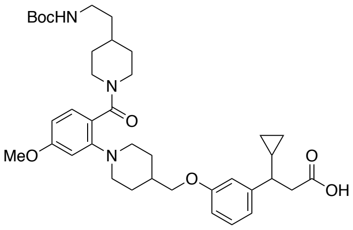 | 6.20±0.14 | 112±10 |
| TUG-2458 | 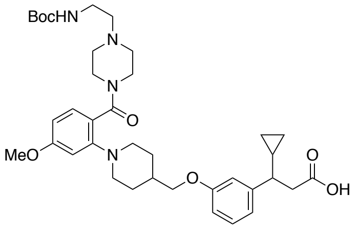 | 5.38±0.09 | 133±5 |
| TUG-2455 | 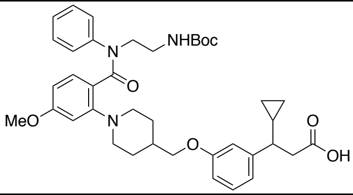 | 6.63±0.10 | 103±12 |
| TUG-2457 | 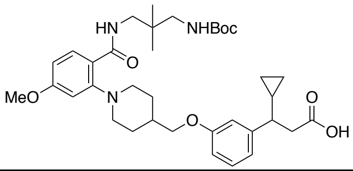 | 6.62±0.15 | 108±3 |
| TUG-2489 | 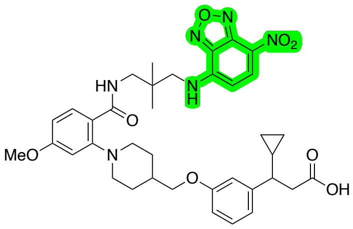 | 5.29±0.04 | 150±15 |

| | | | |
|---|---|---|---|
| TUG-2467 | 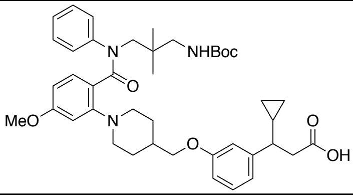 | 6.78±0.02 | 124±9 |
| TUG-2490 | 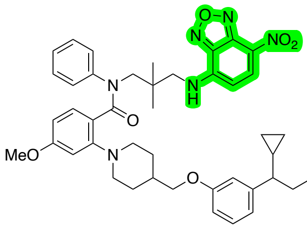 | 5.95±0.16 | 113±14 |
| TUG-2597 | 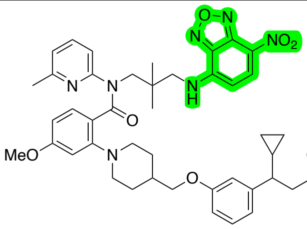 | 7.27±0.15 | 101±4 |
| TUG-2598 | 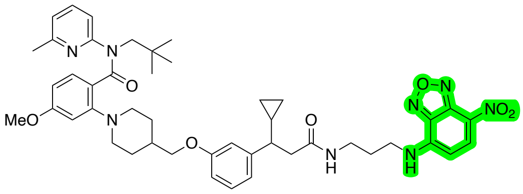 | NR | NR |

NR- No response

*a. $E_{Max}$ is percentage of response to 10 μM T360*

***Supplemental Table 2.** FFA1 Site one Tracer and Precursor $Ca^{2+}$ Data*

| ID | Structure | $pEC_{50}$ | EMax[a] |
|---|---|---|---|
| TUG-770 | 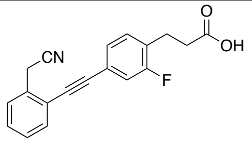 | 7.12±0.09 | 78±7 |
| TUG-905 | 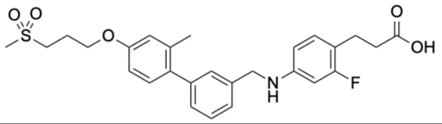 | 6.96±0.14 | 66±7 |
| TUG-1460 | 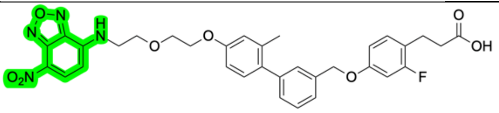 | ND[b] | |
| TUG-2287 | 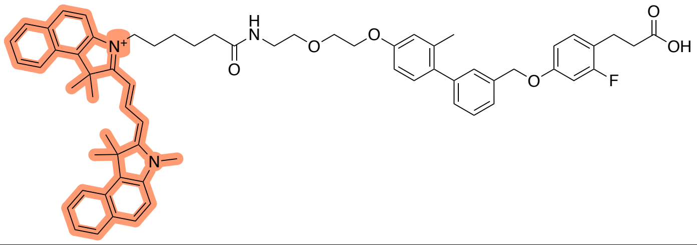 | NR | NR |
| TUG-2355 | 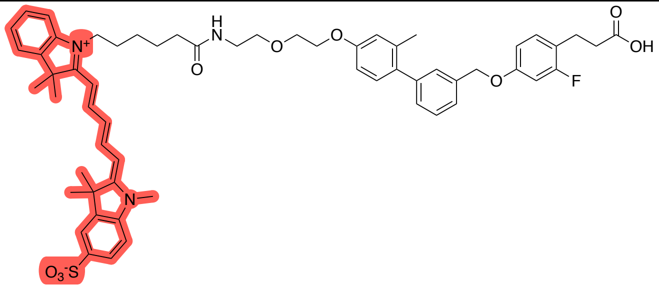 | 5.43±0.06 | 87±10 |
| TUG-2591 | 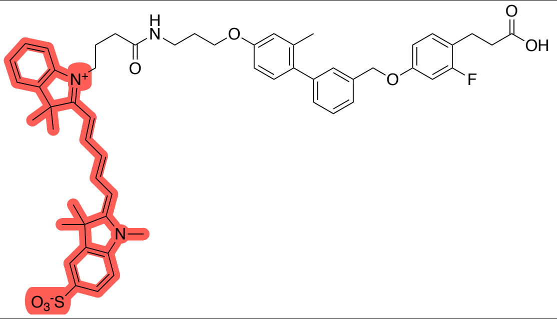 | 6.47±0.16 | 73±9 |

NR- No response
ND- Not determined

a. *$E_{Max}$ is percentage of response to 10 µM T360*
b. *Activity of TUG-1460 is described in [1]*

***Supplemental Table 3.*** *Summary of Competition Binding Data*

| | DISCo-BRET[a] | | | | Nano-BRET[b] | | | | | |
|---|---|---|---|---|---|---|---|---|---|---|
| | SulfoCy5/NBD | | NBD/Nluc | | TUG-2591 | | TUG-2597 | | TUG-2884 | |
| | $pIC_{50}$ | pKi | $pIC_{50}$ | pKi | $pIC_{50}$ | pKi | $pIC_{50}$ | pKi | $pIC_{50}$ | pKi |
| **TUG-905** | 7.6±0.03 (-6.8%) | 8.0±0.03 | 7.8±0.1 (380%) | NA | 7.4±0.05 (-4.2%) | 7.5±0.05 | 8.0±0.05 (220%) | NA | 6.6±0.1 (3.5%) | 7.0±0.1 |
| **TUG-770** | 6.7±0.1 (0.0%) | 7.1±0.1 | 6.9±0.2 (350%) | NA | 6.1±0.02 (0.5%) | 6.3±0.05 | 8.0±0.1 (270%) | NA | ND | ND |
| **TAK-875** | 7.2±0.1 (-3.0%) | 7.6±0.1 | 7.3±0.1 (360%) | NA | 6.7±0.04 (-13%) | 7.0±0.04 | 7.7±0.1 (430%) | NA | ND | ND |
| **Cpd-B** | 6.4±0.1 (5.5%) | 6.8±0.1 | 6.6±0.1 (330%) | NA | 6.0±0.02 (-0.7%) | 6.2±0.02 | 7.6±0.1 (320%) | NA | ND | ND |
| **aLA** | ~3.7 | NA | ~2.9 | NA | ND | ND | ND | ND | ND | ND |
| **T360** | 7.6±0.1 (25%) | NA | 7.1±0.2 (2.7%) | 7.3±0.2 | 8.3±0.1 (150%) | NA | 7.1±0.03 (5.3%) | 7.4±0.03 | NC | NA |
| **Cpd 6h** | 7.4±0.1 (16%) | NA | 7.3±0.1 (9.3%) | 7.5±0.1 | 8.0±0.2 (160%) | NA | 7.2±0.03 (-16%) | 7.5±0.03 | ND | ND |
| **GW1100** | NC | NA | NC | NA | NC | ND | ND | ND | NC | NA |
| **PPTQ** | 6.0±0.1 (8.8%) | 6.4±0.1 | 6.2±0.2 (3.8%) | NA | ND | ND | ND | ND | 5.8±0.06 (-15%) | 6.3±0.06 |
| **TUG-2743** | 6.0±0.1 (3.2%) | 6.5±0.1 | 6.1±0.2 (10%) | NA | ND | ND | ND | ND | ND | ND |
| **TUG-2744** | 6.2±0.07 (26%) | 6.6±0.07 | NC | NA | ND | ND | ND | ND | ND | ND |
| **TUG-2745** | 6.0±0.06 (-15%) | 6.4±0.06 | NC | NA | ND | ND | ND | ND | ND | ND |

NC- No competition
NA- Not applicable
ND- Not determined

*a.* Obtained from DISCo-BRET experiments with 100 nM TUG-2591 and 316 nM TUG-2597. BRET was measured using either SulfoCy5/NBD or NBD/Nluc emission ratios and fit to a one site binding model to obtain $pIC_{50}$ values. The 'bottom' of the $IC_{50}$ curve is shown in parentheses as a percentage of binding obtained in the absence of competing ligand. Binding affinity, pKi values were determined for ligands binding to the same site as the relevant tracer using $K_d$ values for: TUG-2591 (57 nM), obtained from DISCo-BRET saturation experiments; or TUG-2597 (380 nM), obtained from Nano-BRET saturation experiments.

*b.* Obtained from NanoBRET experiments with TUG-2591 (100 nM), TUG-2597 (316 nM), or TUG-2884 (2 μM). BRET was measured using either SulfoCy5/Nluc (TUG-2591) or NBD/Nluc (TUG-2597 and TUG-2884) emission ratios and fit to a one site binding model to obtain $IC_{50}$ values. The 'bottom' of the $IC_{50}$ curve fit is shown in parentheses as a percentage of binding obtained in the absence of competing ligand. Binding affinity, pKi values were determined for ligands binding to the same site as the relevant tracer using the $K_d$ values obtained in NanoBRET saturation experiments: TUG-2591 (200 nM), TUG-2597 (380 nM), and TUG-2884 (940 nM).

***Supplemental Table 4.*** *Ligand Dissociation Rate Data*

| **DISCo-BRET**[a] | Displacing Ligand | | |
|---|---|---|---|
| | TUG-905 | T360 | TUG-905 T360 |
| TUG-2597 / TUG-2591 | $k_{Off}$ (min$^{-1}$) | $k_{Off}$ (min$^{-1}$) | $K_{Off}$ (min$^{-1}$) |
| 100 nM/ 316 nM | | | |
| SulfoCy5/NBD | 0.036 (0.03-0.04) | 0.27 (0.25-0.30) | 0.20 (0.19-0.20) |
| NBD/Nluc | ND | 0.24 (0.19-0.28) | 0.22 (0.19-0.25) |
| 1.0 μM / 3.16 μM | | | |
| SulfoCy5/NBD | 0.032 (0.01-0.06) | 0.18 (0.16-0.19) | 0.21 (0.19-0.22) |
| NBD/Nluc | ND | 0.24 (0.11-0.45) | 0.40 (0.20-0.72) |

**Nano-BRET**

| | $k_{Off}$ (min$^{-1}$) | $k_{On}$ (M$^{-1}$min$^{-1}$) | $K_d$ (nM) |
|---|---|---|---|
| TUG-2591[b] | 0.41 (0.39-0.42) | 1600000 | 250 |
| TUG-2597[c] | 0.38 (0.33-0.43) | 3500000 | 110 |

*a.* DISCo-BRET experiments conducted using FFA1-Nluc and co-adding TUG-2591 (100 nM) and TUG-2597 (316 nM), followed by initiating dissociation by adding TUG-905 (10 μM), T360 (10 μM) or TUG-905 and T360 (10 μM each). Binding was monitored from the same experiment using both the SulfoCy5/NBD and NBD/Nluc BRET ratios. Data presented are the $k_{off}$ in min$^{-1}$ with the 95% CI shown in parentheses.

*b.* Nano-BRET experiment using Nluc-FFA1, adding TUG-2591 (100 nM) followed by TUG-905 (10 μM) to initiate dissociation. Data were fit to an association then dissociation binding model, which was used to define $k_{Off}$, $k_{On}$ and $K_d$. Data for $k_{off}$ in min$^{-1}$ with the 95% CI in parentheses.

*c.* Nano-BRET experiment using Nluc-FFA1, adding TUG-2597 (316 nM) followed by T360 (10 μM) to initiate dissociation. Data were fit to an association then dissociation binding model, which was used to define $k_{Off}$, $k_{On}$ and $K_d$. Data for $k_{off}$ in min$^{-1}$ with the 95% CI in parentheses.

***Supplemental Table 5.*** *Ligand Association Rate Data*

| **TUG-2597** | $k_{On}$ ($M^{-1}min^{-1}$) | $k_{Off}$ ($min^{-1}$) | $K_d$ (nM) |
|---|---|---|---|
| *DISCo-BRET* | | | |
| NBD/Nluc[a] | 1900000<br>(1.5x10$^6$ - 2.2 x10$^6$) | *0.24* | 130 |
| SulfoCy5/NBD[a] | 1700000<br>(1.4x10$^6$ - 2.0 x10$^6$) | *0.24* | 140 |
| *Nano-BRET* | | | |
| Single Concentration[a] | 1400000<br>(1.1x10$^6$ - 1.8 x10$^6$) | *0.38* | 270 |
| Multiple Concentration[b] | 720000<br>(6.2x10$^5$ - 8.3x10$^5$) | 0.39<br>(0.32 - 0.46) | 540 |

| **TUG-2591** | $k_{On}$ ($M^{-1}min^{-1}$) | $k_{Off}$ ($min^{-1}$) | $K_d$ (nM) |
|---|---|---|---|
| *DISCo-BRET* | | | |
| Single Concentration[c] | 1200000<br>(7.3x10$^5$- 1.7x10$^6$) | 0.11 | 91 |
| Multiple Concentration[d] | 500000<br>(4.5x10$^5$ – 5.6x10$^5$) | 0.11<br>(0.09 – 0.13) | 220 |
| *Nano-BRET* | | | |
| Multiple Concentration[e] | 7500000<br>(6.0x10$^6$ – 9.4x10$^6$) | 1.24<br>(1.0 – 1.5) | 170 |

*a.* Values obtained from data presented in **Sup. Fig. 10A**. $k_{Off}$ was based on the value calculated in ***Supplemental Table 4***. 95% CI for calculated $k_{on}$ values are shown in parentheses.

*b.* Values obtained from data presented in **Sup. Fig. 10B**. 95% CI for calculated $k_{on}$ and $k_{off}$ values are shown in parentheses. 95% CI for calculated $k_{on}$ and $k_{off}$ values are shown in parentheses.

*c.* Values obtained from data presented in **Fig. 6C**. $K_{Off}$ was based on the value calculated in the multiple ligand experiment in **Fig. 6H**. 95% CI for calculated $k_{on}$ values are shown in parentheses.

*d.* Values obtained from data presented in **Fig. 6H**. 95% CI for calculated $k_{on}$ and $k_{off}$ values are shown in parentheses.

*e.* Values obtained from data presented in **Fig. 6I**. 95% CI for calculated $k_{on}$ and $k_{off}$ values are shown in parentheses.

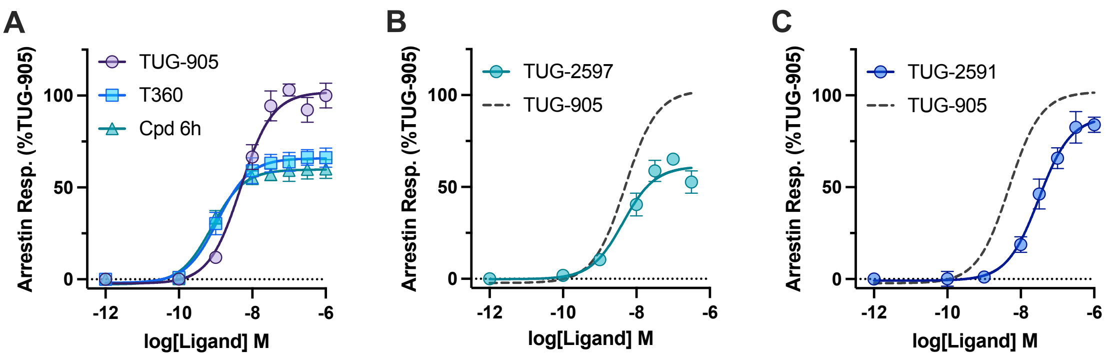


D

| Compound | $pEC_{50}$ | $E_{Max}$ (% TUG-905) |
|---|---|---|
| TUG-905 | 8.3 ± 0.2 | 103 ± 2% |
| T360 | 8.9 ± 0.4 | 66 ± 8% |
| Cpd 6h | 9.1 ± 0.1 | 54 ± 6% |
| TUG-2597 | 7.5 ± 0.4 | 60 ± 6% |
| TUG-2591 | 8.5 ± 0.3 | 91 ± 6% |

***Supplemental Figure 1***. *FFA1 allosteric site two ligands are partial agonist for arrestin recruitment*. Recruitment of arrestin-3 to the cell membrane was assessed in HEK293T cells transfected with FFA1 and with a bystander NanoBiT-based arrestin biosensor. **A**. Arrestin-3 recruitment in response to FFA1 site one ligand, TUG-905, or site two ligands, T360 and Cpd 6h are shown as a percentage of the response to 10 μM TUG-905. Both site two agonists appear to be partial agonists relative to the site one ligand. Arrestin recruitment in response to the green tracer TUG-2597 (**B**), shows partial agonism consistent with site two interaction, while recruitment in response to the red tracer, TUG-2591 (**C**) shows near full agonism, consistent with site one interaction. The TUG-905 concentration response curve from **A** is shown in **B** and **C** for reference (dashed line). **D**. Curve fit values for $pEC_{50}$ ± SEM and $E_{Max}$ ± SEM for the data presented in **A-C**. Data are from N=3 independent experiments.

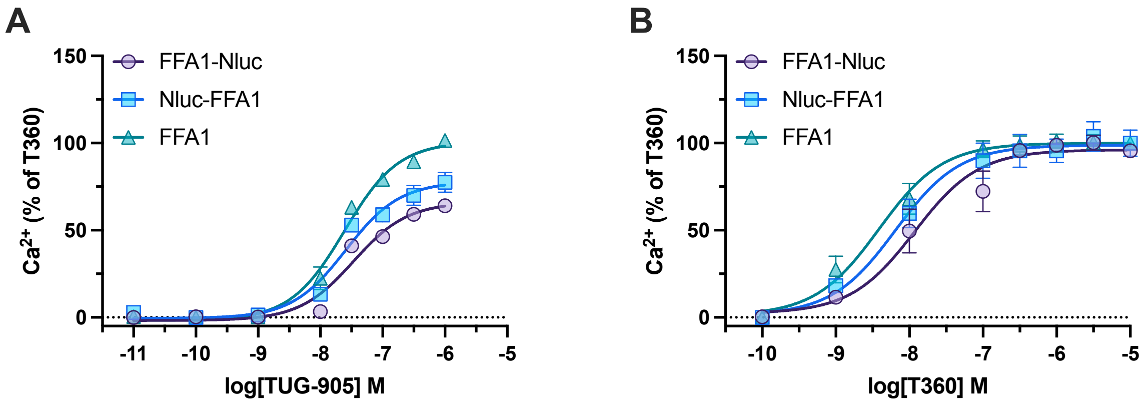


***Supplemental Figure 2.*** *Modification of FFA1 with Nluc at either the N or C terminal has little effect on receptor pharmacology.* $Ca^{2+}$ mobilization responses to increasing concentrations of TUG-905 (**A**) or T360 (**B**) were measured in Flp-In T-REx 293 cells engineered to express FFA1 constructs either tagged at the C (FFA1-Nluc) or N (Nluc-FFA1) terminal with Nluc, or completely untagged (FFA1). Data show a modest increase in efficacy for the untagged receptor for the partial agonist, TUG-905, and a modest increase in potency for the full agonist, T360. These results may suggest slightly higher expression of the untagged receptor, but similar ligand pharmacology across all three FFA1 constructs. Data are presented as a percentage of the $Ca^{2+}$ mobilization response obtained to 10 µM T360 at each FFA1 construct. Data are from N=4 independent experiments.

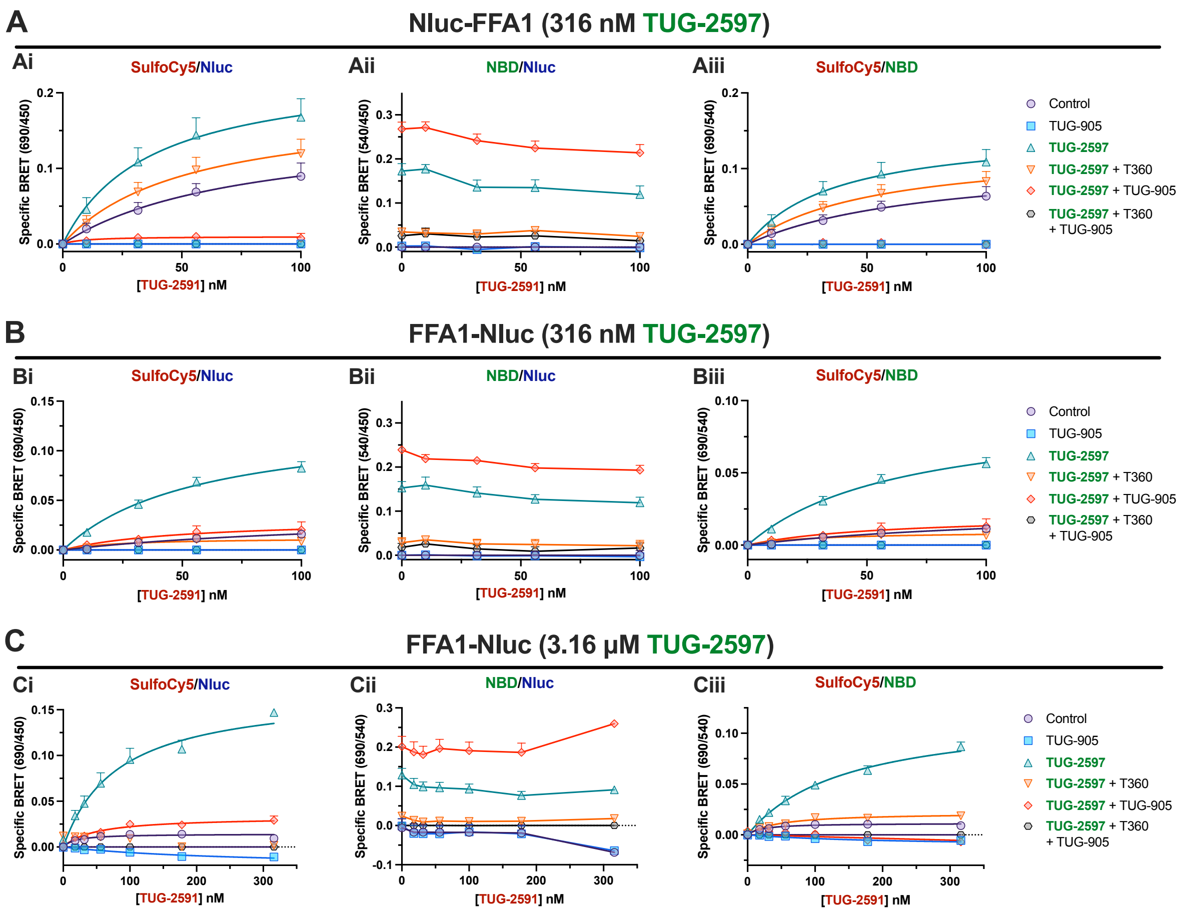


***Supplemental Figure 3.*** *DISCo-BRET can be measured using a C terminal Nluc tagged FFA1 construct with simple BRET ratios*. Saturation binding experiments for the red site one FFA1 tracer, TUG-2591, were conducted using an N (**A**) or C (**B**) terminal Nluc tagged FFA1 construct, either without any additional compounds (control), or in the presence of a competing ligand, TUG-905 (10 μM); the green tracer, TUG-2597 (316 nM); the competing ligand for the green tracer, T360 (10 μM); or combinations of these three ligands. Luminesces was monitored at three wavelengths: 450 nm for Nluc emission, 540 nm for NBD emission, and 690 nm for SulfoCy5 emission. The data were then expressed as a simple BRET ratio for SulfoCy5/Nluc wavelength emission (**i**), NBD/Nluc wavelength emission (**ii**), or SulfoCy5/NBD wavelength emission (**iii**). Data are from N= 4 independent experiments and yielded $K_d$ values for TUG-2591 of 57 nM (**Bi**) and 64 nM (**Biii**). **C.** Comparable experiments carried out in cells expressing the FFA1-Nluc construct, but using a 3.16 μM concentration of the green tracer, TUG-2597, and 30 μM concentrations of the competing ligands (TUG-905 and T360). Data are from N=4 independent experiments and yielded $K_d$ values for TUG-2591 of 81 nM (**Ci**) and 140 nM (**Ciii**).

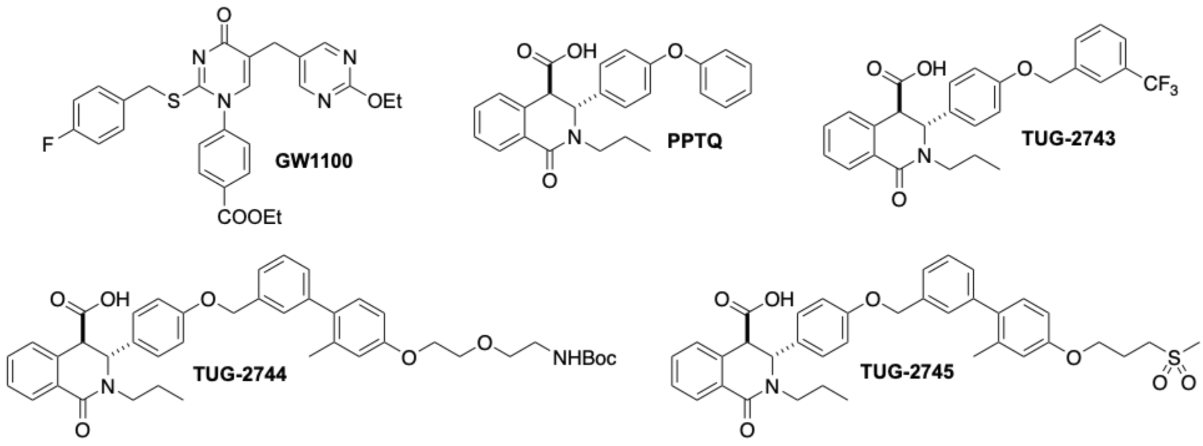


***Supplemental Figure 4.*** *Chemical Structures of FFA1 antagonists.*

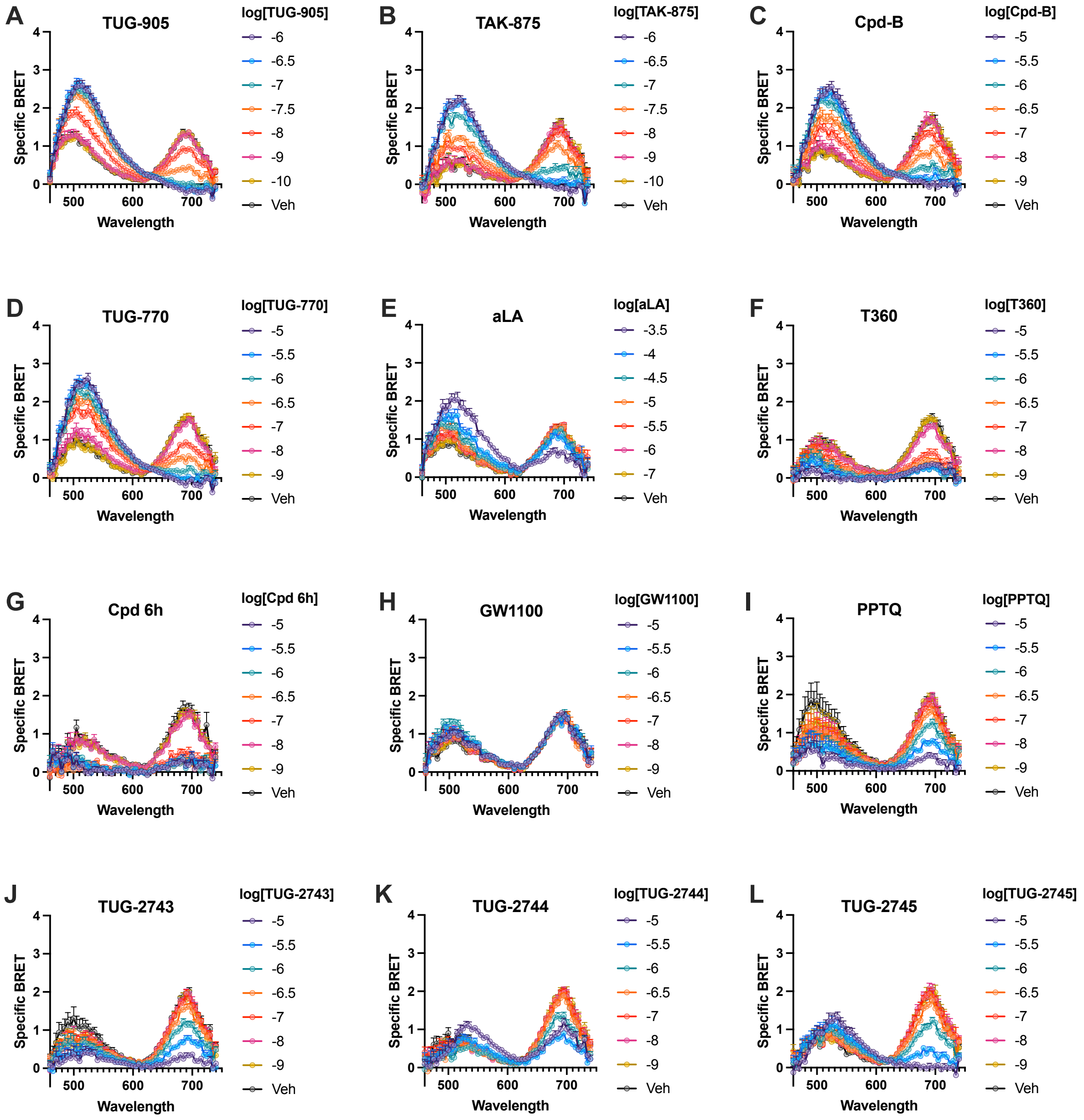


***Supplemental Figure 5.*** *BRET Spectral analysis reveals the effects of allosteric site one and site two competition on DISCo-BRET.* BRET spectra were obtained in the presence of both the red allosteric site one tracer, TUG-2591 (100 nM) and the allosteric site two green tracer, TUG-2597 (316 nM) with increasing concentrations of: TUG-905 (**A**), TAK-875 (**B**), Cpd-B (**C**), TUG-770 (**D**), α-linolenic acid (aLA) (**E**), T360 (**F**), Cpd 6h (**G**), GW1100 (**H**), PPTQ (**I**), TUG-2743 (**J**), TUG-2744 (**K**), and TUG-2745 (**L**). Specific BRET spectra were obtained by subtracting the luminescence spectrum obtained from the non-specific conditions that included TUG-2591 and TUG-2597 as well as their two respective competing ligands, TUG-905 (10 μM) and T360 (10 μM). Data are presented from N=3-7 independent experiments.

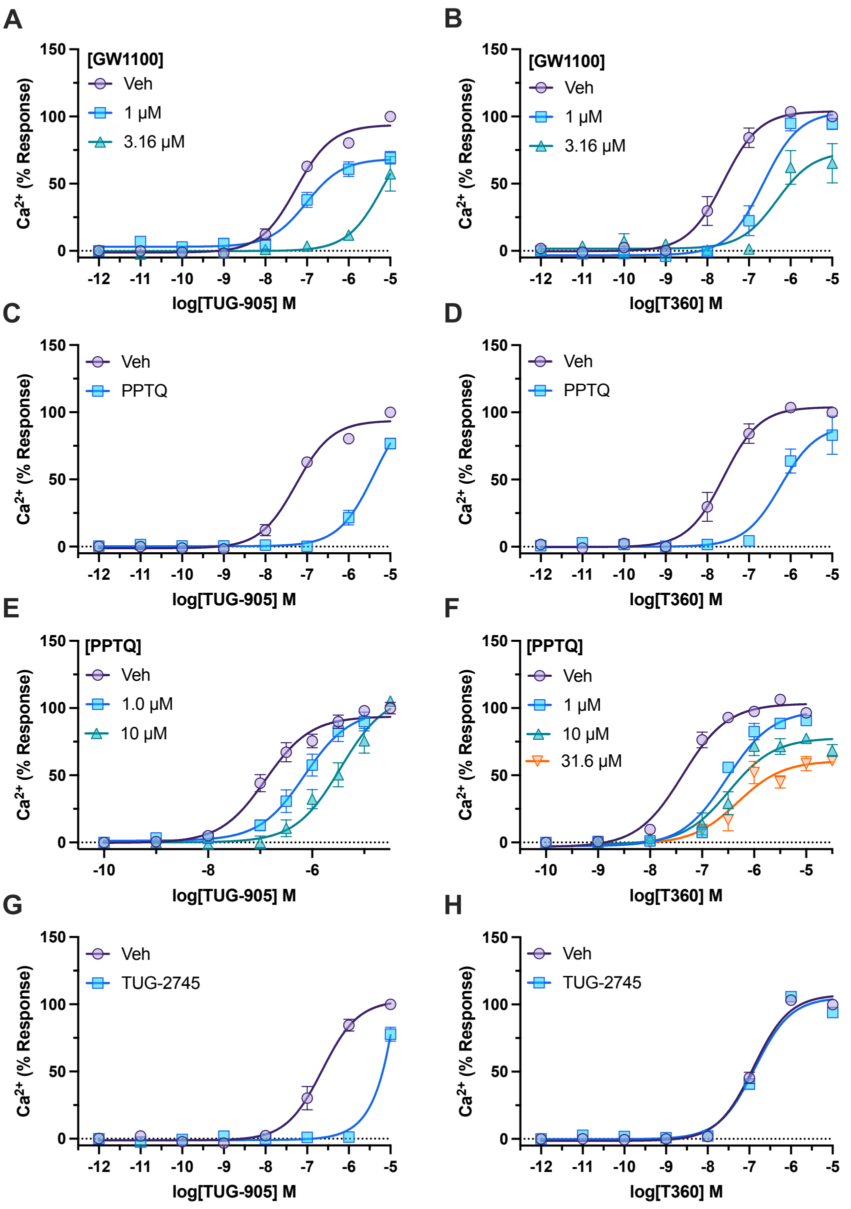


***Supplemental Figure 6.*** *FFA1 antagonists exhibit different modes of action when inhibiting site one vs site two agonists.* $Ca^{2+}$ mobilization responses were measured in Flp-In T-REx 293 cells expressing FFA1-Nluc. $Ca^{2+}$ responses to increasing concentrations of FFA1 agonists TUG-901 (**A**) or T360 (**B**) following pre-treated with vehicle (0.1% DMSO), 1.0 or 3.16 μM GW1100, demonstrate that GW1100 inhibits signalling by each agonist. Similar experiments show inhibition of responses to TUG-905 (**C**) and T360 (**D**) in cells pre-treated with 10 μM PPTQ. The mode of antagonism for PPTQ was assessed by generating concentration responses to TUG-905 (**E**) or T360 (**F**), following pre-treatment with the indicated concentrations of PPTQ or vehicle (0.316% DMSO). The results are consistent with competitive antagonism of TUG-905 and allosteric modulation of T360. $Ca^{2+}$ responses to TUG-905 (**G**) or T360 (**H**) are shown from cells pre-treated with TUG-2745 (10 μM) or vehicle (0.1% DMSO). This compound inhibits signalling by the site one agonist, TUG-905, but has no effect on the site two agonist, T360. Data presented are from N=3-4 independent experiments.

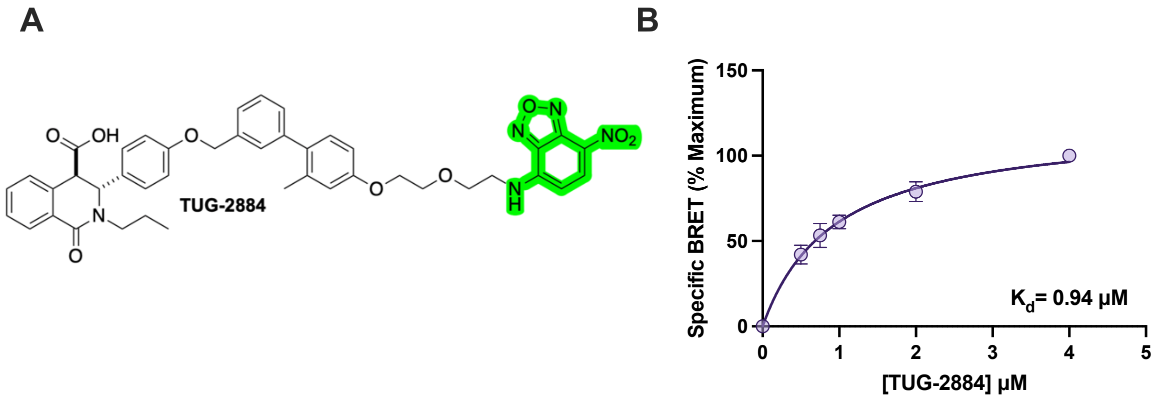


***Supplemental Figure 7****. TUG-2884 is an FFA1 antagonist fluorescent tracer.* **A**. The chemical structure of TUG-2884 with the location of the NBD green fluorophore indicated. **B**. Nano-BRET saturation binding assays in Flp-In T-REx cells expressing Nluc-FFA1. Data show normalized specific binding BRET obtained after subtracting non-specific binding (obtained in the presence of 10 µM TUG-905). Data are fit to a single site specific binding saturation model, yielding a $K_d$ value of 0.94 µM for TUG-2884. Data are presented from N=3 independent experiments.

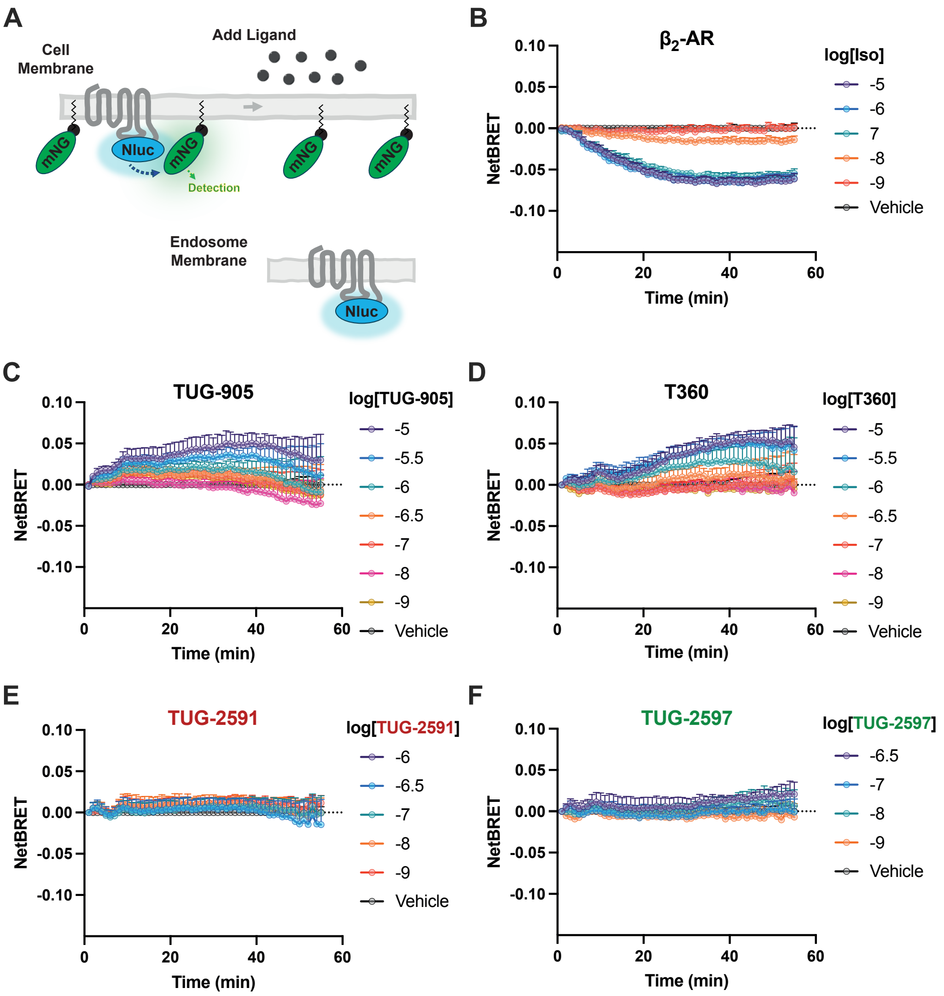


***Supplemental Figure 8.*** *FFA1 is not internalized following treatment with either site one or site two agonists.* **A.** A bystander BRET internalization assay employed an mNG protein anchored to the cell membrane with the CAAX motif and a receptor tagged at its C terminal with Nluc. In the absence of agonist treatment, there is high bystander BRET while internalization of the receptor leads to a reduction in BRET. **B.** Bystander BRET internalization assay in cells transfected with $\beta_2$-Adrenoceptor-Nluc and treated with increasing concentrations of $\beta_2$-Adrenoceptor agonist, isoprenaline. BRET ratios are shown as the NetBRET response after subtracting the BRET ratio obtained from vehicle treated cells. The data show a clear decrease in BRET, representative of internalisation of the $\beta_2$-Adrenoceptor-Nluc construct. Comparable experiments were conducted in cells expressing FFA1-Nluc and treated with increasing concentrations of TUG-905 (**C**), T360 (**D**), TUG-2591 (**E**) and TUG-2597 (**F**). No internalization was observed following treatment with any FFA1 ligands, and TUG-905 and T360 both appeared to produce modest increases in BRET, suggesting these ligands increase FFA1 expression at the cell surface. The maximum concentrations used for TUG-2591 (1 μM) and TUG-2597 (0.316 μM), were chosen based on concentrations used in kinetic binding assays, and because higher concentrations of these tracer ligands resulted in non-specific interference in the BRET internalization assay. Data are presented from N=4 independent experiments.

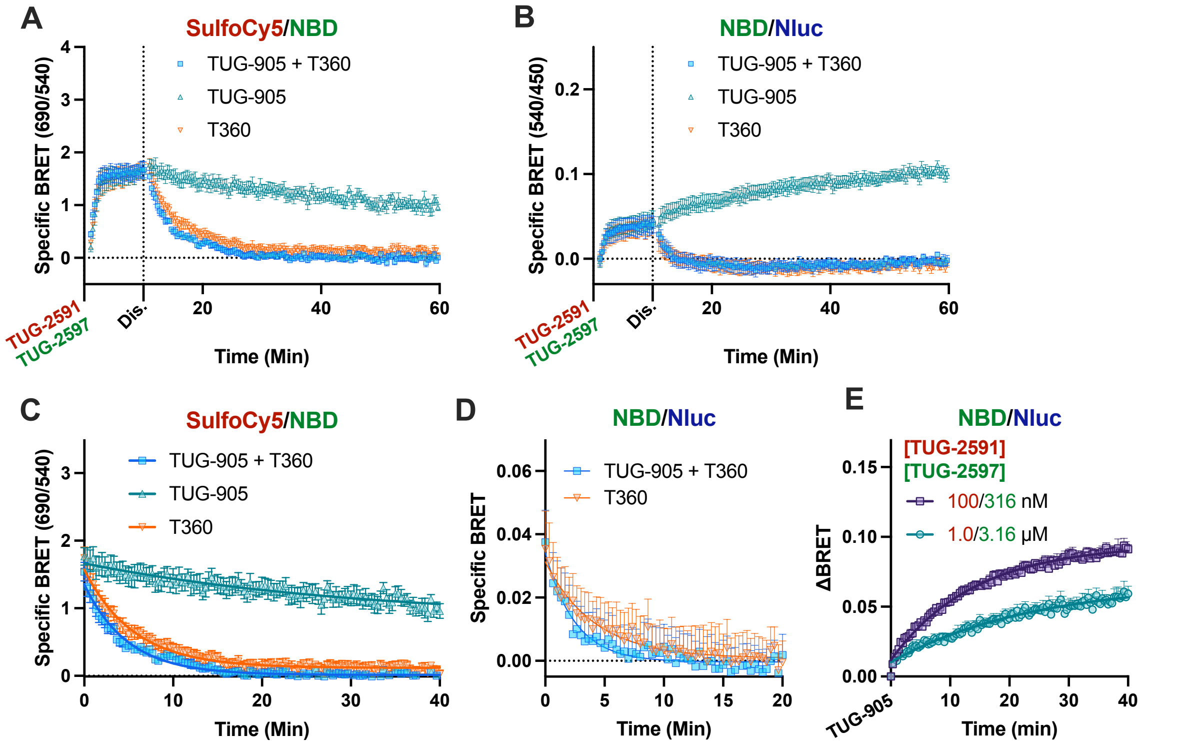


***Supplemental Figure 9.*** *DISCo-BRET off-rate kinetics with increased concentrations of tracers.* DISCo-BRET kinetic binding experiments with cells treated with TUG-2591 (1.0 μM) and TUG-2597 (3.16 μM) together, before the addition (marked as Dis.) of TUG-905 (30 μM), T360 (30 μM) or both TUG-905 and T360 (30 μM each). Data are shown as the 690/540 nm (**A**), or 540/450 nm (**B**) emission ratios. **C**. The dissociation from **A** is fit to a one site exponential decay model. **D**. The dissociation portion from **B** is fit to a one site exponential decay model. **E**. A comparison of the increase in 540 nm / 450 nm ratio following addition of TUG-905 in cells pre-treated with either 100/316 nM (from **Figure 5C**) or 1.0/3.16 μM (from panel **B** of this figure) concentrations of TUG-2591/TUG-2597. Data are presented as the ΔBRET by subtracting the BRET ratio immediately before TUG-905 addition. Data are fit to a one phase association model, yielding half-times of 9.8 min (100/316 nM) or 16.6 min (1.0/3.16 μM). Data are from N=4 independent experiments.

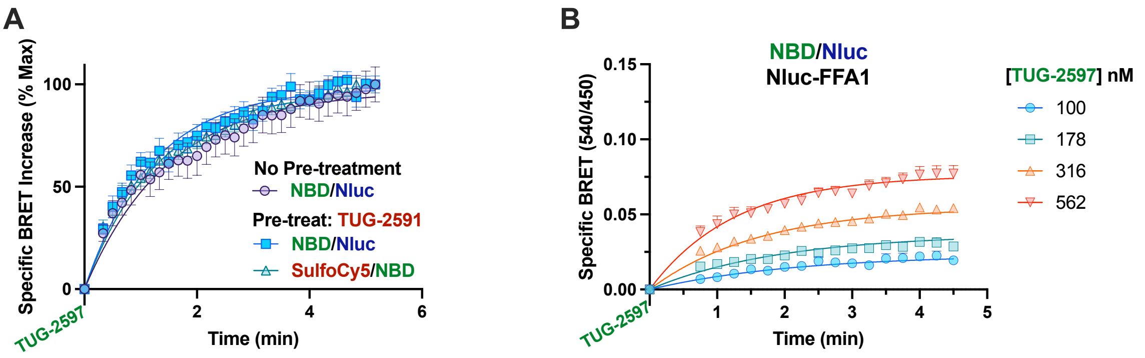


***Supplemental Figure 10.*** *TUG-2597 Association binding kinetics is largely unaffected by TUG-2591 binding*. Analysis of DISCo-BRET data shown in **Figure 6B**, **6E** and **6F** are shown. **A**. DISCo-BRET association experiments follow the change in specific BRET after treatment with TUG-2597, either in the absence or presence of TUG-2591 pre-treatment. Following TUG-2591 pre-treatment, curves are shown tracking both BRET between TUG-2597 and Nluc (NBD/Nluc: 540 nm / 450 nm) and between TUG-2591 and TUG-2597 (SulfoCy5/NBD: 690 nm / 540 nm). **B**. Nano-BRET competition data using N terminal Nluc tagged FFA1 and increasing concentrations of TUG-2597. Specific BRET was determined by subtracting the BRET obtained in the presence of T360 (10 μM). Data are from N=4 independent experiments and are fit to a multiple concentration of labelled ligand binding association model.

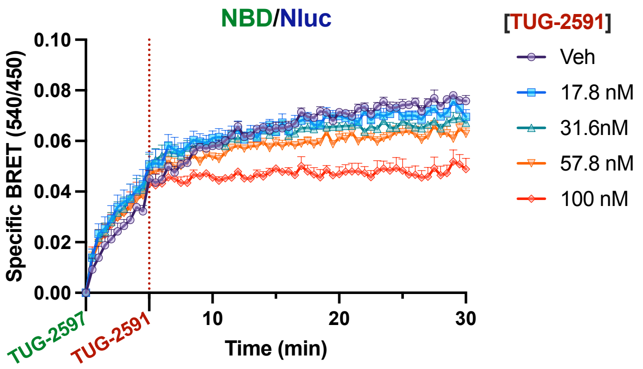


***Supplemental Figure 11.*** *Addition of increasing concentrations of TUG-2591 has a modest effect on BRET between TUG-2597 and Nluc*. Data shown are the NBD/Nluc (540 nm / 450 nm) emission ratios obtained from the experiment presented in **Figure 6G**. Cells were treated first with TUG-2597 (316 nM), followed by addition of TUG-2591 (100 nM). Specific BRET was obtained by subtracting BRET obtained in the presence of both TUG-905 and T360 (10 µM each). Data are shown from N=4 independent experiments.

***Supplemental Chemistry Methods***

**General experimental information**

All commercially available starting materials were used without further purification unless otherwise stated. Anhydrous reactions were carried out in flame-dried glassware under argon or nitrogen atmosphere. Inert atmosphere inside the reaction vessel was generally obtained by capping the flask with a septum, that was pierced with a needle attached to a Schlenk line connected to the *in-house* vacuum. DCM, THF and DMF were dried using a Glass Contour Solvent System built by SG Water USA. *N,N*-Diisopropylethylamine (DIPEA), triethylamine ($Et_3N$), MeCN and 1,4-dioxane were stored over 3 Å sieves. Water used in reactions was demineralized and water used for freeze-drying was filtered Milli-Q. Thin layer chromatography (TLC) was performed on TLC pre-coated silica gel 60 F254 plates (Merck), and visualized under UV light (254 or 365 nm). Purification of compounds was performed using silica gel 60 (0.040-0.063 mm, Merck); automated flash column chromatography was performed on a Reveleris® X2 Flash Chromatography System (Büchi); preparative HPLC on a Dionex Ultimate HPLC system (Thermo Scientific), using a Gemini-NX C18 column (21.2 × 250 mm × 5 μm, 110 Å) and a flow of 20 mL/min, unless otherwise stated. Mobile phase A: $H_2O$:TFA, 100:0.1, v/v. Mobile phase B: MeCN:$H_2O$:TFA, 90:10:0.1, v/v/v. Data were acquired and processed using the Chromeleon Software v. 6.80. Nuclear Magnetic Resonance (NMR) spectra were recorded on 400 or 600 MHz Bruker instruments ($^1H$ NMRs were obtained at 400 or 600 MHz, $^{13}C$ NMRs were obtained at 100 or 151 MHz and $^{19}F$ NMR at 376 MHz) at 300 K. Signals are reported in ppm (δ) using the solvent as reference except for $^{19}F$ NMR which were recorded without internal standard and used primarily for purity analysis. Mass spectrometry (MS) was performed on an Aquity UPLC instrument or on an Agilent 6130 Mass Spectrometer instrument using electron spray ionization (ESI). High-resolution mass spectra (HRMS) were recorded on a QExactive Orbitrap mass spectrometer equipped with a SMALDI5 ion source. The sample was analysed in the positive ion mode using a peak from the DHB matrix for internal mass calibration whereby a mass accuracy of 2 ppm or better was achieved. Analytical High Performance Liquid Chromatography (HPLC) was performed on a Dionex UltiMate HPLC system (Thermo Scientific) using a Gemini-NX C18 column (4.6 × 250 mm, 3 μm, 110 Å);

Mobile phase A: $H_2O$:TFA, 100:0.1, v/v. Mobile phase B: MeCN: $H_2O$:TFA, 90:10:0.1, v/v/v; UV detection at 254 nm, unless otherwise stated. Data were acquired and processed using the Chromeleon Software v. 6.80. All test compounds were of ≥95% purity.

**Synthetic procedures**

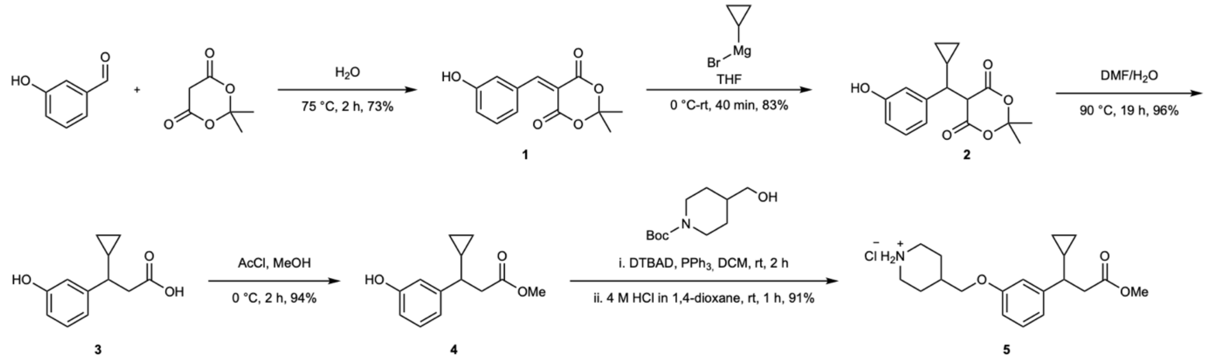


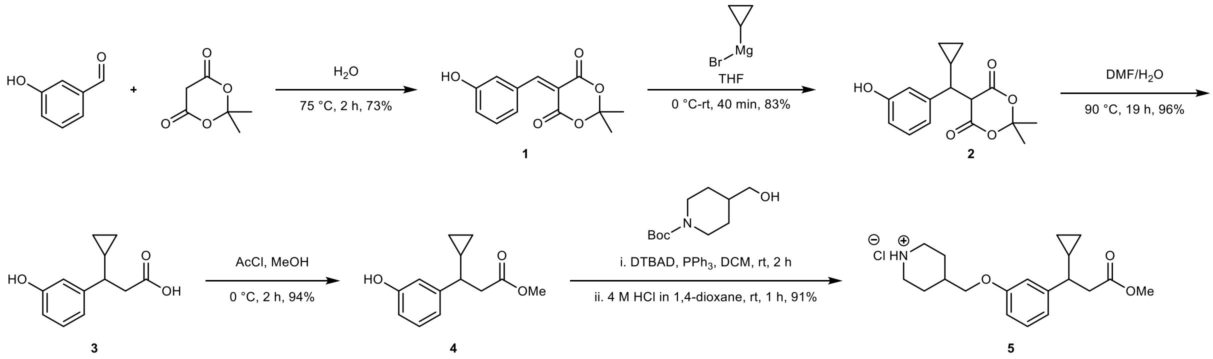


**5-(3-Hydroxybenzylidene)-2,2-dimethyl-1,3-dioxane-4,6-dione (1)**

Compound synthesized according to Brown *et al.*[2] but using 3-hydroxy benzaldehyde. A flask was charged with 3-hydroxy benzaldehyde (2.50 g, 20.5 mmol) and $H_2O$ (23 mL), and the resulting suspension was heated in an oil-bath to 75 °C until it became a solution. Upon solubilization, a slurry of Meldrum's acid (3.80 g, 26.4 mmol) in $H_2O$ (21 mL) was added portion-wise, and the reaction turned into a yellow suspension that was stirred at 75 °C. After 2 h the reaction was stopped, cooled to rt and the yellow solid was filtered by vacuum filtration. The solid was washed with water and freeze-dried to obtained 3.74 g (73% crude) of the product as a bright yellow solid that was used in the next step without further purification: **$^1$H NMR** (600 MHz, DMSO-$d_6$) δ 9.79 (s, 1H), 8.25 (s, 1H), 7.47 (t, $J$ = 2.1 Hz, 1H), 7.42 – 7.37 (m, 1H), 7.31 (t, $J$ = 7.9 Hz, 1H), 7.00 (ddd, $J$ = 8.2, 2.5, 0.9 Hz, 1H), 1.76 (s, 6H); **$^{13}$C NMR** (151 MHz, DMSO-$d_6$) δ 162.6, 159.4, 157.1,

156.6, 132.9, 129.5, 124.5, 120.4, 118.6, 115.5, 104.5, 27.0. Spectra in agreement with literature.[3]

**Methyl 3-cyclopropyl-3-(3-hydroxyphenyl)propanoate (4)**

A vial charged with MeOH (1 mL) was cooled to 0 °C and acetyl chloride (643 µL, 9.04 mmol) was added dropwise. The mixture was stirred at 0 °C for 10 min. Then, a solution of compound **3**, synthesized according to Brown *et al.*[3] (373 mg, 1.81 mmol) in MeOH (2.80 mL) was added dropwise, and the stirring was continued at 0 °C for 2 h. Upon completion monitored by TLC, the reaction mixture was concentrated *in vacuo*, diluted with EtOAc and washed with sat. aq. $NaHCO_3$ (×3). The aqueous phase was extracted with EtOAc (×2). The organic layers were combined, dried over $MgSO_4$, filtered, and concentrated *in vacuo* to provide 375 mg (94% crude) of a brown oil, that was used in the next step without further purification. Spectra in agreement with literature.[3]

**Methyl 3-cyclopropyl-3-(3-(piperidin-4-ylmethoxy)phenyl)propanoate hydrochloride (5)**

Compound **4** (300 mg, 1.36 mmol), $PPh_3$ (357 mg, 1.36 mmol) and *N*-boc 4-hydroxymethyl piperidine (147 mg, 0.68 mmol) were charged in a flask and dissolved in anh. DCM (14 mL), under Ar. The flask was evacuated and backfilled with Ar (×3), di-*tert*-butyl azodicarboxylate (DTBAD) (314 mg, 1.36 mmol) was added and the reaction was stirred at rt. After 2 h, the reaction was monitored by TLC (EtOAc:Petroleum ether, 1:1), and the consumption of the limiting reagent was observed. Then, 4 M HCl in 1,4-dioxane (5 mL) was added dropwise and the reaction was stirred at rt for 1 h. Upon completion, the reaction mixture was concentrated *in vacuo*. The crude product was purified by flash column chromatography (DCM:MeOH, 9:1 → 4:1) to afford 219 mg (91%) of a white solid as HCl salt: **$^1$H NMR** (400 MHz, $CDCl_3$) δ 9.72 (br s, 1H), 9.43 (br s, 1H), 7.20 (t, *J* = 7.9 Hz, 1H), 6.83 (d, *J* = 7.9 Hz, 1H), 6.77 – 6.69 (m, 2H), 3.84 (d, *J* = 5.8 Hz, 2H), 3.63 – 3.52 (m, 5H), 3.13 – 2.86 (m, 2H), 2.82 – 2.66 (m, 2H), 2.40 – 2.28 (m, 1H), 2.17 – 2.03 (m, 3H), 1.94 – 1.75 (m, 2H), 1.07 – 0.94 (m, 1H), 0.63 – 0.52 (m, 1H), 0.51 – 0.37 (m, 1H), 0.31 – 0.20 (m, 1H), 0.20 – 0.04 (m, 1H); **$^{13}$C NMR** (101 MHz, $CDCl_3$) δ 172.9, 158.7, 146.1, 129.5, 120.2, 113.9, 112.2, 71.2, 51.6, 47.3, 43.8, 41.6, 34.6, 25.9, 17.2, 5.4, 4.2; **ESI-MS** m/z 318.4 ($M+H^+$).

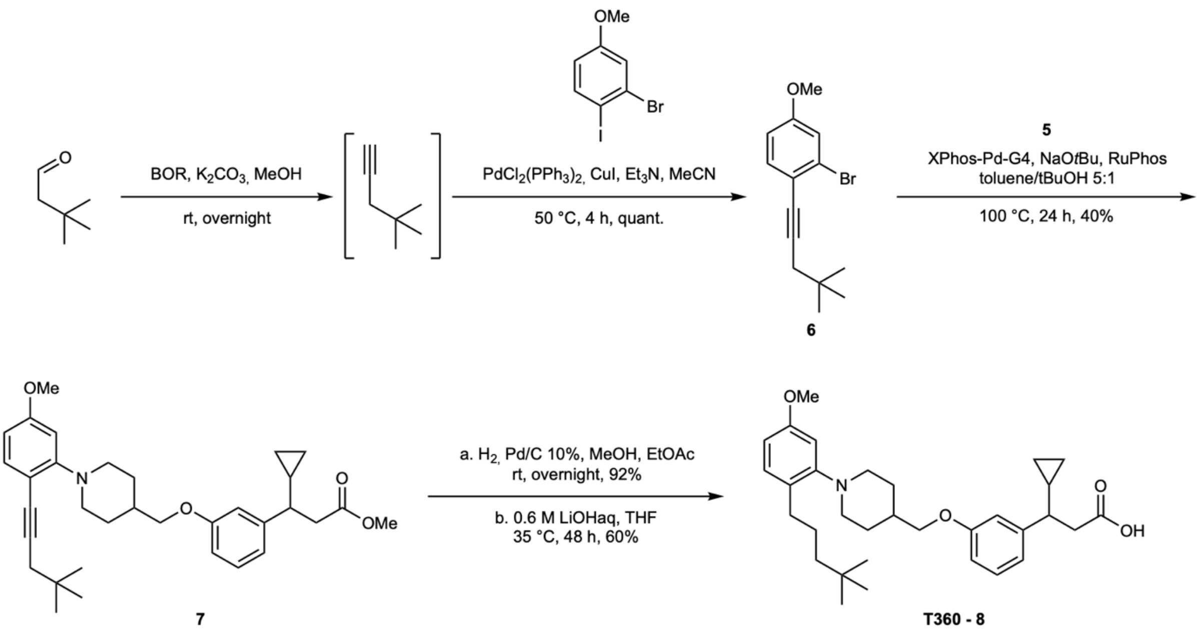

**2-Bromo-1-(4,4-dimethylpent-1-yn-1-yl)-4-methoxybenzene (6)**

Compound synthesized according to Capitta *et al.*[4] but using 3,3-dimethylbutyraldehyde and 4-iodo anisole. A microwave vial under Ar was charged with $K_2CO_3$ (138 mg, 1.00 mmol), capped with a septum, and a solution of 3,3-dimethylbutyraldehyde (63 µL, 0.50 mmol) in MeOH (200 µL) was added. Then, a solution of dimethyl-1-diazo-2-oxopropylphosphonate (90 µL, 0.60 mmol) in MeOH (200 µL) was added, and upon addition, the reaction mixture bubbled and warmed up for few minutes. The vial was capped, and the reaction was stirred at rt overnight. To the resulting yellow suspension, $Pd(PPh_3)_2Cl_2$ (11 mg, 3 mol%) and CuI (5 mg, 5 mol%) were quickly added under Ar, followed by dropwise addition of a solution of $Et_3N$ (70 µL, 0.50 mmol) and 3-bromo 4-iodo anisole (50 µL, 0.33 mmol) in anh. MeCN (2.8 mL). The reaction mixture was stirred at 50 °C for 4 h. Upon completion, the reaction was cooled to rt, concentrated *in vacuo* and the crude mixture was directly purified by flash column chromatography (Petroleum ether) to afford 116 mg (quant.) of a yellow oil: $R_f$ = 0.33 (Petroleum ether); **$^1$H NMR** (400 MHz, $CDCl_3$) δ 7.35 (d, *J* = 8.6 Hz, 1H), 7.10 (d, *J* = 2.5 Hz, 1H), 6.78 (dd, *J* = 8.7, 2.6 Hz, 1H), 3.79 (s, 3H), 2.32 (s, 2H), 1.08 (s, 9H); **$^{13}$C NMR** (101 MHz, $CDCl_3$) δ 159.3, 134.0, 126.2, 118.6, 117.7, 113.6, 92.1, 80.6, 55.7, 34.8, 31.7, 29.3.

**Methyl-3-cyclopropyl-3-(3-((1-(2-(4,4-dimethylpent-1-yn-1-yl)-5-methoxyphenyl)-piperidin-4-yl)methoxy)phenyl)propanoate (7)**

To a Schlenk under Ar, RuPhos (1 mg, 5 mol%), XPhos-Pd-G4 (2 mg, 5 mol%), NaO*t*-Bu (9 mg, 0.09 mmol), compound **5** (15 mg, 0.04 mmol) and compound **6** (12 mg, 0.04 mmol) were added. The Schlenk flask was evacuated and backfilled with Ar (×3) before adding a degassed mixture of toluene/*t*-BuOH (5:1, 0.30 mL). The reaction was stirred at 100 °C for 24 h. Then, the reaction mixture was cooled to rt, concentrated *in vacuo* and directly purified by flash column chromatography (EtOAc:*n*-heptane, 5:95 → 10:90) to give 7 mg (40%) of a pale yellow oil: $R_f$ = 0.26 (EtOAc:Petroleum ether, 1:9); **$^1$H NMR** (400 MHz, $CDCl_3$) δ 7.32 (d, *J* = 8.4 Hz, 1H), 7.22 (t, *J* = 7.8 Hz, 1H), 6.87 – 6.72 (m, 3H), 6.55 – 6.34 (m, 2H), 3.85 (d, *J* = 6.0 Hz, 2H), 3.82 – 3.69 (m, 5H), 3.62 (s, 3H), 2.86 – 2.61 (m, 4H), 2.43 – 2.26 (m, 3H), 1.98 – 1.91 (m, 3H), 1.67 – 1.57 (m, 2H), 1.09 – 0.99 (m, 10H), 0.63 – 0.52 (m, 1H), 0.51 – 0.38 (m, 1H), 0.32 – 0.21 (m, 1H), 0.21 – 0.11 (m, 1H); **$^{13}$C NMR** (151 MHz, $CDCl_3$) δ 173.0, 160.1, 159.3, 156.2, 146.0, 135.2, 129.5, 119.7, 113.9, 112.3, 110.5, 106.0, 104.9, 92.4, 80.5, 72.7, 55.4, 51.7, 51.6, 47.4, 41.7, 36.2, 35.1, 31.6, 29.7, 29.3, 17.2, 5.4, 4.2; **ESI-MS** m/z 518.5 ($M+H^+$).

**4-((3-(2-Carboxy-1-cyclopropylethyl)phenoxy)methyl)-1-(2-(4,4-dimethylpentyl)-5-methoxyphenyl)piperidin-1-ium 2,2,2-trifluoroacetate (T360 - 8)**

**Step 1:** To a solution of compound **7** (17 mg, 0.03 mmol) in a mixture of MeOH (107 µL) and EtOAc (50 µL) under Ar, Pd/C 10% (2 mg, 12 wt %) was added. The flask was evacuated and backfilled with Ar (×3) and then with $H_2$ (×3) and the reaction was stirred at rt overnight under $H_2$ atmosphere. Upon completion, the crude mixture was filtered through a Celite® pad, that washed with EtOAc, and the filtrate was concentrated *in vacuo* to give 15 mg (92% crude) of a yellow oil that was used in the next step without further purification: $R_f$ = 0.39 (EtOAc:Petroleum ether, 1:9); **$^1$H NMR** (400 MHz, $CDCl_3$) δ 7.22 (t, *J* = 7.8 Hz, 1H), 7.11 (d, *J* = 8.4 Hz, 1H), 6.87 – 6.74 (m, 3H), 6.66 (d, *J* = 2.6 Hz, 1H), 6.58 (dd, *J* = 8.3, 2.6 Hz, 1H), 3.86 (d, *J* = 5.8 Hz, 2H), 3.79 (s, 3H), 3.62 (s, 3H), 3.16 – 3.07 (m, 2H), 2.82 – 2.58 (m, 4H), 2.58 – 2.48 (m, 2H), 2.40 – 2.32 (m, 1H), 1.98 – 1.89 (m, 3H), 1.65 – 1.45 (m, 4H), 1.32 – 1.19 (m, 2H), 1.10 – 0.94 (m, 1H), 0.88 (s, 9H), 0.64 – 0.51 (m, 1H), 0.49 – 0.38 (m, 1H), 0.32 – 0.22 (m, 1H), 0.22 – 0.11 (m, 1H); **$^{13}$C NMR** (101 MHz, $CDCl_3$) δ 173.0, 159.3, 158.4, 153.6, 146.0, 130.6, 130.1, 129.5, 119.7, 114.0, 112.3,

108.0, 107.1, 72.8, 55.4, 53.2, 51.6, 47.4, 44.7, 41.7, 36.2, 31.4, 30.5, 30.1, 29.6, 26.1, 17.2, 5.4, 4.2; **ESI-MS** m/z 522.7 (M+H$^+$).

**Step 2:** Compound from Step 1 (15 mg, 0.03 mmol) was dissolved in THF (326 µL), 0.6 M $LiOH_{aq}$ (550 µL, 0.33 mmol) was added, and the reaction was stirred at 35 °C for 48 h. Upon completion, the reaction was diluted with $H_2O$, acidified with 1 M $HCl_{aq}$ and extracted with EtOAc (×3). The organic layers were combined, washed with brine, dried over $Na_2SO_4$ and filtered. The residue was concentrated *in vacuo*, purified by preparative HPLC (50 to 100% B over 10 min) and the combined fractions were lyophilized to give 11 mg (60%) of the title compound as a colourless oily TFA salt: $t_R$ = 7.6 min (purity 97.03% by HPLC at 215 nm); $^1$**H NMR** (400 MHz, $CD_3OD$) δ 7.36 (d, *J* = 8.6 Hz, 1H), 7.21 (t, *J* = 7.9 Hz, 1H), 7.14 – 7.11 (m, 1H), 6.99 (dd, *J* = 8.6, 2.5 Hz, 1H), 6.89 – 6.83 (m, 2H), 6.82 – 6.76 (m, 1H), 3.97 (d, *J* = 5.8 Hz, 2H), 3.84 (s, 3H), 3.61 – 3.50 (m, 4H), 2.82 – 2.62 (m, 4H), 2.37 – 2.26 (m, 1H), 2.27 – 2.11 (m, 3H), 2.07 – 1.88 (m, 2H), 1.74 – 1.61 (m, 2H), 1.38 – 1.27 (m, 2H), 1.11 – 1.00 (m, 1H), 0.93 (s, 9H), 0.65 – 0.54 (m, 1H), 0.46 – 0.35 (m, 1H), 0.34 – 0.26 (m, 1H), 0.20 – 0.09 (m, 1H); $^{13}$**C NMR** (151 MHz, $CD_3OD$) δ 176.2, 161.7 (q, *J* = 37.6 Hz), 160.6, 160.3, 147.4, 133.0, 132.9, 130.5, 130.4, 128.7, 121.1, 117.6 (q, *J* = 293.0 Hz), 115.0, 113.3, 107.8, 107.7, 72.4, 57.0, 56.2, 45.1, 42.3, 35.0, 31.1, 31.0, 29.8, 28.4, 27.1, 18.2, 5.8, 4.5; **ESI-MS** m/z 508.6 (M+H$^+$). Spectra in agreement with literature.[5]

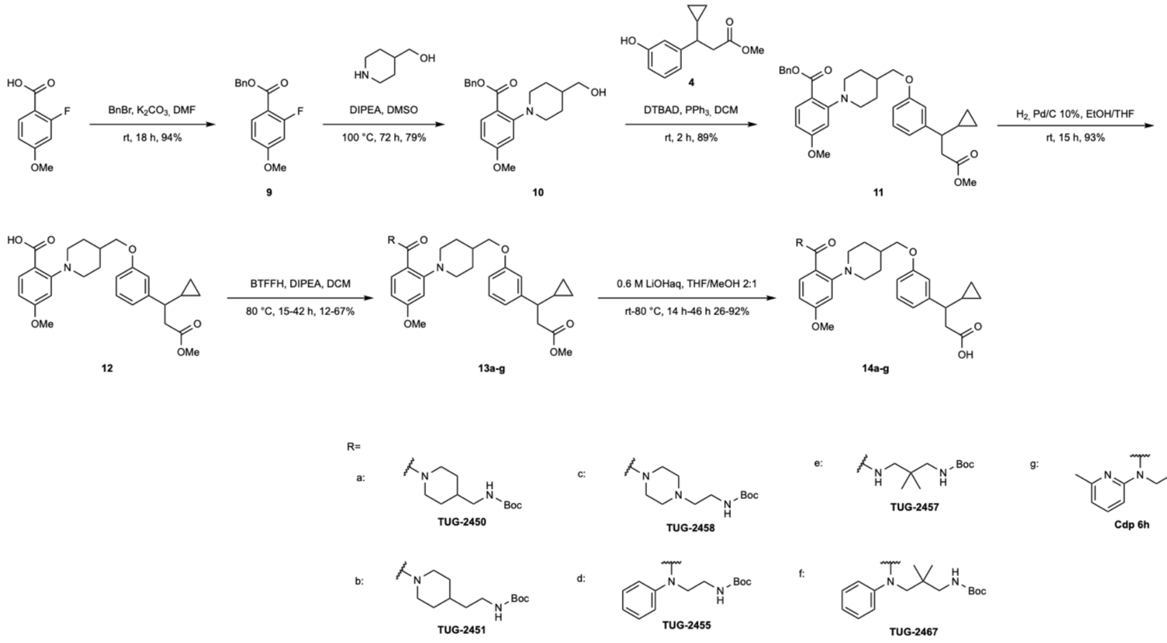

**Benzyl 2-(4-(hydroxymethyl)piperidin-1-yl)-4-methoxybenzoate (10)**

To a solution of compound **9**[4] (100 mg, 0.38 mmol) and piperidin-4-ylmethanol (43 mg, 0.38 mmol) in DMSO (1.92 mL), DIPEA (134 µL, 0.77 mmol) was added dropwise. The vial was sealed, and the resulting yellow solution was stirred at 100 °C for 72 h. Then, the reaction mixture as a dark orange solution, was cooled to rt, diluted with $H_2O$ and extracted with EtOAc (×3). The combined organic layers were washed with brine, dried over $MgSO_4$, filtered and concentrated *in vacuo*. The crude reaction mixture was purified by flash column chromatography (EtOAc:*n*-heptane, 1:1) affording 108 mg (79%) of a yellow oil: $R_f$ = 0.25 (EtOAc:*n*-heptane, 1:1); **$^1$H NMR** (600 MHz, $CD_3OD$) δ 7.75 (d, *J* = 8.6 Hz, 1H), 7.47 – 7.42 (m, 2H), 7.40 – 7.33 (m, 2H), 7.34 – 7.28 (m, 1H), 6.56 (d, *J* = 2.4 Hz, 1H), 6.53 (dd, *J* = 8.7, 2.5 Hz, 1H), 5.29 (s, 2H), 3.80 (s, 3H), 3.38 (d, *J* = 6.6 Hz, 2H), 3.30 – 3.23 (m, 2H), 2.64 (td, *J* = 11.9, 2.4 Hz, 2H), 1.77 – 1.66 (m, 2H), 1.57 – 1.47 (m, 1H), 1.33 (qd, *J* = 12.4, 3.9 Hz, 2H); **$^{13}$C NMR** (151 MHz, $CD_3OD$) δ 168.9, 165.1, 157.1, 138.0, 134.8, 129.6, 129.4, 129.2, 117.0, 106.9, 106.0, 68.1, 67.4, 55.8, 54.1, 39.7, 30.1; **ESI-MS** m/z 365.3 ($M+H^+$). Spectra in agreement with literature.[5]

**Benzyl 2-(4-((3-(1-cyclopropyl-3-methoxy-3-oxopropyl)phenoxy)methyl)piperidin-1-yl)-4-methoxybenzoate (11)**

In a flask under Ar, compound **10** (69 mg, 0.19 mmol) was dissolved in anh. DCM (2.00 mL), and a solution of compound **4** (56 mg, 0.25 mmol) in anh. DCM (1.90 mL) was added, followed by $PPh_3$ (76 mg, 0.29 mmol). The reaction mixture was cooled to 0 °C and DTBAD (67 mg, 0.29 mmol) was added. The reaction mixture was allowed to warm to rt and was stirred for 2 h under Ar. Then, the reaction was concentrated *in vacuo* and the crude mixture was directly purified by flash column chromatography (DCM:EtOAc, 97:3) to afford 96 mg (89%) of a pale yellow oil: $R_f$ = 0.78 (DCM:EtOAc, 95:5); **$^1$H NMR** (400 MHz, $CDCl_3$) δ 7.87 (d, *J* = 8.7 Hz, 1H), 7.49 – 7.42 (m, 2H), 7.42 – 7.28 (m, 3H), 7.21 (t, *J* = 7.8 Hz, 1H), 6.89 – 6.73 (m, 3H), 6.61 – 6.41 (m, 2H), 5.32 (s, 2H), 3.90 – 3.75 (m, 5H), 3.62 (s, 3H), 3.39 (d, *J* = 11.6 Hz, 2H), 2.85 – 2.63 (m, 4H), 2.40 – 2.30 (m, 1H), 2.01 – 1.79 (m, 3H), 1.57 – 1.44 (m, 2H), 1.09 – 0.94 (m, 1H), 0.64 – 0.52 (m, 1H), 0.51 – 0.36 (m, 1H), 0.32 – 0.19 (m, 1H), 0.20 – 0.11 (m, 1H); **$^{13}$C NMR** (151 MHz, $CDCl_3$) δ 173.0, 166.9, 163.6, 159.3, 156.0, 146.0, 136.6, 134.4, 129.5, 128.7, 128.5, 119.8, 114.4, 114.0,

113.6, 112.2, 105.6, 105.2, 72.6, 66.4, 55.5, 53.0, 51.6, 47.4, 41.7, 36.0, 29.3, 17.2, 5.4, 4.2; **ESI-MS** m/z 558.7 ($M+H^+$). Spectra in agreement with literature.[5]

**2-(4-((3-(1-Cyclopropyl-3-methoxy-3-oxopropyl)phenoxy)methyl)piperidin-1-yl)-4-methoxybenzoic acid (12)**

Compound synthesised similar to Furukawa *et al.*[5] but with a hydrogen ballon instead of an H-Cube: $\boldsymbol{R_f}$ = 0.11 (EtOAc:*n*-heptane, 1:1); $^{1}$H NMR (600 MHz, $CDCl_3$) δ 8.26 (d, *J* = 8.4 Hz, 1H), 7.22 (t, *J* = 7.9 Hz, 1H), 6.93 – 6.88 (m, 2H), 6.84 (d, *J* = 7.7 Hz, 1H), 6.81 – 6.74 (m, 2H), 3.90 – 3.86 (m, 5H), 3.62 (s, 3H), 3.19 (d, *J* = 11.6 Hz, 2H), 2.99 – 2.91 (m, 2H), 2.80 – 2.67 (m, 2H), 2.38 – 2.31 (m, 1H), 2.14 – 2.01 (m, 3H), 1.74 – 1.64 (m, 2H), 1.07 – 0.98 (m, 1H), 0.61 – 0.55 (m, 1H), 0.48 – 0.40 (m, 1H), 0.30 – 0.23 (m, 1H), 0.20 – 0.12 (m, 1H); $^{13}$C NMR (151 MHz, $CDCl_3$) δ 172.9, 167.3, 163.9, 159.0, 153.0, 146.1, 134.3, 129.6, 120.1, 117.8, 113.9, 112.23, 112.17, 108.8, 71.9, 55.8, 54.3, 51.7, 47.4, 41.7, 35.5, 29.5, 17.2, 5.4, 4.2; **ESI-MS** m/z 468.2 ($M+H^+$). Spectra in agreement with literature.[5]

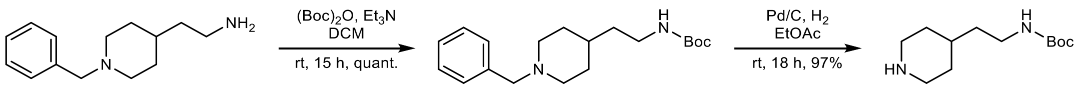


A B

***tert*-Butyl (2-(1-benzylpiperidin-4-yl)ethyl)carbamate (A)**

To a solution of 2-(1-benzylpiperidin-4-yl)ethan-1-amine (150 mg, 0.69 mmol) in anh. DCM (1.40 mL), di-*tert*-butyl dicarbonate ($Boc_2O$) (215 mg, 0.98 mmol) was added. The solution was cooled to 0 °C and $Et_3N$ (124 µL, 0.89 mmol) was added dropwise. The reaction mixture was stirred at rt for 15 h. Upon completion, the crude mixture was diluted with EtOAc, quenched with sat. aq. $NH_4Cl$, and the aqueous phase was extracted with EtOAc (×3). The combined organic layers were washed with brine, dried over $MgSO_4$, filtered and concentrated *in vacuo* to give 237 mg (quant. crude) of a yellow oil that was used in the next step without further purification: $\boldsymbol{R_f}$ = 0.79 (DCM:MeOH, 9:1); **$^{1}$H NMR** (400 MHz, $CDCl_3$) δ 7.42 – 7.23 (m, 5H), 4.45 (br s, 1H), 3.54 (s, 2H), 3.21 – 3.07 (m, 2H), 3.00 – 2.84 (m, 2H), 2.12 – 1.89 (m, 2H), 1.71 – 1.64 (m, 2H), 1.43 (m, 14H); **$^{13}$C NMR** (101 MHz, $CDCl_3$) δ 156.1, 129.6, 128.4, 127.4, 79.2, 63.3, 53.7, 38.4, 36.8, 33.3, 32.0, 31.4, 28.6; ESI-MS m/z 319.4 ($M+H^+$).

***tert*-Butyl (2-(piperidin-4-yl)ethyl)carbamate (B)**

To a solution of compound **A** (197 mg, 0.62 mmol) in EtOAc (1.20 mL) under $N_2$, Pd/C 10% (20 mg, 10 wt %) was added. The flask was evacuated and backfilled with Ar (×3) and then with $H_2$ (×3). The reaction mixture was stirred at rt for 18 h under $H_2$ atmosphere. Upon completion, the reaction mixture was filtered through a Celite® pad, that was washed with EtOAc and the filtrate was concentrated *in vacuo* to give 137 mg (97% crude) of a colorless oil that was used in the next step without further purification: **$^1$H NMR** (400 MHz, $CDCl_3$) δ 4.49 (br s, 1H), 3.19 – 3.03 (m, 4H), 2.64 – 2.53 (m, 3H), 1.73 – 1.64 (m, 2H), 1.46 – 1.35 (m, 12H), 1.23 – 1.07 (m, 2H); **$^{13}$C NMR** (101 MHz, $CDCl_3$) δ 156.1, 79.2, 46.6, 38.1, 37.4, 33.8, 33.1, 28.6; **ESI-MS** m/z 229.2 ($M+H^+$). Spectra in agreement with literature.[6]

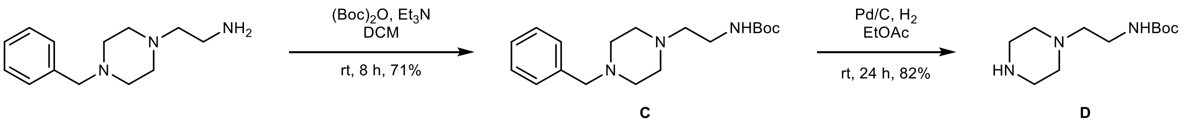


***tert*-Butyl (2-(4-benzylpiperazin-1-yl)ethyl)carbamate (C)**

Compound synthesized similar to Hauel *et al.*[7] but with addition of base. To a solution of 2-(4-benzylpiperazin-1-yl)ethan-1-amine (130 mg, 0.59 mmol) in anh. DCM (1.20 mL), $Boc_2O$ (194 mg, 0.89 mmol) was added. The solution was cooled to 0 °C and $Et_3N$ (124 μL, 0.89 mmol) was added dropwise. The resulting solution was stirred at rt for 8 h. Upon completion, the reaction mixture was diluted with EtOAc, quenched with sat. aq. $NH_4Cl$ and the aqueous phase was extracted with EtOAc (×3), the combined organic layers were washed with brine, dried over $MgSO_4$, filtered and concentrated *in vacuo* to give 135 mg (71% crude) of a yellow oil that was used without further purification: $\boldsymbol{R_f}$ = 0.23 (EtOAc); **$^1$H NMR** (600 MHz, $CDCl_3$) δ 7.33 – 7.29 (m, 4H), 7.27 – 7.23 (m, 1H), 5.01 (s, 1H), 3.53 (s, 2H), 3.25 – 3.20 (m, 2H), 2.60 – 2.40 (m, 10H), 1.45 (s, 9H); **$^{13}$C NMR** (151 MHz, $CDCl_3$) δ 156.1, 137.9, 129.4, 128.4, 127.3, 79.3, 63.1, 57.3, 52.99, 52.97, 37.2, 28.6. Spectra in agreement with literature.[8]

***tert*-Butyl (2-(piperazin-1-yl)ethyl)carbamate (D)**

Compound synthesized similar to Hauel *et al.*[7] but using EtOH as solvent. To a solution of compound **C** (95 mg, 0.30 mmol) in abs. EtOH (0.56 mL) under $N_2$, Pd/C 10% (10 mg,

10 wt%) was added. The flask was evacuated and backfilled with Ar (×3), then with $H_2$ (×3) and the reaction mixture was stirred for 24 h at rt under $H_2$ atmosphere. Upon completion, the reaction mixture was filtered through a Celite® pad, that was washed with EtOAc and the filtrate was concentrated *in vacuo* to give 75 mg (82% crude) of a colourless oil that was used in the next step without further purification: **$^{1}$H NMR** (400 MHz, $CDCl_3$) δ 4.96 (s, 1H), 3.27 – 3.17 (m, 2H), 2.97 – 2.83 (m, 3H), 2.52 – 2.37 (m, 8H), 1.44 (s, 9H); **$^{13}$C NMR** (101 MHz, $CDCl_3$) δ 156.1, 79.3, 57.8, 54.0, 45.9, 37.1, 28.6; **ESI-MS** m/z 230.2 ($M+H^+$). Spectra in agreement with literature.[9]

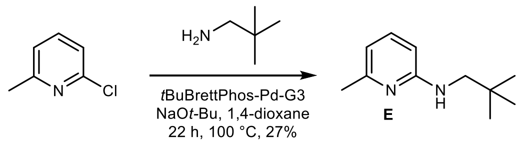


**6-Methyl-*N*-neopentylpyridin-2-amine (E)**

To a Schlenk flask under Ar, *t*BuBrettPhos-Pd-G3 (7 mg, 1 mol%), NaO*t*-Bu (90 mg, 0.94 mmol), 2-chloro-6-methylpyridine (86 µL, 0.78 mmol) and 2,2-dimethylpropan-1-amine (185 µL, 1.57 mmol) were added. The Schlenk flask was evacuated and backfilled with Ar (×3) before adding anh. 1,4-dioxane (0.78 mL). The reaction mixture was stirred for 22 h at 100 °C then, diluted with EtOAc and filtered through a Celite® pad. The filtrate was concentrated *in vacuo* and purified by flash column chromatography (EtOAc:*n*-heptane, 5:95 → 10:90) to give 26 mg (27%) of a yellow oil: ***R$_f$*** = 0.64 (EtOAc:*n*-heptane, 1:1); **$^{1}$H NMR** (400 MHz, $CDCl_3$) δ 7.32 (t, *J* = 7.8 Hz, 1H), 6.41 (d, *J* = 7.2 Hz, 1H), 6.20 (d, *J* = 8.3 Hz, 1H), 4.65 (br s, 1H), 2.99 (d, *J* = 6.1 Hz, 2H), 2.36 (s, 3H), 0.98 (s, 9H); **$^{13}$C NMR** (151 MHz, $CDCl_3$) δ 159.2, 156.8, 138.2, 112.0, 102.3, 54.4, 32.1, 27.7, 24.3; **ESI-MS** m/z 179.3 ($M+H^+$). Spectra in agreement with literature.[10]

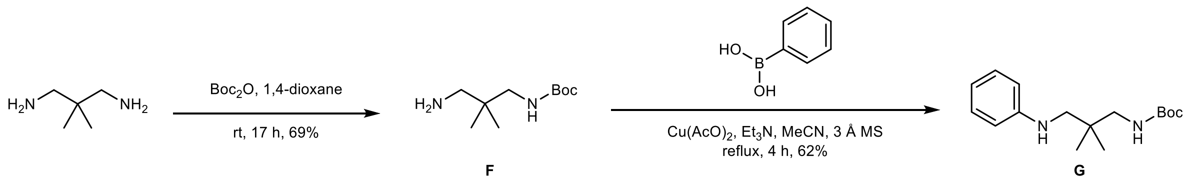


***tert*-Butyl (3-amino-2,2-dimethylpropyl)carbamate (F)**

Compound synthesized similar to Chadwick *et al.*[11] but using 2,2-dimethylpropane-1,3-diamine. To a solution of 2,2-dimethylpropane-1,3-diamine (1.00 mL, 8.33 mmol) in 1,4-dioxane (3.00 mL), $Boc_2O$ (227 mg, 1.04 mmol) dissolved in 1,4-dioxane (3.00 mL) was added dropwise over 5 min. The resulting suspension was stirred at rt for 17 h. Upon completion, the reaction mixture was concentrated *in vacuo*. The residue was resuspended in $H_2O$, filtered, and the water phase was extracted with DCM (×4). The combined organic layers were concentrated *in vacuo* to give 145 mg (69% crude) of a white solid which was used in the next step without further purification: **$^1$H NMR** (400 MHz, DMSO-$d_6$) δ 6.80 (t, *J* = 5.9 Hz, 1H), 2.77 (d, *J* = 6.4 Hz, 2H), 2.24 (s, 2H), 1.53 (s, 1H), 1.38 (s, 9H), 0.72 (s, 6H); **$^{13}$C NMR** (151 MHz, DMSO-$d_6$) δ 156.1, 77.3, 49.9, 47.4, 36.1, 28.2, 23.2; **ESI-MS** m/z 203.4 (M+H$^+$). Spectra in agreement with literature.[12]

**_tert_-Butyl (2,2-dimethyl-3-(phenylamino)propyl)carbamate (G)**

Compound synthesized similar to Vantourout *et al.*[13] but using phenyl boronic acid and compound **F**. To a microwave vial under Ar, 3Å molecular sieves were added and the vial was evacuated and backfilled with Ar (×3). Phenyl boronic acid (12 mg, 0.09 mmol), compound **F** (30 mg, 0.15 mmol) and $Cu(AcO)_2$ (18 mg, 0.09 mmol) were added. The vial was purged again with Ar, anh. MeCN (200 µL) was added followed by dropwise addition of $Et_3N$ (28 µL, 0.09 mmol). The vial was capped, and the reaction was stirred at reflux for 4 h. Upon completion, the reaction mixture was cooled to rt, diluted with EtOAc and filtered through a Celite® pad. The filtrate was concentrated *in vacuo* and purified by flash column chromatography (EtOAc:*n*-heptane, 1:9) to give 17 mg (62%) of a white solid: $R_f$ = 0.71 (EtOAc:*n*-heptane, 1:1); **$^1$H NMR** (400 MHz, $CDCl_3$) δ 7.22 – 7.13 (m, 2H), 6.74 – 6.68 (m, 3H), 4.76 (br s, 1H), 3.07 (d, *J* = 6.8 Hz, 2H), 2.93 (s, 2H), 1.44 (s, 9H), 0.97 (s, 6H); **$^{13}$C NMR** (151 MHz, $CDCl_3$) δ 156.8, 148.4, 129.4, 117.9, 113.8, 79.7, 52.3, 48.5, 36.3, 28.5, 24.0; **ESI-MS** m/z 279.2 (M+H$^+$).

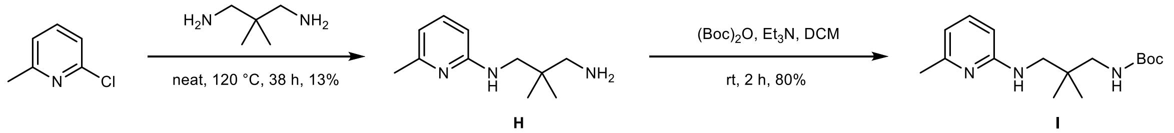


**2,2-Dimethyl-*N*1-(6-methylpyridin-2-yl)propane-1,3-diamine (H)**

In a microwave vial, 2-chloro-6-methylpyridine (171 µL, 1.57 mmol) and 2,2-dimethylpropane-1,3-diamine (1.13 mL, 9.42 mmol) were added, and the vial was capped. The reaction mixture was stirred at 120 °C for 38 h. The reaction mixture was cooled to rt, diluted with $H_2O$ and extracted with EtOAc (×4). The organic layers were combined and washed with brine, dried over $MgSO_4$, filtered and concentrated *in vacuo*, and the crude material was purified by flash column chromatography (10% MeOH in DCM + 2% $NH_4OH$ → 15% MeOH in DCM + 2% $NH_4OH$) affording 40 mg (13%) of a colourless oil: $R_f$ = 0.17 (DCM:MeOH, 10% + 2% $NH_4OH$); **$^1$H NMR** (400 MHz, $CDCl_3$) δ 7.27 (d, *J* = 15.5 Hz, 1H), 6.38 (d, *J* = 7.2 Hz, 1H), 6.19 (d, *J* = 8.3 Hz, 1H), 4.91 (br s, 1H), 3.15 (s, 2H), 2.94 (br s, 2H), 2.56 (s, 2H), 2.35 (s, 3H), 0.95 (s, 6H); **$^{13}$C NMR** (101 MHz, $CDCl_3$) δ 158.9, 156.7, 138.0, 112.0, 103.8, 50.1, 50.0, 35.9, 24.3, 23.9; **ESI-MS** m/z 194.2 ($M+H^+$).

**_tert_-Butyl (2,2-dimethyl-3-((6-methylpyridin-2-yl)amino)propyl)carbamate (I)**

To a solution of compound **H** (40 mg, 0.21 mmol) in anh. DCM (420 µL), $Boc_2O$ (68 mg, 0.31 mmol) was added. The solution was cooled to 0 °C and $Et_3N$ (42 µL, 0.31 mmol) was added. The resulting solution was stirred at rt for 2 h. Upon completion, the reaction mixture was diluted with EtOAc, quenched with sat. aq. $NH_4Cl$ and the aqueous phase was extracted with EtOAc (×3). The combined organic layers were washed with sat. aq. $NH_4Cl$ (×2), dried over $MgSO_4$, filtered and concentrated *in vacuo* to give 48 mg (80% crude) of a white amorphous solid that was used in the next step without further purification: $R_f$ = 0.61 (EtOAc:*n*-heptane, 1:1); **$^1$H NMR** (400 MHz, $CDCl_3$) δ 7.39 – 7.34 (m, 1H), 6.41 (d, *J* = 7.2 Hz, 1H), 6.29 (d, *J* = 8.4 Hz, 1H), 3.18 (d, *J* = 6.4 Hz, 2H), 2.95 (d, *J* = 6.7 Hz, 2H), 2.46 (s, 3H), 1.45 (s, 9H), 0.95 (s, 6H); **ESI-MS** m/z 294.4 ($M+H^+$).

**Methyl 3-(3-((1-(2-(4-(((_tert_-butoxycarbonyl)amino)methyl)piperidine-1-carbonyl)-5-methoxyphenyl)piperidin-4-yl)methoxy)phenyl)-3-cyclopropylpropanoate (13a)**

Compound **12** (11 mg, 0.02 mmol) was added to a microwave vial and dissolved in anh. DCM (100 µL). Fluoro-*N*,*N*,*N'*,*N'*-bis(tetramethylene)formamidinium hexafluorophosphate[14] (BTFFH) (10 mg, 0.03 mmol) and DIPEA (18 µL, 0.11 mmol) were added and the reaction was stirred at rt under Ar for 30 min. Then, *tert*-butyl (piperidin-4-ylmethyl)carbamate (7 mg, 0.03 mmol) was added, the vial was sealed and the reaction mixture was stirred at 80 °C for 21 h. The reaction mixture was cooled to rt, diluted with $H_2O$ and extracted with EtOAc (×3). The combined organic layers were washed with brine, dried over $MgSO_4$, filtered and concentrated *in vacuo*. The crude material was purified by flash column chromatography (EtOAc:*n*-heptane, 1:1) affording 5 mg (35%) of an off-white solid: ***R**f* = 0.68 (EtOAc); **$^{1}$H NMR** (600 MHz, $CD_3OD$) δ 7.21 – 7.14 (m, 2H), 6.83 – 6.79 (m, 2H), 6.79 – 6.75 (m, 1H), 6.74 – 6.68 (m, 2H), 4.70 – 4.61 (m, 1H), 3.94 – 3.84 (m, 2H), 3.83 (s, 3H), 3.57 (s, 3H), 3.55 – 3.50 (m, 1H), 3.45 – 3.42 (m, 2H), 3.06 (br s, 1H), 2.98 (d, *J* = 6.7 Hz, 2H), 2.93 – 2.66 (m, 5H), 2.33 – 2.26 (m, 1H), 2.00 – 1.51 (m, 7H), 1.42 (s, 9H), 1.26 – 1.17 (m, 2H), 1.10 – 1.01 (m, 1H), 0.61 – 0.54 (m, 1H), 0.44 – 0.36 (m, 1H), 0.28 – 0.21 (m, 1H), 0.17 – 0.09 (m, 1H); **$^{13}$C NMR** (151 MHz, $CD_3OD$) δ 174.6, 171.9, 163.2, 160.6, 158.6, 151.9, 147.1, 130.8, 130.4, 123.8, 120.8, 114.9, 113.4, 109.1, 106.7, 79.9, 73.3, 56.7, 55.9, 52.0, 46.7, 43.2, 42.3, 38.0, 36.9, 31.3, 30.7, 30.4, 28.8, 18.1, 5.8, 4.6; **ESI-MS** m/z 664.8 ($M+H^+$).

**2,2,2-Trifluoroacetic acid-3-(3-((1-(2-(4-(((*tert*-butoxycarbonyl)amino)methyl)-piperidine-1-carbonyl)-5-methoxyphenyl)piperidin-4-yl)methoxy)phenyl)-3-cyclo-propylpropanoic acid (TUG-2450 - 14a)**

Compound **13a** (5 mg, 7.5 µmol) was dissolved in a mixture of THF/MeOH (2:1, 100 µL), 0.6 M $LiOH_{aq}$ (10 µL, 6.0 µmol) was added, and the reaction mixture was stirred at 50 °C for 13 h. Then, additional 0.6 M $LiOH_{aq}$ (10 µL, 6.0 µmol) was added, and the reaction was stirred at 80 °C for 23 h. Upon completion, the reaction mixture was cooled to rt, diluted with $H_2O$, acidified with 1 M $HCl_{aq}$ and extracted with EtOAc (×3). The organic layers were combined, washed with brine, dried over $MgSO_4$, filtered and concentrated *in vacuo*. The crude compound was purified by preparative HPLC (25 to 100% B over 15 min) and the combined fractions were lyophilized affording 3 mg (39%) of an off-white solid as TFA salt: ***t**R* = 12.9 min (purity 96.10% by HPLC); **$^{1}$H NMR** (600 MHz, $CD_3OD$) δ 7.19 (t, *J* = 8.0 Hz, 1H), 7.15 (d, *J* = 8.2 Hz, 1H), 6.85 – 6.80 (m, 2H), 6.79 – 6.74 (m, 1H), 6.70 – 6.64 (m, 2H), 4.65 (d, *J* = 13.0 Hz, 1H), 3.90 (d, *J* = 5.6 Hz, 2H), 3.82 (s, 3H), 3.54 – 3.47 (m, 1H), 3.44 – 3.36 (m, 2H), 3.09 – 3.01 (m, 1H), 2.97 (d, *J* = 6.7 Hz, 2H), 2.90 – 2.73 (m, 4H), 2.68 (dd, *J* = 14.9, 8.5 Hz, 1H), 2.35 – 2.27 (m, 1H), 1.98 – 1.88 (m, 3H), 1.86 – 1.79 (m, 1H), 1.78 – 1.69 (m, 1H), 1.66 – 1.50 (m, 3H), 1.42 (s, 9H), 1.27 – 1.15 (m, 2H), 1.10 – 1.01 (m, 1H), 0.62 – 0.55 (m, 1H), 0.44 – 0.36 (m, 1H), 0.33 – 0.26 (m, 1H), 0.18 – 0.11 (m, 1H); **$^{13}$C NMR** (151 MHz, $CD_3OD$) δ 176.2, 172.1, 163.1, 160.6, 158.7, 152.5, 147.2, 130.6, 130.3, 124.1, 120.9, 115.0, 113.4, 108.6, 106.5, 80.0, 73.4, 55.9, 46.7, 43.1, 42.4, 38.0, 37.1, 31.3, 30.7, 30.6, 30.5, 28.8, 18.2, 5.8, 4.6; **$^{19}$F NMR** (376 MHz,

$CD_3OD$) δ -77.1; **HRMS (MALDI)** calcd for $C_{37}H_{52}N_3O_7^+$ ($M+H^+$) 650.3799, found 650.3793.

**2,2,2-Trifluoroacetic acid-3-(3-((1-(2-(4-(2-((*tert*-butoxycarbonyl)amino)ethyl)-piperidine-1-carbonyl)-5-methoxyphenyl)piperidin-4-yl)methoxy)phenyl)-3-cyclo-propylpropanoic acid (TUG-2451 - 14b)**

**Step 1:** Compound **12** (5 mg, 0.01 mmol) was added to a microwave vial and dissolved in anh. DCM (50 µL). BTFFH (5 mg, 0.01 mmol) and DIPEA (8 µL, 0.04 mmol) were added and the reaction was stirred under Ar at rt for 30 min. Compound **B** (4 mg, 0.01 mmol) dissolved in anh. DCM (50 µL) was then added, the vial was sealed, and the reaction mixture was stirred at 80 °C for 24 h. The reaction mixture was cooled to rt, diluted with $H_2O$ and extracted with EtOAc (×3). The combined organic layers were washed with brine, dried over $MgSO_4$, filtered and concentrated *in vacuo* affording 8 mg of **13b** as a pale yellow oil that was used in the next step without further purification: $\boldsymbol{R_f}$ = 0.71 (EtOAc); **ESI-MS** m/z 678.8 ($M+H^+$).

**Step 2:** To a solution of the compound from Step 1 (8 mg, 0.01 mmol) in a mixture of THF/MeOH (2:1, 1 mL), 0.6 M $LiOH_{aq}$ (183 µL, 0.11 mmol) was added, and the reaction mixture was stirred at 60 °C for 14 h. The reaction mixture was then diluted with $H_2O$, acidified with 1 M $HCl_{aq}$ and extracted with EtOAc (×3). The organic layers were combined, washed with brine, dried over $MgSO_4$, filtered and concentrated *in vacuo*. The crude was purified by preparative HPLC (25 to 100% B over 15 min) and the combined fractions were lyophilized, affording 2 mg (26% after 2 steps) of the title compound as an off-white solid TFA salt: $\boldsymbol{t_R}$ = 13.2 min (purity 96.13% by HPLC); **$^1$H NMR** (600 MHz, $CD_3OD$) δ 7.26 (d, *J* = 8.5 Hz, 1H), 7.19 (t, *J* = 7.9 Hz, 1H), 6.88 (s, 1H), 6.86 – 6.74 (m, 4H), 4.63 (s, 1H), 3.91 (d, *J* = 5.5 Hz, 2H), 3.85 (s, 3H), 3.68 – 3.54 (m, 1H), 3.52 – 3.47 (m, 2H), 3.13 – 3.03 (m, 5H), 2.83 (s, 1H), 2.76 (dd, *J* = 14.9, 6.6 Hz, 1H), 2.69 (dd, *J* = 14.9, 8.5 Hz, 1H), 2.32 (td, *J* = 9.0, 6.6 Hz, 1H), 2.13 – 1.96 (m, 3H), 1.87 (s, 1H), 1.77 – 1.58 (m, 4H), 1.47 – 1.40 (m, 11H), 1.26 – 1.20 (m, 2H), 1.10 – 1.01 (m, 1H), 0.63 – 0.55 (m, 1H), 0.44 – 0.37 (m, 1H), 0.33 – 0.26 (m, 1H), 0.18 – 0.09 (m, 1H); **$^{13}$C NMR** (151 MHz, $CD_3OD$) δ 176.2, 171.4, 163.3, 160.5, 158.6, 150.5, 147.3, 131.1, 130.4, 121.0, 115.0, 113.4, 110.4, 107.3, 79.9, 73.0, 64.4, 56.1, 43.7, 42.4, 38.7, 37.5, 36.4, 34.7, 33.0, 30.0,

28.8, 18.2, 5.8, 4.6; **$^{19}$F NMR** (376 MHz, $CD_3OD$) δ -77.4; **HRMS (MALDI)** calcd for $C_{38}H_{54}N_3O_7^+$ (M+H$^+$) 664.3956, found 664.3957.

**Methyl 3-(3-((1-(2-(4-(2-((*tert*-butoxycarbonyl)amino)ethyl)piperazine-1-carbonyl)-5-methoxyphenyl)piperidin-4-yl)methoxy)phenyl)-3-cyclopropylpropanoate (13c)**

Compound **12** (25 mg, 0.05 mmol) was added to a microwave vial and dissolved in anh. DCM (100 µL). BTFFH (26 mg, 0.08 mmol) and DIPEA (42 µL, 0.24 mmol) were added and the reaction was stirred under Ar at rt for 30 min. Compound **D** (16 mg, 0.07 mmol) dissolved in anh. DCM (200 µL) was added, the vial was sealed, and the reaction was stirred at 80 °C for 24 h. The reaction mixture was cooled to rt, diluted with $H_2O$ and extracted with EtOAc (×3). The combined organic layers were washed with brine, dried over $MgSO_4$, filtered and concentrated *in vacuo*. The crude compound was purified by automated flash column chromatography (DCM:EtOAc, 1:1 → 0:100) affording 7 mg (19%) of an off-white solid: ***R$_f$*** = 0.15 (EtOAc); **$^1$H NMR** (600 MHz, $CD_3OD$) δ 7.19 (t, *J* = 8.1 Hz, 1H), 7.14 (d, *J* = 8.3 Hz, 1H), 6.82 – 6.79 (m, 2H), 6.78 – 6.76 (m, 1H), 6.66 – 6.60 (m, 2H), 4.02 – 3.96 (m, 1H), 3.91 – 3.85 (m, 2H), 3.81 (s, 3H), 3.62 – 3.54 (m, 4H), 3.48 – 3.37 (m, 2H), 3.27 – 3.17 (m, 4H), 2.91 (td, *J* = 11.6, 2.6 Hz, 1H), 2.82 – 2.76 (m, 1H), 2.76 – 2.70 (m, 2H), 2.68 – 2.63 (m, 1H), 2.58 (td, *J* = 12.2, 2.4 Hz, 1H), 2.53 – 2.45 (m, 3H), 2.32 – 2.24 (m, 2H), 2.04 – 1.98 (m, 1H), 1.96 – 1.88 (m, 1H), 1.85 – 1.81 (m, 1H), 1.62 – 1.47 (m, 2H), 1.41 (s, 9H), 1.10 – 1.01 (m, 1H), 0.61 – 0.54 (m, 1H), 0.44 – 0.37 (m, 1H), 0.28 – 0.21 (m, 1H), 0.19 – 0.09 (m, 1H); **$^{13}$C NMR*** (151 MHz, $CD_3OD$) δ 174.6, 172.5, 163.2, 160.7, 158.4, 153.2, 147.0, 130.7, 130.4, 123.9, 120.8, 114.9, 113.5, 108.2, 106.2, 80.1, 73.5, 58.7, 55.8, 55.7, 54.3, 53.9, 52.0, 51.4, 48.2, 42.8, 42.4, 42.3, 38.4, 37.2, 30.8, 30.7, 28.8, 26.5, 18.1, 5.8, 4.6; **ESI-MS** m/z 679.8 (M+H$^+$).

*Splitting of peaks was observed.

**3-(3-((1-(2-(4-(2-((*tert*-Butoxycarbonyl)amino)ethyl)piperazine-1-carbonyl)-5-methoxyphenyl)piperidin-4-yl)methoxy)phenyl)-3-cyclopropylpropanoic acid hydrochloride (TUG-2458 - 14c)**

To a solution of compound **13c** (6 mg, 0.01 mmol) dissolved in a mixture of THF/MeOH (2:1, 1 mL), 0.6 M $LiOH_{aq}$ (204 µL, 0.06 mmol) was added, and the reaction mixture was stirred at rt for 20 h. Upon completion, the reaction was diluted with $H_2O$, acidified with

1 M $HCl_{aq}$ and extracted with EtOAc (×3). The organic layers were combined, washed with brine, dried over $MgSO_4$, filtered and concentrated *in vacuo*. The crude was purified by flash column chromatography (DCM:EtOAc, 1:1→ DCM:MeOH, 9:1) affording 5 mg (74%) of an off-white solid as HCl salt: $t_R$ = 11.0 min (purity 95.11% by HPLC); **$^1$H NMR** (600 MHz, $CD_3OD$) δ 7.18 (t, *J* = 7.8 Hz, 1H), 7.14 (d, *J* = 8.3 Hz, 1H), 6.86 – 6.81 (m, 2H), 6.78 – 6.74 (m, 1H), 6.66 – 6.60 (m, 2H), 4.05 – 3.97 (m, 1H), 3.91 – 3.86 (m, 2H), 3.81 (s, 3H), 3.60 – 3.51 (m, 1H), 3.47 – 3.36 (m, 2H), 3.28 – 3.16 (m, 4H), 2.94 – 2.88 (m, 1H), 2.78 – 2.72 (m, 2H), 2.70 – 2.65 (m, 2H), 2.61 – 2.48 (m, 4H), 2.35 – 2.24 (m, 2H), 2.01 – 1.96 (m, 1H), 1.94 – 1.87 (m, 1H), 1.83 (d, *J* = 12.8 Hz, 1H), 1.62 – 1.47 (m, 2H), 1.42 (s, 9H), 1.09 – 1.00 (m, 1H), 0.62 – 0.55 (m, 1H), 0.44 – 0.37 (m, 1H), 0.33 – 0.26 (m, 1H), 0.18 – 0.09 (m, 1H); **$^{13}$C NMR** (151 MHz, $CD_3OD$) δ 176.6, 172.5, 163.2, 160.6, 158.4, 153.2, 147.3, 130.8, 130.3, 123.8, 120.9, 115.0, 113.3, 108.3, 106.2, 80.1, 73.5, 58.6, 55.8, 55.7, 54.2, 53.9, 51.3, 48.0, 42.7, 38.3, 37.3, 30.8, 30.7, 28.8, 18.3, 5.8, 4.6; **HRMS (MALDI)** calcd for $C_{37}H_{53}N_4O_7^+$ (M+H$^+$) 665.3908, found 665.3905.

**Methyl 3-(3-((1-(2-((2-((*tert*-butoxycarbonyl)amino)ethyl)(phenyl)carbamoyl)-5-methoxyphenyl)piperidin-4-yl)methoxy)phenyl)-3-cyclopropylpropanoate (13d)**

Compound **12** (23 mg, 0.05 mmol) was added to a microwave vial and dissolved in anh. DCM (300 µL). BTFFH (23 mg, 0.07 mmol) and DIPEA (39 µL, 0.22 mmol) were added and the reaction was stirred under Ar for 30 min. *tert*-Butyl (2-(phenylamino)ethyl)carbamate[15] (15 mg, 0.06 mmol) was added, the vial was sealed and the reaction was stirred at 80 °C for 15 h. The reaction mixture was cooled to rt, diluted with $H_2O$ and extracted with EtOAc (×3). The combined organic phases were washed with brine, dried over $MgSO_4$, filtered and concentrated *in vacuo*. The crude compound was purified by flash column chromatography (EtOAc:*n*-heptane, 1:1) affording 22 mg (67%) of a colorless oil: $R_f$ = 0.31 (EtOAc:*n*-heptane, 1:1); **$^1$H NMR** (400 MHz, $CD_3OD$) δ 7.31 – 7.00 (m, 7H), 6.87 – 6.78 (m, 3H), 6.60 – 6.46 (m, 1H), 6.25 (s, 1H), 4.02 (s, 2H), 3.93 (d, *J* = 5.5 Hz, 2H), 3.69 (s, 3H), 3.58 (s, 3H), 3.41 – 3.33 (m, 2H), 2.84 – 2.68 (m, 2H), 2.59 (s, 2H), 2.34 – 2.25 (m, 1H), 1.95 – 1.77 (m, 3H), 1.72 – 1.54 (m, 2H), 1.37 (s, 9H), 1.13 – 0.99 (m, 1H), 0.63 – 0.52 (m, 1H), 0.46 – 0.35 (m, 1H), 0.30 – 0.20 (m, 1H), 0.20 – 0.08 (m, 1H); **$^{13}$C NMR** (151 MHz, $CD_3OD$) δ 174.6, 174.3, 162.9, 160.7, 158.2, 152.9, 147.0, 143.9, 131.7, 130.4, 129.0, 127.6, 125.6, 120.7, 114.9, 113.5,

107.8, 105.9, 80.1, 73.5, 55.6, 54.8, 52.0, 42.3, 39.9, 37.2, 30.4, 28.8, 18.1, 5.8, 4.6; **ESI-MS** m/z 686.4 (M+H$^+$).

**3-(3-((1-(2-((2-((*tert*-Butoxycarbonyl)amino)ethyl)(phenyl)carbamoyl)-5-methoxy-phenyl)piperidin-4-yl)methoxy)phenyl)-3-cyclopropylpropanoic acid hydrochloride (TUG-2455 - 14d)**

To a solution of compound **13d** (19 mg, 0.03 mmol) in a mixture of THF/MeOH (2:1, 1 mL), 0.6 M $LiOH_{aq}$ (310 µL, 0.19 mmol) was added, and the reaction mixture was stirred at rt for 18 h. The reaction was diluted with $H_2O$, acidified with 1 M $HCl_{aq}$ and extracted with EtOAc (×4). The organic layers were combined, washed with brine, dried over $MgSO_4$, filtered and concentrated *in vacuo* affording 18 mg (92%) of a colourless oil as HCl salt without further purification: $\boldsymbol{t_R}$ = 12.9 min (purity 96.30% by HPLC); **$^1$H NMR** (400 MHz, $CD_3OD$) δ 7.30 – 6.99 (m, 7H), 6.87 – 6.76 (m, 3H), 6.55 – 6.48 (m, 1H), 6.27 (s, 1H), 4.02 (br s, 2H), 3.93 (d, *J* = 5.6 Hz, 2H), 3.69 (s, 3H), 3.37 (s, 2H), 2.85 – 2.53 (m, 4H), 2.38 – 2.27 (m, 1H), 1.97 – 1.82 (m, 3H), 1.71 – 1.58 (m, 2H), 1.37 (s, 9H), 1.13 – 0.99 (m, 1H), 0.64 – 0.53 (m, 1H), 0.46 – 0.35 (m, 1H), 0.35 – 0.25 (m, 1H), 0.21 – 0.07 (m, 1H); **$^{13}$C NMR** (151 MHz, $CD_3OD$) δ 176.3, 174.3, 162.9, 160.7, 158.3, 152.9, 147.3, 144.0, 131.8, 130.3, 129.0, 127.6, 125.5, 120.8, 114.9, 113.4, 107.9, 106.0, 80.1, 73.4, 55.6, 50.5, 42.4, 39.9, 37.2, 33.1, 30.8, 30.3, 28.8, 23.7, 18.3, 14.4, 5.8, 4.6; **HRMS (MALDI)** calcd for $C_{39}H_{50}N_3O_7^+$ (M+H$^+$) 672.3643, found 672.3637.

**Methyl 3-(3-((1-(2-((3-((*tert*-butoxycarbonyl)amino)-2,2-dimethylpropyl)carbamoyl)-5-methoxyphenyl)piperidin-4-yl)methoxy)phenyl)-3-cyclopropylpropanoate (13e)**

Compound **12** (26 mg, 0.05 mmol) was added to a microwave vial and dissolved in anh. DCM (100 µL). BTFFH (27 mg, 0.08 mmol) and DIPEA (43 µL, 0.25 mmol) were added and the reaction was stirred under Ar for 30 min. Compound **F** (15 mg, 0.06 mmol) was added followed by anh. DCM (200 µL). The vial was sealed, and the reaction was stirred at 80 °C for 41 h. The reaction mixture was cooled to rt, diluted with $H_2O$ and extracted with EtOAc (×3). The combined organic layers were washed with brine, dried over $MgSO_4$, filtered and concentrated *in vacuo*. The crude material was purified by automated flash column chromatography (EtOAc:*n*-heptane, 1:1) affording 20 mg (57%) of a colourless oil: $\boldsymbol{R_f}$ = 0.50 (EtOAc:*n*-heptane, 0:100→1:1); **$^1$H NMR** (600 MHz, $CD_3OD$)

δ 7.95 (d, *J* = 8.7 Hz, 1H), 7.19 (t, *J* = 8.1 Hz, 1H), 6.88 (d, *J* = 2.5 Hz, 1H), 6.85 – 6.74 (m, 4H), 6.64 (t, *J* = 6.6 Hz, 1H), 3.92 (d, *J* = 5.3 Hz, 2H), 3.85 (s, 3H), 3.58 (s, 3H), 3.30 (s, 2H), 3.23 – 3.14 (m, 2H), 2.91 (d, *J* = 6.6 Hz, 2H), 2.89 – 2.83 (m, 2H), 2.79 (dd, *J* = 14.8, 6.9 Hz, 1H), 2.72 (dd, *J* = 14.8, 8.4 Hz, 1H), 2.38 – 2.27 (m, 1H), 2.08 – 1.95 (m, 3H), 1.70 – 1.60 (m, 2H), 1.43 (s, 9H), 1.09 – 1.01 (m, 1H), 0.95 (s, 6H), 0.61 – 0.54 (m, 1H), 0.47 – 0.37 (m, 1H), 0.29 – 0.22 (m, 1H), 0.19 – 0.11 (m, 1H); **$^{13}$C NMR** (151 MHz, $CD_3OD$) δ 174.6, 169.7, 164.5, 160.6, 158.7, 155.6, 147.1, 133.7, 130.4, 121.2, 120.8, 114.9, 113.3, 110.3, 108.6, 79.9, 73.2, 56.0, 54.8, 52.0, 47.9, 42.3, 37.8, 36.9, 33.0, 30.8, 30.1, 28.8, 24.0, 18.1, 5.8, 4.6.

**3-(3-((1-(2-((3-((*tert*-Butoxycarbonyl)amino)-2,2-dimethylpropyl)carbamoyl)-5-methoxyphenyl)piperidin-4-yl)methoxy)phenyl)-3-cyclopropylpropanoic acid hydrochloride (TUG-2457 - 14e)**

To a solution of compound **13e** (18 mg, 0.03 mmol) dissolved in a mixture of THF/MeOH (2:1, 1 mL), 0.6 M $LiOH_{aq}$ (314 µL, 0.19 mmol) was added, and the reaction mixture was stirred at rt for 16 h. The reaction was diluted with $H_2O$, with 1 M $HCl_{aq}$ and extracted with EtOAc (×3). The organic layers were combined, washed with brine, dried over $MgSO_4$, filtered and concentrated *in vacuo* affording 17 mg (91%) of a pale yellow solid as HCl salt without further purification: $\boldsymbol{t_R}$ = 12.4 min (purity 97.06% by HPLC); **$^1$H NMR** (400 MHz, $CD_3OD$) δ 7.95 (d, *J* = 8.7 Hz, 1H), 7.19 (t, *J* = 7.9 Hz, 1H), 6.94 – 6.74 (m, 5H), 6.69 – 6.60 (m, 0.5H, partially exchanging), 3.92 (d, *J* = 5.2 Hz, 2H), 3.85 (s, 3H), 3.30 (s, 2H, overlapping with $CD_3OD$), 3.24 – 3.17 (m, 2H), 2.94 – 2.83 (m, 4H), 2.81 – 2.63 (m, 2H), 2.37 – 2.25 (m, 1H), 2.05 – 1.97 (m, 3H), 1.72 – 1.60 (m, 2H), 1.43 (s, 9H), 1.05 (m, 1H), 0.94 (s, 6H), 0.64 – 0.53 (m, 1H), 0.46 – 0.34 (m, 1H), 0.34 – 0.25 (m, 1H), 0.20 – 0.09 (m, 1H); **$^{13}$C NMR** (151 MHz, $CD_3OD$) 176.2, 169.7, 164.5, 160.6, 158.7, 155.5, 147.3, 133.6, 130.3, 121.0, 120.9, 115.0, 113.3, 110.4, 108.6, 80.0, 73.1, 56.0, 54.8, 48.6, 47.9, 42.4, 37.8, 36.8, 30.8, 28.8, 24.0, 18.3, 5.8, 4.6; **HRMS (MALDI)** calcd for $C_{36}H_{52}N_3O_7^+$ ($M+H^+$) 638.3799, found 638.3796.

**1-(2-((3-((*tert*-Butoxycarbonyl)amino)-2,2-dimethylpropyl)(phenyl)carbamoyl)-5-methoxyphenyl)-4-((3-(1-cyclopropyl-3-methoxy-3-oxopropyl)phenoxy)-methyl)-piperidin-1-ium 2,2,2-trifluoroacetate (13f)**

Compound **12** (23 mg, 0.05 mmol) was added to a microwave vial and dissolved in anh. DCM (100 µL). BTFFH (24 mg, 0.07 mmol) and DIPEA (39 µL, 0.22 mmol) were added and the reaction was stirred under Ar at rt for 40 min. Compound **G** (17 mg, 0.06 mmol) was added, the vial was sealed, and the reaction was stirred at 80 °C for 18 h. The reaction mixture was cooled to rt, diluted with $H_2O$ and extracted with EtOAc (×3). The organic layers were combined and washed with brine, dried over $MgSO_4$, filtered and concentrated *in vacuo*. The crude compound was purified by preparative HLPC affording 5 mg (12%) of an off-white solid TFA salt: $\boldsymbol{R_f}$ = 0.58 (EtOAc:*n*-heptane, 1:1); $\boldsymbol{t_R}$ = 11.7 min (purity 95.94% by HPLC); **$^1$H NMR** (600 MHz, $CD_3OD$) δ 7.27 – 7.06 (m, 7H), 6.84 – 6.82 (m, 2H), 6.81 – 6.78 (m, 1H), 6.55 (d, *J* = 8.2 Hz, 2H), 4.05 – 3.91 (m, 4H), 3.72 (s, 3H), 3.58 (s, 3H), 3.04 (s, 2H), 2.83 – 2.70 (m, 2H), 2.37 – 2.27 (m, 1H), 2.06 – 1.93 (m, 3H), 1.73 – 1.63 (m, 2H), 1.43 (s, 9H), 1.11 – 1.02 (m, 1H), 0.81 – 0.78 (m, 6H), 0.62 – 0.54 (m, 1H), 0.45 – 0.37 (m, 1H), 0.29 – 0.22 (m, 1H), 0.17 – 0.11 (m, 1H); **$^{13}$C NMR** (151 MHz, $CD_3OD$) δ 175.4, 174.6, 162.7, 160.7, 158.6, 153.1, 147.0, 144.9, 131.4, 130.4, 130.3, 129.0, 128.9, 127.9, 127.4, 126.3, 120.8, 114.9, 113.5, 108.3, 106.4, 80.0, 73.6, 57.3, 55.6, 52.0, 42.4, 39.3, 37.2, 28.8, 28.3, 24.9, 18.1, 5.8, 4.6; **ESI-MS** m/z 728.4 ($M+H^+$).

**2,2,2-Trifluoroacetic acid-3-(3-((1-(2-((3-((*tert*-butoxycarbonyl)amino)-2,2-dimethyl-propyl)(phenyl)carbamoyl)-5-methoxyphenyl)piperidin-4-yl)methoxy)phenyl)-3-cyclo-propylpropanoic acid (TUG-2467 - 14f)**

To a solution of compound **13f** (5 mg, 0.01 mmol) in a mixture of THF/MeOH (2:1, 1 mL), 0.6 M $LiOH_{aq}$ (250 µL, 0.15 mmol) was added, and the reaction mixture was stirred at rt for 41 h. The reaction was diluted with $H_2O$, acidified with 1 M $HCl_{aq}$ and extracted with EtOAc (×3). The organic layers were combined, washed with brine, dried over $MgSO_4$, filtered and concentrated *in vacuo* affording 4 mg (72%) of a colourless oil as TFA salt without further purification: $\boldsymbol{t_R}$ = 9.5 min (purity 97.04% by HPLC); **$^1$H NMR** (600 MHz, Acetone-$d_6$) δ 7.23 – 7.08 (m, 5H), 7.01 (t, *J* = 7.3 Hz, 1H), 6.94 – 6.90 (m, 1H), 6.88 (d, *J* = 7.6 Hz, 1H), 6.80 (dd, *J* = 8.0, 2.6 Hz, 1H), 6.57 – 6.47 (m, 2H), 6.33 (s, 1H), 3.95 (d, *J* = 5.9 Hz, 2H), 3.70 (s, 3H), 3.63 – 3.44 (m, 1H), 3.17 – 3.01 (m, 2H), 2.87 – 2.69 (m, 4H), 2.42 – 2.31 (m, 1H), 1.92 – 1.86 (m, 3H), 1.72 – 1.60 (m, 2H), 1.43 (s, 9H), 1.12 – 1.07 (m, 1H), 0.75 (s, 6H), 0.60 – 0.52 (m, 1H), 0.42 – 0.35 (m, 1H), 0.35 – 0.28 (m, 1H), 0.22 – 0.15 (m, 1H); **$^{13}$C NMR** (151 MHz, Acetone-$d_6$) δ 173.3, 173.2, 161.8, 160.2, 157.0, 147.2, 145.0,

131.3, 130.0, 128.7, 128.6, 126.7, 120.6, 114.8, 113.0, 108.3, 106.3, 78.4, 73.0, 56.8, 55.5, 47.9, 41.5, 38.9, 36.5, 28.7, 25.0, 18.1, 5.6, 4.5; **HRMS (MALDI)** calcd for $C_{42}H_{56}N_3O_7^+$ (M+H$^+$) 714.4112, found 714.4112.

**Methyl 3-cyclopropyl-3-(3-((1-(5-methoxy-2-((6-methylpyridin-2-yl)(neopentyl)-carbamoyl)-phenyl)piperidin-4-yl)methoxy)phenyl)propanoate (13g)**

Compound **12** (25 mg, 0.05 mmol) was added to a microwave vial and dissolved in anh. DCM (300 µL). BTFFH (25 mg, 0.08 mmol) and DIPEA (42 µL, 0.24 mmol) were added and the reaction was stirred under Ar for 30 min. Compound **E** (12 mg, 0.07 mmol) was added, the vial was sealed, and the reaction was stirred at 80 °C for 42 h. The reaction mixture was cooled to rt, diluted with $H_2O$ and extracted with EtOAc (×3). The combined organic phases were washed with brine, dried over $MgSO_4$, filtered and concentrated *in vacuo*. The crude compound was purified by flash column chromatography (EtOAc:*n*-heptane, 1:3) affording 6 mg (19%) of a pale yellow oil: $\boldsymbol{R_f}$ = 0.61 (EtOAc:*n*-heptane, 1:1); **$^1$H NMR** (400 MHz, $CD_3OD$) δ 7.31 – 7.13 (m, 3H), 6.96 – 6.84 (m, 1H), 6.83 – 6.75 (m, 4H), 6.59 – 6.48 (m, 2H), 6.28 (s, 1H), 4.32 – 4.07 (m, 2H), 3.91 – 3.86 (m, 2H), 3.74 (s, 3H), 3.58 (s, 3H), 3.39 (s, 1H), 2.91 – 2.59 (m, 3H), 2.43 (s, 3H), 2.36 – 2.25 (m, 1H), 1.85 – 1.59 (m, 4H), 1.12 – 0.99 (m, 1H), 0.84 (s, 9H), 0.63 – 0.52 (m, 1H), 0.47 – 0.36 (m, 1H), 0.30 – 0.20 (m, 1H), 0.20 – 0.08 (m, 1H); **$^{13}$C NMR** (151 MHz, $CD_3OD$) δ 174.7, 174.4, 163.2, 160.7, 157.8, 156.7, 152.9, 147.0, 137.9, 132.3, 130.4, 125.4, 121.2, 120.7, 119.7, 115.0, 113.4, 107.8, 105.8, 73.7, 57.3, 55.7, 52.0, 42.3, 37.3, 35.0, 30.0, 28.9, 28.8, 24.2, 18.1, 5.8, 4.6; **ESI-MS** m/z 628.8 (M+H$^+$).

**2,2,2-Trifluoroacetic acid-3-cyclopropyl-3-(3-((1-(5-methoxy-2-((6-methylpyridin-2-yl)(neopentyl)carbamoyl)phenyl)piperidin-4-yl)methoxy)phenyl)propanoic acid (Cpd 6h - 14g)**

To a solution **13g** (6 mg, 0.01 mmol) in a mixture of THF/MeOH (2:1, 1 mL), 0.6 M $LiOH_{aq}$ (210 µL, 0.13 mmol) was added, and the reaction mixture was stirred at rt for 42 h. The reaction was diluted with $H_2O$, acidified with 1 M $HCl_{aq}$ and extracted with EtOAc (×3). The combined organic layers were washed with brine, dried over $MgSO_4$, filtered and concentrated *in vacuo*. The crude material was purified by preparative HPLC (30 to 100% B over 15 min) and the combined fractions were lyophilized affording 3 mg (44%)

of a fluffy white solid as TFA salt: $t_R$ = 13.5 min (purity 99.83% by HPLC); $^1$**H NMR** (600 MHz, $CD_3OD$) δ 7.51 – 7.40 (m, 1H), 7.21 (t, *J* = 8.1 Hz, 1H), 7.15 (d, *J* = 8.6 Hz, 1H), 6.98 (d, *J* = 7.5 Hz, 1H), 6.90 – 6.83 (m, 3H), 6.82 – 6.77 (m, 1H), 6.73 – 6.61 (m, 2H), 4.14 (s, 2H), 3.94 (d, *J* = 5.7 Hz, 2H), 3.78 (s, 3H), 2.76 (dd, *J* = 14.9, 6.7 Hz, 1H), 2.70 (dd, *J* = 14.9, 8.4 Hz, 1H), 2.38 (s, 3H), 2.33 (td, *J* = 9.0, 6.7 Hz, 1H), 2.03 – 1.86 (m, 3H), 1.74 – 1.62 (m, 2H), 1.11 – 1.02 (m, 1H), 0.88 (s, 9H), 0.63 – 0.55 (m, 1H), 0.45 – 0.37 (m, 1H), 0.34 – 0.27 (m, 1H), 0.19 – 0.12 (m, 1H); $^{13}$**C NMR** (151 MHz, $CD_3OD$) δ 176.3, 173.4, 163.5, 160.5, 158.3, 156.7, 147.3, 139.0, 132.7, 130.4, 121.9, 121.0, 119.6, 115.0, 113.3, 73.0, 58.5, 56.1, 42.4, 36.4, 35.1, 29.6, 28.9, 24.0, 18.3, 5.8, 4.6; $^{19}$**F NMR** (376 MHz, $CD_3OD$) δ -77.4; **HRMS (MALDI)** calcd for $C_{37}H_{48}N_3O_5^+$ (M+H$^+$) 614.3588, found 614.3583. Spectra in agreement with literature.[5]

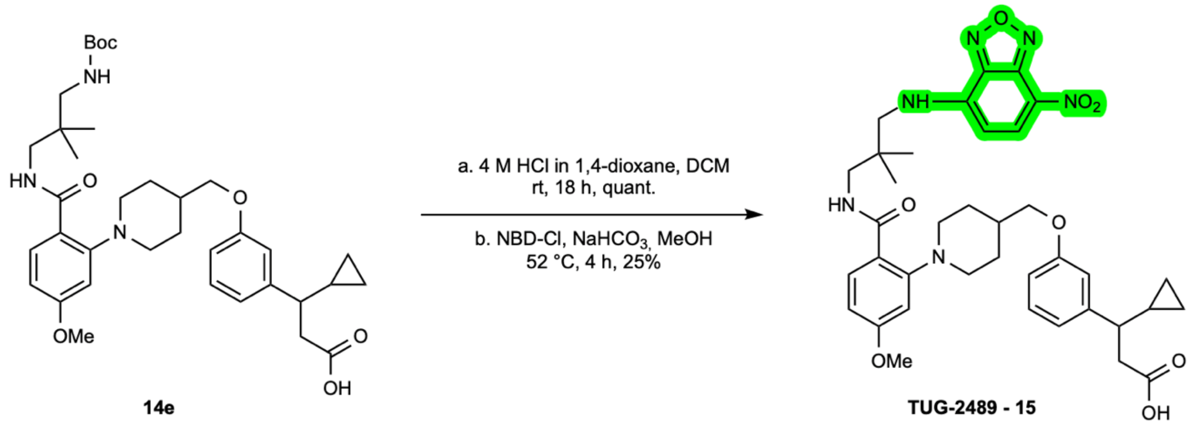


**14e** **TUG-2489 - 15**

**2,2,2-Trifluoroacetic acid-3-cyclopropyl-3-(3-((1-(2-((2,2-dimethyl-3-((7-nitrobenzo[c]-[1,2,5]oxadiazol-4-yl)amino)propyl)carbamoyl)-5-methoxyphenyl)piperidin-4-yl)-methoxy)phenyl)propanoic acid (TUG-2489 - 15)**

**Step 1:** To a solution of compound **14e** (13 mg, 0.02 mmol) in DCM (300 µL) 4 M HCl in 1,4-dioxane (60 µL) was added dropwise at rt and the stirring was continued for 18 h. Upon completion, the reaction mixture was concentrated *in vacuo* and co-evaporated with DCM (×3) to give 12 mg (quant. crude) of an off-white solid that was used without further purification: $t_R$ = 9.4 min (purity 95.20% by HPLC); $^1$**H NMR** (400 MHz, $CD_3OD$) δ 8.18 (d, *J* = 8.9 Hz, 1H), 7.58 (d, *J* = 2.4 Hz, 1H), 7.26 (dd, *J* = 9.0, 2.4 Hz, 1H), 7.22 (t, *J* = 7.9 Hz, 1H), 6.94 – 6.78 (m, 3H), 4.11 – 3.95 (m, 5H), 3.86 – 3.69 (m, 4H), 3.42 (s, 2H), 2.85 (s, 2H), 2.82 – 2.64 (m, 2H), 2.46 – 2.23 (m, 4H), 2.00 – 1.83 (m, 2H), 1.12 (s, 6H), 1.11 – 0.98 (m, 1H), 0.65 – 0.54 (m, 1H), 0.46 – 0.35 (m, 1H), 0.36 – 0.25 (m, 1H), 0.20 – 0.08 (m, 1H); $^{13}$**C NMR** (151 MHz, $CD_3OD$) δ 176.2, 170.7, 165.8, 160.2, 147.4, 131.9,

130.4, 121.2, 116.5, 115.0, 113.3, 109.9, 72.1, 57.1, 56.7, 54.8, 47.7, 47.6, 42.4, 36.5, 34.6, 29.1, 28.7, 23.8, 18.3, 5.8, 4.6; **ESI-MS** m/z 538.4 ($M+H^+$).

**Step 2:** Compound from Step 1 (15 mg, 0.02 mmol) was added to a dry vial and dissolved in MeOH (0.67 mL) and $NaHCO_3$ (12 mg, 0.13 mmol) was added. The vial was then covered with aluminum foil and NBD-Cl (16 mg, 0.08 mmol) was added. The reaction mixture was stirred under Ar at 52 °C in a heating block for 4 h, in the dark. The reaction mixture was then cooled to rt, and the solvent was evaporated under $N_2$ flow. The crude material was redissolved in EtOAc and the mixture was quenched with 1 M $HCl_{aq}$. The aqueous phase was extracted with EtOAc (×3), the combined organic layers were dried over $MgSO_4$, filtered and concentrated *in vacuo*. The crude compound was purified by preparative HPLC (40 to 100% B over 20 min) and the combined fractions were lyophilized to give 5 mg (25%) of an orange solid as TFA salt: $t_R$ = 8.2 min (purity 99.82% by HPLC at 450 nm); **$^1$H NMR** (400 MHz, $CD_3OD$) δ 8.50 (d, *J* = 8.9 Hz, 1H), 8.10 (d, *J* = 9.0 Hz, 1H), 7.21 (t, *J* = 8.1 Hz, 1H), 6.89 – 6.83 (m, 2H), 6.82 – 6.75 (m, 1H), 6.50 (d, *J* = 8.9 Hz, 1H), 4.02 – 3.92 (m, 5H), 3.70 – 3.42 (m, 6H), 2.82 – 2.65 (m, 2H), 2.44 – 2.19 (m, 2H), 1.86 (br s, 2H), 1.19 – 1.03 (m, 7H), 0.66 – 0.56 (m, 1H), 0.46 – 0.37 (m, 1H), 0.35 – 0.27 (m, 1H), 0.20 – 0.12 (m, 1H); **$^{19}$F NMR** (376 MHz, $CD_3OD$) δ -77.0; **HRMS (MALDI)** calcd for $C_{37}H_{45}N_6O_8^+$ ($M+H^+$) 701.3293, found 701.3267.

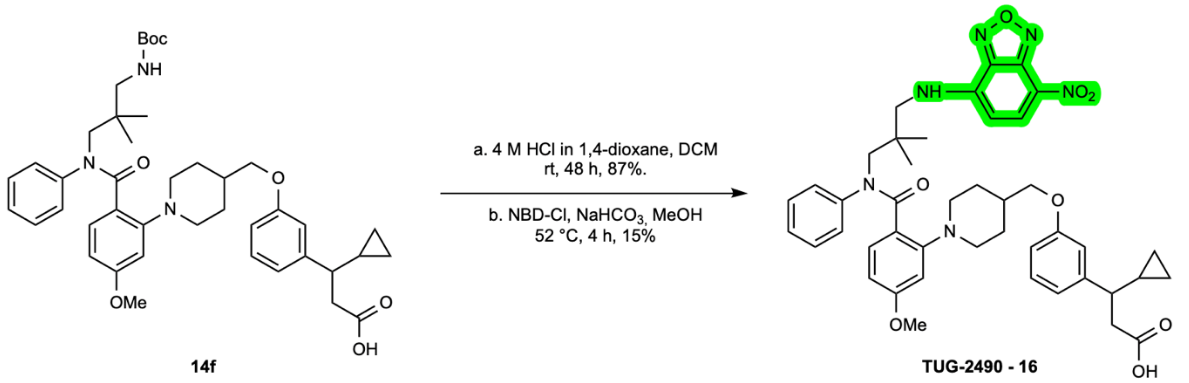


**2,2,2-Trifluoroacetic acid-3-cyclopropyl-3-(3-((1-(2-((2,2-dimethyl-3-((7-nitrobenzo[*c*]-[1,2,5]oxadiazol-4-yl)amino)propyl)(phenyl)carbamoyl)-5-methoxyphenyl)piperidin-4-yl)methoxy)phenyl)propanoic acid (TUG-2490 - 16)**

**Step 1:** To a solution of compound **14f** (18 mg, 0.03 mmol) in anh. DCM (400 µL) 4 M HCl in 1,4-dioxane (60 µL) was added dropwise at rt and the stirring was continued for 48 h. Upon completion, the reaction mixture was concentrated *in vacuo* and co-evaporated with DCM (×3) to give 15 mg (87%) as an off-white foam that was used without further purification: $^{1}$**H NMR** (600 MHz, $CD_3OD$) δ 7.49 – 7.39 (m, 2H), 7.34 – 7.15 (m, 6H), 6.90 – 6.78 (m, 3H), 6.67 (dd, *J* = 9.0, 2.2 Hz, 1H), 4.07 (s, 2H), 4.01 (d, *J* = 6.1 Hz, 1H), 3.80 (s, 3H), 3.06 (s, 2H), 2.82 – 2.64 (m, 2H), 2.37 – 2.27 (m, 1H), 2.26 – 2.14 (m, 3H), 2.09 – 1.93 (m, 2H), 1.11 – 1.02 (m, 1H), 1.00 (s, 6H), 0.64 – 0.55 (m, 1H), 0.45 – 0.37 (m, 1H), 0.34 – 0.27 (m, 1H), 0.19 – 0.11 (m, 1H); $^{13}$**C NMR** (151 MHz, $CD_3OD$) δ 176.2, 174.6, 164.0, 160.42, 160.37, 147.4, 147.2, 145.3, 130.5, 130.4, 128.7, 128.5, 121.1, 121.0, 115.0, 114.96, 113.40, 113.38, 72.5, 58.2, 56.5, 42.4, 42.3, 37.6, 35.2, 29.1, 25.0, 18.3, 18.1, 5.8, 4.6; **ESI-MS** m/z 614.4 ($M+H^+$).

**Step 2:** Compound from Step 1 (9 mg, 0.01 mmol) was added to a dry vial and dissolved in MeOH (0.35 mL) and $NaHCO_3$ (7 mg, 0.08 mmol) was added. The vial was then covered with aluminum foil and NBD-Cl (8 mg, 0.04 mmol) was added. The reaction mixture was stirred under Ar at 52 °C in a heating block for 4 h, in the dark. The reaction mixture was then cooled to rt, and the solvent was evaporated under $N_2$ flow. The crude material was redissolved in EtOAc and the mixture was quenched with 1 M $HCl_{aq}$. The aqueous phase was extracted with EtOAc (×3), the combined organic layers were dried over $MgSO_4$, filtered and concentrated *in vacuo*. The crude compound was purified by preparative HPLC (40 to 100% B over 20 min) and the combined fractions were lyophilized to give 2 mg (15%) of an orange solid as TFA salt: $t_R$ = 13.3 min (purity 98.23% by HPLC at 450 nm); $^{1}$**H NMR** (600 MHz, $CD_3OD$) δ 8.44 (d, *J* = 8.9 Hz, 1H), 7.29 – 7.02 (m, 7H), 6.83 (d, *J* = 7.6 Hz, 1H), 6.79 (s, 1H), 6.68 (d, *J* = 8.2 Hz, 1H), 6.57 – 6.39 (m, 3H), 4.14 (s, 2H), 3.84 (d, *J* = 5.6 Hz, 2H), 3.71 (s, 3H), 3.66 – 3.55 (m, 2H), 2.88 – 2.61 (m, 3H), 2.37 – 2.27 (m, 1H), 1.99 – 1.81 (m, 3H), 1.70 – 1.54 (m, 2H), 1.09 – 1.00 (m, 1H), 0.96 (s, 6H), 0.62 – 0.55 (m, 1H), 0.44 – 0.37 (m, 1H), 0.33 – 0.26 (m, 1H), 0.18 – 0.12 (m, 1H); $^{19}$**F NMR** (376 MHz, $CD_3OD$) δ -77.4; **HRMS (MALDI)** calcd for $C_{43}H_{49}N_6O_8^+$ ($M+H^+$) 777.3606, found 777.3573.

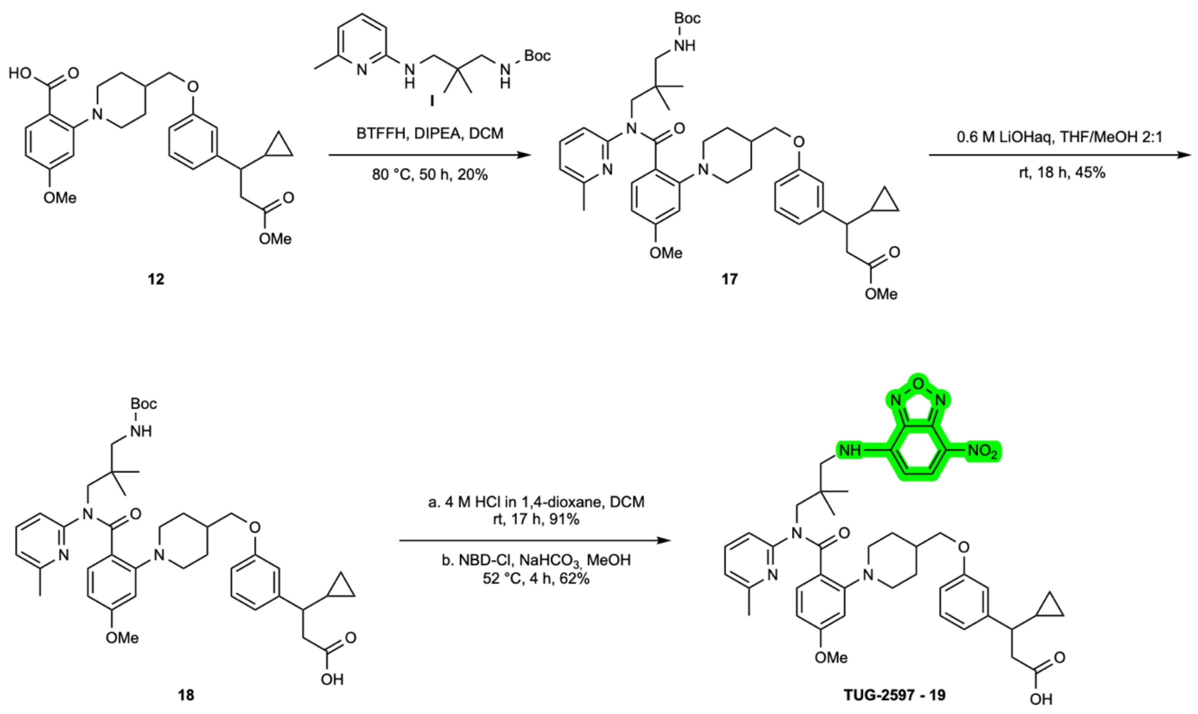


**Methyl 3-(3-((1-(2-((3-((*tert*-butoxycarbonyl)amino)-2,2-dimethylpropyl)(6-methyl-pyridin-2-yl)carbamoyl)-5-methoxyphenyl)piperidin-4-yl)methoxy)phenyl)-3-cyclo-propylpropanoate (17)**

Compound **12** (25 mg, 0.05 mmol) was added to a microwave vial and dissolved in anh. DCM (500 µL). BTFFH (41 mg, 0.13 mmol) and DIPEA (67 µL, 0.38 mmol) were added, the reaction was stirred under Ar at rt for 30 min. Then, compound **I** (43 mg, 0.14 mmol) was added, the vial was sealed, and the reaction mixture was stirred at 80 °C for 50 h. The reaction mixture was cooled to rt, diluted with $H_2O$ and extracted with EtOAc (×3). The combined organic layers were washed with brine, dried over $MgSO_4$, filtered and concentrated *in vacuo*. The crude compound was purified by automated flash column chromatography (EtOAc:*n*-heptane, 0:100→1:1) affording 14 mg (20%) of a white foam: $\boldsymbol{R_f}$ = 0.60 (EtOAc:*n*-heptane, 1:1); **$^1$H NMR** (400 MHz, $CDCl_3$) δ 7.28 (s, 1H), 7.22 (t, *J* = 7.8 Hz, 1H), 7.16 – 7.07 (m, 1H), 6.88 – 6.70 (m, 4H), 6.51 (dd, *J* = 8.5, 2.3 Hz, 1H), 6.36 – 6.06 (m, 2H), 4.47 – 4.04 (m, 2H), 3.88 – 3.81 (m, 2H), 3.77 (s, 3H), 3.62 (s, 3H), 2.90 (s, 2H), 2.82 – 2.68 (m, 2H), 2.64 – 2.52 (m, 1H), 2.47 (s, 3H), 2.35 (dt, *J* = 9.9, 7.6 Hz, 1H), 1.92 – 1.52 (m, 5H), 1.45 (s, 9H), 1.06 – 0.98 (m, 1H), 0.94 – 0.53 (m, 7H), 0.49 – 0.38 (m, 1H), 0.33 – 0.22 (m, 1H), 0.20 – 0.10 (m, 1H); **$^{13}$C NMR** (151 MHz, $CDCl_3$) δ 173.0, 171.3, 161.8, 159.2, 156.8, 155.5, 146.0, 136.4, 131.9, 129.5, 119.8, 117.7, 113.9, 112.3, 78.6,

72.6, 60.5, 55.5, 53.6, 51.6, 47.4, 41.7, 38.1, 35.9, 32.0, 29.8, 28.7, 28.5, 24.4, 22.8, 21.2, 17.2, 14.4, 14.3, 5.4, 4.2; **ESI-MS** m/z 743.7 (M+H$^+$).

**2,2,2-Trifluoroacetic acid-3-(3-((1-(2-((3-((*tert*-butoxycarbonyl)amino)-2,2-dimethyl-propyl)(6-methylpyridin-2-yl)carbamoyl)-5-methoxyphenyl)piperidin-4-yl)methoxy)-phenyl)-3-cyclopropylpropanoic acid (18)**

To a solution of compound **17** (13 mg, 0.02 mmol) dissolved in a mixture of THF/MeOH (2:1, 1 mL), 0.6 M $LiOH_{aq}$ (200 µL, 0.12 mmol) was added, and the reaction mixture was stirred at rt for 18 h. The reaction was diluted with $H_2O$, acidified with 1 M $HCl_{aq}$ and extracted with EtOAc (×3). The organic phases were combined, washed with brine, dried over $MgSO_4$, filtered and concentrated *in vacuo*. The crude material was purified by preparative HPLC (40 to 100% B over 15 min) and the combined fractions were lyophilized affording 7 mg (45%) of a white solid as TFA salt: $t_R$ = 14.1 min (purity 97.76% by HPLC); **$^1$H NMR** (400 MHz, $CD_3OD$) δ 7.42 (t, *J* = 7.8 Hz, 1H), 7.26 – 7.12 (m, 2H), 6.98 (d, *J* = 7.5 Hz, 1H), 6.90 – 6.76 (m, 4H), 6.71 – 6.60 (m, 2H), 4.16 (s, 2H), 3.92 (d, *J* = 5.8 Hz, 2H), 3.78 (s, 3H), 3.11 – 2.94 (m, 4H), 2.77 (dd, *J* = 14.9, 6.8 Hz, 1H), 2.69 (dd, *J* = 14.9, 8.3 Hz, 1H), 2.42 (s, 3H), 2.38 – 2.27 (m, 1H), 2.05 – 1.93 (m, 3H), 1.78 – 1.59 (m, 2H), 1.43 (s, 9H), 1.13 – 0.99 (m, 1H), 0.79 (s, 6H), 0.65 – 0.54 (m, 1H), 0.46 – 0.35 (m, 1H), 0.35 – 0.25 (m, 1H), 0.21 – 0.10 (m, 1H); **$^{13}$C NMR** (151 MHz, $CD_3OD$) δ 176.2, 174.0, 163.7, 160.5, 158.6, 158.4, 156.6, 147.3, 139.2, 132.8, 130.4, 122.1, 121.0, 119.6, 115.0, 113.4, 107.1, 80.0, 73.1, 56.1, 55.2, 42.4, 39.3, 36.4, 29.7, 28.8, 24.7, 24.0, 18.3, 5.8, 4.6; **HRMS (MALDI)** calcd for $C_{42}H_{57}N_4O_7^+$ (M+H$^+$) 729.4221, found 729.4220.

**2,2,2-Trifluoroacetic acid-3-cyclopropyl-3-(3-((1-(2-((2,2-dimethyl-3-((7-nitrobenzo[*c*]-[1,2,5]oxadiazol-4-yl)amino)propyl)(6-methylpyridin-2-yl)carbamoyl)-5-methoxy-phenyl)piperidin-4-yl)methoxy)phenyl)propanoic acid (TUG-2597 - 19)**

**Step 1:** Compound **18** (6 mg, 0.01 mmol) was dissolved in anh. DCM (600 µL), and 4 M HCl in dioxane (50 µL) was added. The reaction mixture was stirred at rt for 17 h. Upon completion monitored by TLC, the reaction mixture was concentrated *in vacuo* and co-evaporated with DCM (×3) to give 5 mg (91% crude) of a bright yellow solid that was used in the next step without further purification: **$^1$H NMR** (400 MHz, $CD_3OD$) δ 8.05 – 7.78 (m, 1H), 7.70 – 7.56 (m, 1H), 7.46 (s, 1H), 7.30 (d, *J* = 7.6 Hz, 1H), 7.22 (t, *J* = 8.1 Hz,

1H), 7.15 (d, *J* = 8.8 Hz, 1H), 6.91 – 6.72 (m, 4H), 4.19 (s, 2H), 4.09 – 4.00 (m, 4H), 3.90 – 3.79 (m, 5H), 3.24 (s, 2H), 2.78 (dd, *J* = 14.9, 6.7 Hz, 1H), 2.70 (dd, *J* = 14.9, 8.4 Hz, 1H), 2.49 (s, 3H), 2.42 – 2.28 (m, 4H), 2.18 – 2.03 (m, 2H), 1.11 – 1.04 (m, 1H), 1.02 (s, 6H), 0.65 – 0.54 (m, 1H), 0.46 – 0.35 (m, 1H), 0.36 – 0.26 (m, 1H), 0.21 – 0.07 (m, 1H); **ESI-MS** m/z 629.7 ($M+H^+$).

**Step 2:** Compound from Step 1 (5 mg, 0.01 mmol) was added to a dry vial and dissolved in MeOH (0.50 mL) and $NaHCO_3$ (3 mg, 0.04 mmol) was added. The vial was then covered with aluminum foil and NBD-Cl (4 mg, 0.02 mmol) was added. The reaction mixture was stirred under Ar at 52 °C for 4 h, in the dark. The reaction mixture was then cooled to rt, and the solvent was evaporated gently under $N_2$ flow. The crude material was redissolved in EtOAc and the mixture was quenched with 1 M $HCl_{aq}/H_2O$, 1:1. The aqueous phase was extracted with EtOAc (×3), the combined organic layers were dried over $MgSO_4$, filtered and concentrated *in vacuo*. The crude compound was purified by preparative HPLC (40 to 100% B over 15 min) and the combined fractions were lyophilized to give 4 mg (62%) of an orange solid TFA salt: $t_R$ = 10.3 min (purity 99.99% by HPLC at 450 nm); **$^1$H NMR** (600 MHz, $CD_3OD$) 8.48 (d, *J* = 8.8 Hz, 1H), 7.44 (s, 1H), 7.21 (d, *J* = 8.6 Hz, 1H), 7.15 (t, *J* = 7.9 Hz, 1H), 6.99 (d, *J* = 7.6 Hz, 1H), 6.90 – 6.80 (m, 2H), 6.79 – 6.76 (m, 1H), 6.70 – 6.60 (m, 3H), 6.47 (d, *J* = 8.9 Hz, 1H), 4.32 (s, 2H), 3.85 (d, *J* = 5.7 Hz, 2H), 3.78 (s, 3H), 3.62 – 3.51 (m, 2H), 3.19 – 2.92 (m, 1H), 2.75 (dd, *J* = 14.9, 6.7 Hz, 1H), 2.71 – 2.64 (m, 1H), 2.42 (s, 3H), 2.34 – 2.27 (m, 1H), 2.06 – 1.89 (m, 3H), 1.74 – 1.59 (m, 2H), 1.08 – 1.00 (m, 1H), 0.96 (s, 6H), 0.62 – 0.55 (m, 1H), 0.44 – 0.36 (m, 1H), 0.33 – 0.26 (m, 1H), 0.18 – 0.11 (m, 1H); **$^{19}$F NMR** (376 MHz, $CD_3OD$) δ -77.5; **HRMS (MALDI)** calcd for $C_{43}H_{50}N_7O_8^+$ ($M+H^+$) 792.3715, found 792.3688.

***tert*-Butyl (3-((7-nitrobenzo[*c*][1,2,5]oxadiazol-4-yl)amino)propyl)carbamate (L)**

Compound synthesized similar to Taliani *et al.*[16] but using (3-aminopropyl)carbamate. A solution of NBD-Cl (69 mg, 0.34 mmol) in anh. DMF (0.35 mL) was added dropwise to a stirred solution of *tert*-butyl (3-aminopropyl)carbamate (60 mg, 0.34 mmol) and $Et_3N$ (53 µL, 0.38 mmol) in anh. DMF (0.7 mL) under $N_2$. The dark solution was stirred at rt for 20 h, in the dark. Upon completion, the mixture was diluted with $H_2O$ and extracted with DCM (×3). The combined organic layers were dried over $MgSO_4$, filtered and concentrated *in vacuo*. The crude material was purified by automated flash chromatography (25% → 45% EtOAc in *n*-heptane) affording 74 mg (64%) of an orange-brown solid: $\boldsymbol{R_f}$ = 0.70 (EtOAc); **$^1$H NMR** (400 MHz, $CD_3OD$) δ 8.53 (d, *J* = 8.8 Hz, 1H), 6.35 (d, *J* = 8.9 Hz, 1H), 3.57 (s, 2H), 3.19 (t, *J* = 6.7 Hz, 2H), 1.93 (p, *J* = 6.8 Hz, 2H), 1.42 (s, 9H). Spectra in agreement with literature.[17]

**4-((3-(1-Cyclopropyl-3-((3-((7-nitrobenzo[*c*][1,2,5]oxadiazol-4-yl)amino)propyl)amino)-3-oxopropyl)phenoxy)methyl)-1-(5-methoxy-2-((6-methylpyridin-2-yl)(neopentyl)carbamoyl)phenyl)piperidin-1-ium 2,2,2-trifluoroacetate (TUG-2598 - 20)**

Compound synthesized similar to Robertson *et al.*[17] but with Cpd 6. To a solution of Cpd 6 (5.8 mg, 9.4 µmol) in DMF (0.5 mL) cooled to 0 °C, $Et_3N$ was slowly added (6 µL, 43 µmol), followed by EDC·HCl (2.7 mg, 14 µmol), HOBt·$H_2O$ (2.4 mg, 16 µmol) and 3-((7-nitrobenzo[*c*][1,2,5]oxadiazol-4-yl)amino)-propan-1-aminium chloride (3.9 mg, 14 µmol), that was synthesized from compound **I** as previously reported.[17,18] The reaction was allowed to reach rt and stirred for 15 h, in the dark. Upon completion, the reaction mixture was quenched with 0.5 M $HCl_{aq}$ and extracted with DCM (×4). The combined organic phases were washed with sat. aq. $Na_2CO_3$, brine, dried over $MgSO_4$, filtered and concentrated *in vacuo*. The crude material was purified by preparative HPLC affording 5.6 mg (56%) of a red solid as TFA salt: $\boldsymbol{t_R}$ = 13.9 min (purity 99.71% by HPLC at 450 nm); **$^1$H NMR** (400 MHz, $CD_3OD$) δ 8.50 (d, *J* = 8.8 Hz, 1H), 7.54 (t, *J* = 7.8 Hz, 1H), 7.21 (t, *J* = 7.9 Hz, 1H), 7.10 (d, *J* = 8.6 Hz, 1H), 7.06 – 6.96 (m, 2H), 6.93 – 6.78 (m, 3H), 6.72 (d, *J* = 8.1 Hz, 1H), 6.66 (dd, *J* = 8.7, 2.3 Hz, 1H), 6.15 (d, *J* = 8.8 Hz, 1H), 4.09 (s, 2H), 3.85 (d, *J* = 5.4 Hz, 2H), 3.80 (s, 3H), 3.44 (s, 2H), 3.23 – 3.03 (m, 4H), 2.74 – 2.54 (m, 2H), 2.40 – 2.26 (m, 4H), 2.02 – 1.92 (m, 3H), 1.82 – 1.57 (m, 4H), 1.12 – 1.00 (m, 1H), 0.88 (s, 9H), 0.63 – 0.54 (m, 1H), 0.46 – 0.35 (m, 1H), 0.34 – 0.26 (m, 1H), 0.21 – 0.12 (m, 1H); **$^{19}$F NMR** (376 MHz, $CD_3OD$) δ -77.5; **HRMS (MALDI)** calcd for $C_{46}H_{57}N_8O_7^+$ (M+H$^+$) 833.4344, found 833.4347.

**3-(4-((4'-(2-(2-((*tert*-Butoxycarbonyl)amino)ethoxy)ethoxy)-2'-methyl-[1,1'-biphenyl]-3-yl)methoxy)-2-fluorophenyl)propanoic acid (21)**

Compound **11** from Christiansen *et al.*[1] (79 mg, 0.13 mmol) was dissolved in THF (0.76 mL), MeOH (3 drops) and 0.6 M $LiOH_{aq}$ (0.63 mL, 0.38 mmol) were added. The reaction mixture was stirred for 19 h at rt. After completion, the reaction was diluted with water, acidified with 1 M $HCl_{aq}$ and extracted with EtOAc (×3). The organic layers were combined, washed with brine, dried over $MgSO_4$ and filtered. The residue was concentrated *in vacuo* to give 71 mg (95%) of a clear oil; **$^1$H NMR** (400 MHz, $CDCl_3$) δ 7.45 – 7.31 (m, 3H), 7.29 – 7.22 (m, 1H), 7.18 – 7.08 (m, 2H), 6.86 – 6.79 (m, 2H), 6.72 – 6.65 (m, 2H), 5.06 (s, 2H), 4.18 – 4.12 (m, 2H), 3.87 – 3.79 (m, 2H), 3.63 (t, *J* = 5.2 Hz, 2H), 3.35 (br s, 2H), 2.91 (t, *J* = 7.7 Hz, 2H), 2.64 (t, *J* = 7.7 Hz, 2H), 2.23 (s, 3H), 1.45 (s, 9H); **$^{13}$C NMR** (101 MHz, $CDCl_3$) δ 177.8, 161.6 (d, *J* = 245.4 Hz), 158.8 (d, *J* = 11.1 Hz), 158.1, 142.1, 136.9, 136.5, 134.6, 131.0, 130.9, 129.3, 128.6, 128.5, 125.7, 119.4 (d, *J* = 16.2 Hz), 116.7, 112.0, 110.8 (d, *J* = 3.1 Hz), 102.9 (d, *J* = 25.8 Hz), 70.54, 70.46, 69.7, 67.5, 34.4, 28.5, 23.9 (d, *J* = 2.2 Hz), 20.8; **ESI-MS** m/z 566.3 ($M-H^+$).

**2-(2-((3'-((4-(2-Carboxyethyl)-3-fluorophenoxy)methyl)-2-methyl-[1,1'-biphenyl]-4-yl)oxy)ethoxy)ethan-1-aminium chloride (22)**

To a solution of compound **21** (71 mg, 0.13 mmol) in DCM (0.30 mL) 4 M HCl in 1,4-dioxane (0.26 mL, 1.04 mmol) was added dropwise. The resulting mixture was stirred for 2 h at rt. After completion, the solvent was evaporated *in vacuo* to afford 61 mg (97%) of a white solid that was used in the next step without further purification: $t_R$ = 13.4 min (purity = 98.13% by HPLC); **$^1$H NMR** (400 MHz, $CD_3OD$) δ 7.48 – 7.36 (m, 2H), 7.36 – 7.31 (m, 1H), 7.28 – 7.10 (m, 3H), 6.91 – 6.81 (m, 2H), 6.81 – 6.71 (m, 2H), 5.13 (s, 2H), 4.26 – 4.19 (m, 2H), 3.97 – 3.90 (m, 2H), 3.87 – 3.77 (m, 2H), 3.23 – 3.16 (m, 2H), 2.89 (t, *J* = 7.7 Hz, 2H), 2.57 (t, *J* = 7.7 Hz, 2H), 2.21 (s, 3H); **$^{13}$C NMR** (101 MHz, $CD_3OD$) δ 176.5, 167.1, 162.8 (d, *J* = 243.9 Hz), 160.1 (d, *J* = 10.9 Hz), 159.3, 143.3, 138.3, 137.9, 136.1, 132.0 (d, *J* = 6.8 Hz), 131.8, 130.0, 129.5, 129.4, 126. 8, 120.8 (d, *J* = 16.1 Hz), 117.4, 112.9, 111.9 (d, *J* = 3.1 Hz), 103.5 (d, *J* = 25.9 Hz), 71.2, 71.0, 68.6, 68.1, 40.8, 35.5, 24.9 (d, *J* = 2.3 Hz), 20.8; **ESI-MS** m/z 466.2 ($M-H^+$).

**3-(6-((2-(2-((3'-((4-(2-Carboxyethyl)-3-fluorophenoxy)methyl)-2-methyl-[1,1'-biphenyl]-4-yl)oxy)ethoxy)ethyl)amino)-6-oxohexyl)-1,1-dimethyl-2-((1*E*,3*E*)-3-**

**(1,1,3-trimethyl-1,3-dihydro-2*H*-benzo[*e*]indol-2-ylidene)prop-1-en-1-yl)-1*H*-benzo[*e*]indol-3-ium 2,2,2-trifluoroacetate (TUG-2287 - 23)**

Cy3.5 NHS ester (2.0 mg, 2.7 µmol) was dissolved in anh. DMF (200 µL). Anh. DIPEA (1.4 µL, 8 µmol) and compound **22** (1.5 mg, 3 µmol) were added. The vial was flushed with Ar and capped. The mixture was stirred at rt for 45 h, whereupon it was diluted with MeCN (500 µL) and Milli-Q water (100 µL) and purified by preparative HPLC (50 to 100% B over 23 min). The corresponding fractions were combined and concentrated *in vacuo* to give 2.0 mg (66%) of the title compound as a dark purple solid TFA salt: $t_R$ = 13.8 min (purity 99.99% by HPLC at 579 nm); **$^1$H NMR** (400 MHz, $CD_3OD$) δ 8.72 (t, *J* = 13.5 Hz, 1H), 8.26 (t, *J* = 8.2 Hz, 2H), 8.07 – 7.98 (m, 4H), 7.70 – 7.63 (m, 2H), 7.62 (d, *J* = 8.8 Hz, 1H), 7.57 (d, *J* = 8.8 Hz, 1H), 7.54 – 7.48 (m, 2H), 7.33 – 7.26 (m, 2H), 7.17 (s, 1H), 7.13 (t, *J* = 8.6 Hz, 1H), 7.10 – 7.05 (m, 1H), 6.99 (d, *J* = 8.4 Hz, 1H), 6.81 – 6.77 (m, 1H), 6.77 – 6.72 (m, 1H), 6.71 – 6.64 (m, 2H), 6.49 – 6.40 (m, 2H), 4.98 (s, 2H), 4.19 (t, *J* = 7.7 Hz, 2H), 4.13 – 4.08 (m, 2H), 3.81 – 3.77 (m, 5H), 3.60 (t, *J* = 5.4 Hz, 2H), 3.39 (t, *J* = 5.4 Hz, 2H), 2.84 (t, *J* = 7.7 Hz, 2H), 2.53 (t, *J* = 7.7 Hz, 2H), 2.24 (t, *J* = 7.1 Hz, 2H), 2.08 (s, 3H), 2.08 – 2.01 (m, 12H), 1.89 (p, *J* = 8.2 Hz, 2H), 1.74 (p, *J* = 7.3 Hz, 2H), 1.57 – 1.49 (m, 2H); **$^{19}$F NMR** (376 MHz, $CD_3OD$) δ -77.0, -118.6; **HRMS (MALDI)** calcd for $C_{65}H_{68}FN_3O_6$ (M+H$^+$) 1006.5164, found 1006.5159.

**2-((1*E*,3*E*)-5-((*Z*)-1-(6-((2-(2-((3'-((4-(2-Carboxyethyl)-3-fluorophenoxy)methyl)-2-methyl-[1,1'-biphenyl]-4-yl)oxy)ethoxy)ethyl)amino)-6-oxohexyl)-3,3-dimethylindolin-2-ylidene)penta-1,3-dien-1-yl)-1,3,3-trimethyl-3*H*-indol-1-ium-5-sulfonate (TUG-2355 - 24)**

**Step 1**: A dry flask was charged with 2-((1*E*,3*E*)-5-((*Z*)-1-(5-carboxypentyl)-3,3-dimethylindolin-2-ylidene)penta-1,3-dien-1-yl)-1,3,3-trimethyl-3*H*-indol-1-ium-5-sulfonate[19] (5 mg, 9 µmol), anh. DMF (200 µL), anh. DIPEA (8 µL, 46 µmol), and PyBOP (6 mg, 11 µmol) under Ar. The reaction mixture was stirred at 0 °C for 1 h in the dark. Then, methyl 3-(4-((4'-(2-(2-aminoethoxy)ethoxy)-2'-methyl-[1,1'-biphenyl]-3-yl)methoxy)-2-fluorophenyl)propanoate (7 mg, 13 µmol) was added and the mixture was stirred at rt for 18 h in the dark. After completion, the reaction mixture was added MeCN (0.50 mL), Milli-Q water (0.10 mL) and purified by preparative HPLC (40 to 100% B over 25 min). The corresponding fractions were combined and concentrated *in vacuo* to give 5.3 mg (58%)

of a blue solid: $t_R$ = 14.0 min (purity 99.99% by HPLC at 595 nm); $^1$H NMR (600 MHz, $CD_3OD$) δ 8.32 – 8.17 (m, 2H), 7.88 – 7.83 (m, 2H), 7.51 – 7.47 (m, 1H), 7.42 – 7.33 (m, 3H), 7.32 – 7.21 (m, 4H), 7.17 – 7.10 (m, 2H), 7.05 (d, *J* = 8.3 Hz, 1H), 6.82 (d, *J* = 2.6 Hz, 1H), 6.80 – 6.77 (m, 1H), 6.75 – 6.68 (m, 2H), 6.61 (t, *J* = 12.4 Hz, 1H), 6.32 (d, *J* = 13.8 Hz, 1H), 6.18 (d, *J* = 13.5 Hz, 1H), 5.07 (s, 2H), 4.15 – 4.12 (m, 2H), 4.07 (t, *J* = 7.5 Hz, 2H), 3.84 – 3.80 (m, 2H), 3.63 – 3.60 (m, 5H), 3.54 (s, 3H), 3.39 (t, *J* = 5.4 Hz, 2H), 2.86 (t, *J* = 7.6 Hz, 2H), 2.57 (t, *J* = 7.6 Hz, 2H), 2.21 (t, *J* = 7.2 Hz, 2H), 2.14 (s, 3H), 1.83 – 1.76 (m, 2H), 1.73 – 1.65 (m, 14H), 1.49 – 1.42 (m, 2H).

**Step 2:** To a flask charged with compound from Step 1 (5.0 mg, 5 μmol) was added 4 M $HCl_{aq}$ (24 μL) and THF (250 μL). The reaction mixture was stirred at 50 °C for 18 h in the dark and monitored by HPLC. After completion, the reaction mixture was cooled to rt, added MeCN (500 μL) and Milli-Q $H_2O$ (100 μL) and purified by preparative HPLC (40 to 100% B over 25 min). The corresponding fractions were combined and concentrated *in vacuo* to give 1.18 mg (24%) of the title compound as a blue solid: $t_R$ = 12.0 min (purity 99.20% by HPLC at 650 nm); **$^1$H NMR** (600 MHz, $CD_3OD$) δ 8.30 – 8.18 (m, 2H), 7.88 – 7.84 (m, 2H), 7.52 – 7.47 (m, 1H), 7.42 – 7.34 (m, 3H), 7.32 – 7.22 (m, 4H), 7.18 – 7.12 (m, 2H), 7.06 (d, *J* = 8.3 Hz, 1H), 6.84 – 6.78 (m, 2H), 6.75 – 6.68 (m, 2H), 6.60 (t, *J* = 12.3 Hz, 1H), 6.32 (d, *J* = 13.8 Hz, 1H), 6.18 (d, *J* = 13.4 Hz, 1H), 5.07 (s, 2H), 4.17 – 4.12 (m, 2H), 4.07 (t, *J* = 7.6 Hz, 2H), 3.85 – 3.80 (m, 2H), 3.62 (t, *J* = 5.4 Hz, 2H), 3.54 (s, 3H), 3.39 (t, *J* = 5.3 Hz, 2H), 2.85 (t, *J* = 7.7 Hz, 2H), 2.52 (t, *J* = 7.6 Hz, 2H), 2.24 – 2.15 (m, 2H), 2.14 (s, 3H), 1.84 – 1.76 (m, 2H), 1.75 – 1.65 (m, 14H), 1.49 – 1.41 (m, 2H); **HRMS (MALDI)** calcd for $C_{59}H_{66}FN_3O_9S$ (M+H$^+$) 1012.4576, found 1012.4563.

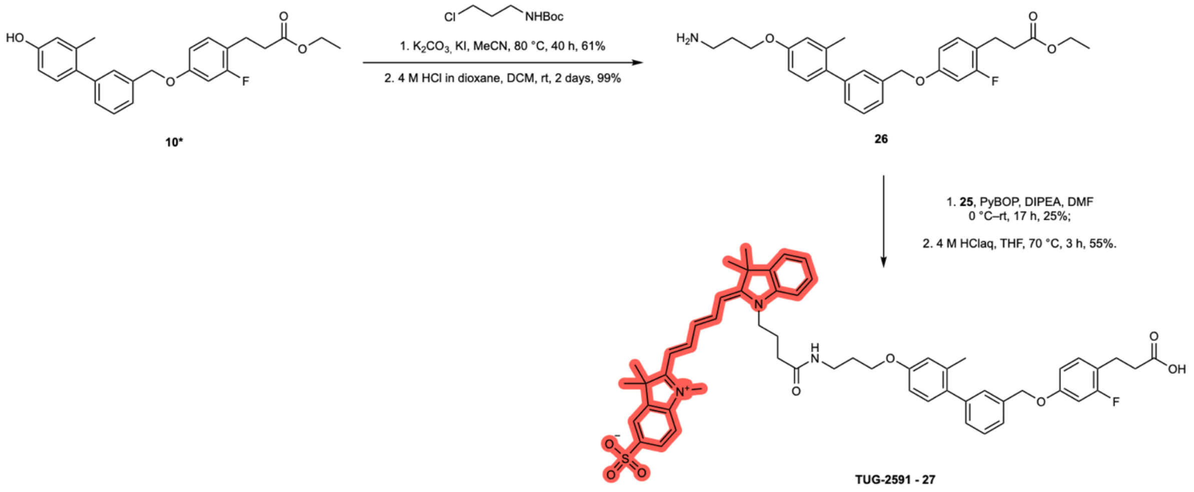

**(*E*)-2-((2*E*,4*E*)-5-(1-(3-Carboxypropyl)-3,3-dimethyl-3*H*-indol-1-ium-2-yl)penta-2,4-dien-1-ylidene)-1,3,3-trimethylindoline-5-sulfonate (25)**

Compound synthesized similar to Bunschoten *et al.*[19] but using 1-(3-carboxypropyl)-2,3,3-trimethyl-3*H*-indol-1-ium bromide[20] (31 mg, 0.09 mmol) and 1,3,3-trimethyl-2-((1*E*,3*E*)-4-(*N*-phenylacetamido)buta-1,3-dien-1-yl)-3*H*-indol-1-ium-5-sulfonate (40 mg, 0.09 mmol). The crude mixture was purified by preparative HPLC to give 5 mg (10%) of the title compound as a blue solid. **$^1$H NMR** (400 MHz, $CDCl_3$) δ: 8.38 – 8.21 (m, 2H), 7.90 – 7.84 (m, 2H), 7.53 (d, *J* = 7.5 Hz, 1H), 7.47 – 7.40 (m, 2H), 7.34 – 7.25 (m, 2H), 6.65 (t, *J* = 12.4 Hz, 1H), 6.47 (d, *J* = 13.8 Hz, 1H), 6.23 (d, *J* = 13.5 Hz, 1H), 4.24 – 4.15 (m, 2H), 3.59 (s, 3H), 2.53 (t, *J* = 6.2 Hz, 2H), 2.12 – 2.02 (m, 2H), 1.74 (s, 12H); **HRMS (ESI)** calcd for $C_{30}H_{35}N_2O_5S$ (M+H$^+$) 535.2261 found 535.2285.

**3-((3'-((4-(3-Ethoxy-3-oxopropyl)-3-fluorophenoxy)methyl)-2-methyl-[1,1'-biphenyl]-4-yl)oxy)propan-1-aminium chloride (26)**

**Step 1:** A dry microwave vial was charged with *tert*-butyl (3-chloropropyl)carbamate[21] (71 mg, 0.37 mmol), anh. MeCN (1.0 mL), KI (24 mg, 0.14 mmol), anh. $K_2CO_3$ (36 mg, 0.26 mmol), and compound **10**[18] (50 mg, 0.12 mmol) under Ar. The mixture was heated to 80 °C and stirred for 40 h. After completion, the mixture was cooled to rt, $H_2O$ was added, and the mixture was extracted with EtOAc (×3), washed with brine, dried over $MgSO_4$, filtered, and concentrated *in vacuo*. The residue was purified by flash column chromatography (0-20% EtOAc in *n*-heptane) to give 42 mg (61%) of a pale yellow oil: $\boldsymbol{R_f}$ = 0.35 (EtOAc:*n*-heptane, 2:1); **$^1$H NMR** (400 MHz, $CDCl_3$) δ 7.45 – 7.31 (m, 3H), 7.30 – 7.22 (m, 2H), 7.18 – 7.05 (m, 2H), 6.86 – 6.73 (m, 2H), 6.76 – 6.63 (m, 2H), 5.06 (s, 2H), 4.76 (br s, 1H), 4.12 (q, *J* = 7.2 Hz, 2H), 4.05 (t, *J* = 6.0 Hz, 2H), 3.37 – 3.30 (m, 2H), 2.90 (t, *J* = 7.7 Hz, 2H), 2.62 – 2.54 (m, 2H), 2.23 (s, 3H), 2.00 (p, *J* = 6.3 Hz, 2H), 1.45 (s, 9H), 1.23 (t, *J* = 7.1 Hz, 3H); **$^{13}$C NMR** (151 MHz, $CDCl_3$) δ 172.9, 161.6 (d, *J* = 245.2 Hz), 158.7 (d, *J* = 10.9 Hz), 158.2, 156.2, 142.1, 136.9, 136.5, 134.5, 131.02, 130.98, 129.3, 128.6, 128.5, 125.7, 119.7 (d, *J* = 16.2 Hz), 116.5, 111.8, 110.7 (d, *J* = 3.2 Hz), 102.8 (d, *J* = 25.9 Hz), 79.4, 70.5, 66.0, 60.6, 38.3, 34.9, 29.7, 28.6, 24.2 (d, *J* = 2.2 Hz), 20.8, 14.3; **ESI-MS** *m/z* 466.2 (M-Boc+H$^+$).

**Step 2:** A vial charged with compound from Step 1 (31 mg, 0.05 mmol) was dissolved in anh. DCM (0.11 mL) and 4 M HCl in 1,4-dioxane (0.12 mL) was added. The mixture was stirred at rt for 24 h, then additional 4 M HCl in 1,4-dioxane (0.24 mL) was added and the reaction was stirred for at rt 22 h. After completion, the mixture was concentrated *in vacuo*, co-evaporated with DCM (×3) to afford 27 mg (99%) of the title compound as a pale yellow sticky solid that was used directly in the next step; **ESI-MS** *m/z* 466.2 ($M+H^+$).

**2-((1*E*,3*E*)-5-((*Z*)-1-(4-((3-((3'-((4-(2-Carboxyethyl)-3-fluorophenoxy)methyl)-2-methyl-[1,1'-biphenyl]-4-yl)oxy)propyl)amino)-4-oxobutyl)-3,3-dimethylindolin-2-ylidene)penta-1,3-dien-1-yl)-1,3,3-trimethyl-3*H*-indol-1-ium-5-sulfonate (TUG-2591 - 27)**

**Step 1:** A dry 5 mL round bottom flask was charged with compound **25** (9 mg, 0.02 mmol), PyBOP (12 mg, 0.02 mmol), anh. DMF (0.15 mL), anh. DIPEA (16 µL, 0.06 mmol) under Ar. The mixture was stirred for 1 h at 0 °C. Then, compound **26** (11 mg, 0.02 mmol) was dissolved in anh. DMF (0.15 mL) and added to the reaction. The mixture was stirred at rt for 16 h. After completion, the mixture was diluted with Milli-Q $H_2O$, MeCN, and MeOH (≈2 mL), filtered, and purified by preparative HPLC (50 to 100% B over 25 min). The combined fractions were lyophilized to give 4.1 mg (25%) of a purple solid: $t_R$ = 9.2 min (purity 99.99% by HPLC at 650 nm); **ESI-MS** *m/z* 982.3 ($M+H^+$).

**Step 2:** To an 8 mL vial charged with compound from Step 1 (4.1 mg, 4.2 µmol) was added THF (0.50 mL), 4 M $HCl_{aq}$ (0.10 mL). The mixture was heated to 70 °C for 3 h. After completion, the mixture was cooled to rt, and concentrated *in vacuo*. The residue was purified by preparative HPLC (50 to 100% B over 10 min, flow rate 17 mL/min) and the combined fractions were lyophilized to give 2.2 mg (55%) of the title compound as a purple solid: $t_R$ = 7.4 min (purity 99.33% by HPLC at 650 nm); **$^1$H NMR** (600 MHz, $CD_3OD$) δ 8.33 – 8.21 (m, 2H), 8.05 (t, *J* = 5.7 Hz, 1H), 7.88 – 7.83 (m, 2H), 7.51 – 7.47 (m, 1H), 7.44 – 7.38 (m, 1H), 7.38 – 7.31 (m, 3H), 7.31 – 7.26 (m, 1H), 7.25 – 7.20 (m, 2H), 7.16 (t, *J* = 8.6 Hz, 1H), 7.13 – 7.08 (m, 1H), 7.02 (d, *J* = 8.3 Hz, 1H), 6.83 – 6.80 (m, 1H), 6.79 – 6.76 (m, 1H), 6.75 – 6.70 (m, 2H), 6.68 – 6.59 (m, 1H), 6.38 (d, *J* = 13.8 Hz, 1H), 6.20 (d, *J* = 13.5 Hz, 1H), 5.06 (s, 2H), 4.06 (t, *J* = 5.9 Hz, 3H), 3.53 (s, 3H), 3.48 – 3.41 (m, 2H), 2.86 (t, *J* = 7.7 Hz, 2H), 2.57 – 2.52 (m, 2H), 2.40 (t, *J* = 6.6 Hz, 2H), 2.12 (s, 3H), 2.10 – 2.05 (m, 2H), 2.01 – 1.97 (m, 2H), 1.71 (s, 6H), 1.68 (s, 6H); **$^{19}$F NMR** (376 MHz, $CD_3OD$) δ 118.6; **HRMS (MALDI)** calcd for $C_{56}H_{60}FN_3O_8S$ ($M+H^+$) 954.4157, found 954.4183.

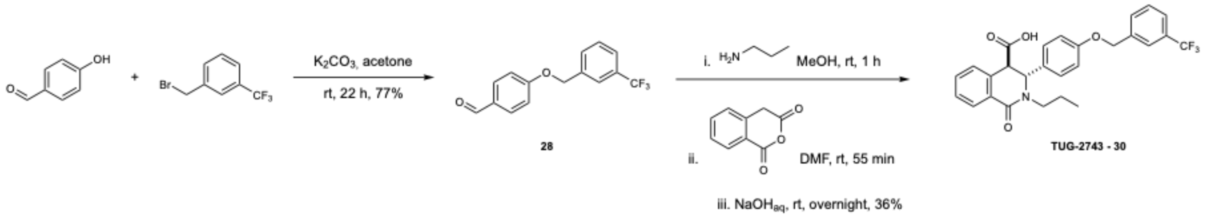

**trans-1-Oxo-2-propyl-3-(4-((3-(trifluoromethyl)benzyl)oxy)phenyl)-1,2,3,4-tetrahydro-isoquinoline-4-carboxylic acid (TUG-2743 - 29)**

Compound synthesized similarly to Humphries *et al.*[22] Compound **28**[23] was added to an 8 mL vial and dissolved in MeOH (720 µL), followed by addition of propylamine (29 µL, 0.36 mmol). The vial was closed with a lid and the resulting colorless solution was stirred at rt for 1 h. After this time, homophthalic anhydride (62 mg, 0.38 mmol) dissolved in anh. DMF (720 µL) was slowly added to the reaction mixture and the reaction was stirred at rt for 1 h. Then, 1 M $NaOH_{aq}$ (720 µL) was added and the stirring was continued at rt for 19 h. The reaction was then quenched by addition of $H_2O$, diluted with EtOAc and 1 M $HCl_{aq}$ was added. The reaction mixture was extracted with EtOAc (×3), the organic layers were combined, washed with brine, dried over $MgSO_4$, filtered and concentrated *in vacuo*. The crude material was purified by flash column chromatography and followed by preparative HPLC (20-100% B over 15 min) to give 63 mg (36%) of an off-white solid: $\boldsymbol{t_R}$ = 9.87 min (purity 99.99% by HPLC); **$^1$H NMR** (600 MHz, $CDCl_3$) δ 8.19 – 8.15 (m, 1H), 7.63 (s, 1H), 7.56 (t, *J* = 8.9 Hz, 2H), 7.48 (t, *J* = 7.7 Hz, 1H), 7.45 – 7.37 (m, 2H), 7.16 – 7.10 (m, 1H), 6.98 (d, *J* = 8.7 Hz, 2H), 6.82 (d, *J* = 8.7 Hz, 2H), 5.24 (s, 1H), 5.01 (s, 2H), 4.01 (ddd, *J* = 13.5, 9.1, 7.0 Hz, 1H), 3.91 (s, 1H), 2.83 (ddd, *J* = 14.0, 9.0, 5.5 Hz, 1H), 1.62 (ddq, *J* = 13.9, 8.9, 7.0 Hz, 2H), 0.87 (t, *J* = 7.4 Hz, 3H); **$^{13}$C NMR** (151 MHz, $CDCl_3$) δ 174.4, 163.9, 158.3, 137.9, 132.2, 131.6, 131.5, 131.2 (q, *J* = 32.4 Hz), 130.7, 129.5, 129.2, 128.8, 128.2, 127.7, 125.0 (q, *J* = 3.9 Hz), 124.2 (q, *J* = 3.9 Hz), 124.1 (q, *J* = 272.3 Hz), 115.2, 69.4, 60.7, 51.3, 48.6, 21.0, 11.5; **$^{19}$F NMR** (376 MHz, $CDCl_3$) δ -62.7; **HRMS** calcd for $C_{27}H_{25}F_3NO_4^+$ ($M+H^+$) 484.1730, found 484.1717.

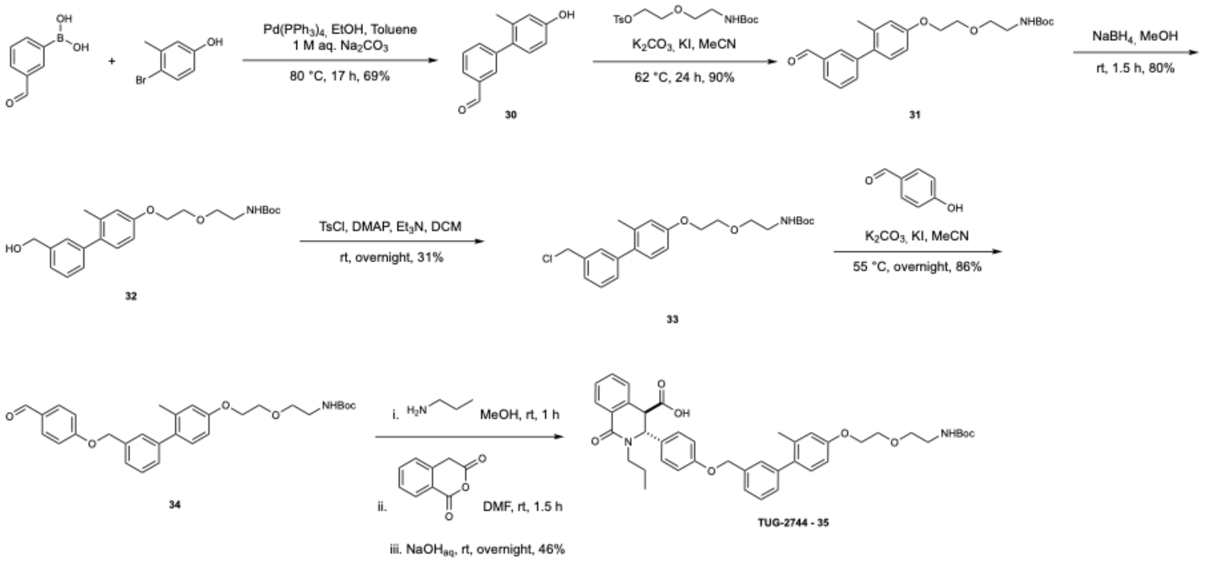

**tert-Butyl (2-(2-((3'-formyl-2-methyl-[1,1'-biphenyl]-4-yl)oxy)ethoxy)ethyl)carbamate (31)**

Compound synthesized similarly to Rexen Ulven *et al.*[24] Compound **30**[25] (60 mg, 0.28 mmol) and 2-(2-((*tert*-butoxycarbonyl)amino)ethoxy)-ethyl 4-methyl-benzenesulfonate (150 mg, 0.42 mmol) were dissolved in MeCN (1.7 mL), then $K_2CO_3$ (78 mg, 0.57 mmol) and a catalytic amount of KI were added. The resulting pale-yellow suspension was stirred under Ar at 62 °C in a pre-heated heating block for 24 h. The reaction mixture was then cooled to rt, quenched by addition of $H_2O$ and extracted with EtOAc (×3). The organic layers were combined, dried over $MgSO_4$, filtered and concentrated *in vacuo*. The crude material was purified by automated flash column chromatography (EtOAc:*n*-heptane, 1:5) to give 102 mg (90%) of a colorless oil: $R_f$ = 0.50 (EtOAc:*n*-heptane, 1:1); **$^1$H NMR** (400 MHz, $CDCl_3$) δ 10.06 (s, 1H), 7.88 – 7.78 (m, 2H), 7.60 – 7.54 (m, 2H), 7.16 (d, *J* = 8.3 Hz, 1H), 6.91 – 6.79 (m, 2H), 4.97 (s, 1H), 4.19 – 4.12 (m, 2H), 3.89 – 3.81 (m, 2H), 3.63 (t, *J* = 5.1 Hz, 2H), 3.36 (q, *J* = 5.4 Hz, 2H), 2.25 (s, 3H), 1.45 (s, 9H); **$^{13}$C NMR** (151 MHz, $CDCl_3$) δ 192.5, 158.5, 156.1, 142.7, 136.9, 136.5, 135.6, 133.5, 130.9, 130.7, 128.9, 128.1, 116.8, 112.2, 79.4, 70.6, 69.6, 67.5, 40.5, 28.6, 20.8.

**tert-Butyl (2-(2-((3'-(chloromethyl)-2-methyl-[1,1'-biphenyl]-4-yl)oxy)ethoxy)ethyl)-carbamate (33)**

To a solution of compound **32** (60 mg, 0.15 mmol) in anh. DCM (1.00 mL), tosyl chloride (37 mg, 0.20 mmol), DMAP (2.00 mg, 0.02 mmol) and $Et_3N$ (52 µL, 0.38 mmol) were added. The reaction mixture was stirred at rt under Ar overnight. The reaction was quenched with sat. aq. $NaHCO_3$ and extracted with DCM (×3). The organic layers were combined, dried over $MgSO_4$ and concentrated *in vacuo*. The crude material was purified by automated flash column chromatography (EtOAc:*n*-heptane, 1:5) to give 20 mg (31%) of a white sticky solid: **$^1$H NMR** (600 MHz, $CDCl_3$) δ 7.39 (t, *J* = 7.5 Hz, 1H), 7.36 – 7.31 (m, 2H), 7.26 – 7.24 (m, 1H), 7.14 (d, *J* = 8.3 Hz, 1H), 6.85 (d, *J* = 2.6 Hz, 1H), 6.81 (dd, *J* = 8.4, 2.7 Hz, 1H), 4.98 (s, 1H), 4.63 (s, 2H), 4.17 – 4.13 (m, 2H), 3.86 – 3.82 (m, 2H), 3.63 (t, *J* = 5.2 Hz, 2H), 3.36 (q, *J* = 5.5 Hz, 2H), 2.25 (s, 3H), 1.45 (s, 9H); **$^{13}$C NMR** (151 MHz, $CDCl_3$) δ 158.2, 156.2, 142.3, 137.4, 136.9, 134.4, 130.9, 129.8, 129.6, 128.6, 126.8, 116.7, 112.0, 79.4, 70.6, 69.7, 67.5, 46.4, 40.5, 28.6, 20.9; **ESI-MS** m/z 320.3 $(M+H-Boc)^+$.

**_tert_-Butyl (2-(2-((3'-((4-formylphenoxy)methyl)-2-methyl-[1,1'-biphenyl]-4-yl)oxy)-ethoxy)ethyl)carbamate (34)**

Compound synthesized similarly to Rexen Ulven *et al.*[24] To a dry vial compound **33** (15 mg, 36 µmol) was added and dissolved in MeCN (1.00 mL). Then, 4-hydrozybenzaldehyde (4 mg, 36 µmol), $K_2CO_3$ (10 mg, 71 µmol) and catalytic amount of KI were added, the vial was closed with a lid and the reaction mixture was stirred overnight at 55 °C in a pre-heated heating block. Upon completion, the reaction mixture was cooled to rt, quenched by addition of $H_2O$ and extracted with EtOAc (×3). The organic layers were combined, washed with brine, dried over $MgSO_4$, filtered and concentrated *in vacuo*. The crude material was purified by flash column chromatography (EtOAc:*n*-heptane, 1:3) to give 15 mg (86%) of a white solid: **$^1$H NMR** (400 MHz, $CDCl_3$) δ 9.89 (s, 1H), 7.88 – 7.80 (m, 2H), 7.47 – 7.34 (m, 3H), 7.32 – 7.25 (m, 1H), 7.15 (d, *J* = 8.3 Hz, 1H), 7.13 – 7.06 (m, 2H), 6.85 (d, *J* = 2.6 Hz, 1H), 6.81 (dd, *J* = 8.3, 2.7 Hz, 1H), 5.19 (s, 2H), 4.15 (dd, *J* = 5.7, 3.8 Hz, 2H), 3.84 (dd, *J* = 5.7, 3.8 Hz, 2H), 3.63 (t, *J* = 5.2 Hz, 2H), 3.40 – 3.31 (m, 2H), 2.23 (s, 3H), 1.45 (s, 9H); **$^{13}$C NMR** (151 MHz, $CDCl_3$) δ 190.9, 163.9, 158.2, 156.1, 142.3, 136.9, 135.9, 134.5, 132.1, 130.9, 130.3, 129.5, 128.64, 128.62, 125.8, 116.7, 115.3, 112.0, 79.4, 70.6, 70.4, 69.7, 67.5, 40.5, 28.6, 20.9; **ESI-MS** m/z 406.4 $(M+H-Boc)^+$.

**_trans_-3-(4-((4'-(2-(2-((_tert_-Butoxycarbonyl)amino)ethoxy)ethoxy)-2'-methyl-[1,1'-biphenyl]-3-yl)methoxy)phenyl)-1-oxo-2-propyl-1,2,3,4-tetrahydroisoquinoline-4-carboxylic acid (TUG-2744 - 35)**

Compound synthesized similarly to Humphries *et al.*[22] Compound **34** (15 mg, 30 µmol) was added to a vial and dissolved in MeOH (120 µL), followed by addition of propylamine

(3 µL, 36 µmol), the vial was closed with a lid and the solution was stirred at rt for 1 h. After this time, homophthalic anhydride (6 mg, 37 µmol) dissolved in anh. DMF (120 µL) was added slowly to the reaction mixture. The reaction was stirred at rt for 1 h before addition of 1 M $NaOH_{aq}$ (120 µL). The vial was closed with a lid again and stirred at rt for 18 h. The reaction was quenched by addition of $H_2O$, acidified to pH 1 by addition of 1 M $HCl_{aq}$ and extracted with EtOAc (×3). The organic layers were combined, washed with brine, dried over $MgSO_4$, filtered, and concentrated *in vacuo*. The crude material was purified by flash column chromatography followed by preparative HPLC (20-100% B over 15 min). After preparative HPLC app. 5% of Boc-deprotected product was observed, so the compound was dissolved in EtOAc and washed with sat. aq. $NH_4Cl$ and 1 M $HCl_{aq}$ (×3). The organic phase was washed with brine, dried over $MgSO_4$, filtered and concentrated *in vacuo* to yield 9.6 mg (46%) of the product as a white solid: $\boldsymbol{t_R}$ = 11.54 min (purity 99.99% by HPLC); **$^1$H NMR** (600 MHz, $CDCl_3$) δ 8.19 – 8.14 (m, 1H), 7.45 – 7.35 (m, 3H), 7.31 (d, *J* = 7.7 Hz, 1H), 7.27 (s, 1H), 7.23 (dt, *J* = 7.6, 1.5 Hz, 1H), 7.15 – 7.09 (m, 2H), 6.97 (d, *J* = 8.7 Hz, 2H), 6.85 – 6.77 (m, 4H), 5.24 (s, 1H), 5.05 – 4.94 (m, 3H), 4.16 – 4.11 (m, 2H), 4.01 (dt, *J* = 13.4, 7.7 Hz, 1H), 3.89 (s, 1H), 3.83 (t, *J* = 4.8 Hz, 2H), 3.62 (t, *J* = 5.3 Hz, 2H), 3.40 – 3.28 (m, 2H), 2.82 (ddd, *J* = 14.1, 9.0, 5.7 Hz, 1H), 2.18 (s, 3H), 1.70 – 1.58 (m, 2H), 1.44 (s, 9H), 0.88 (t, *J* = 7.4 Hz, 3H); **$^{13}$C NMR** (151 MHz, $CDCl_3$) δ 163.7, 158.5, 158.0, 156.1, 141.9, 136.7, 136.5, 134.5, 132.0, 131.7, 131.1, 130.8, 129.4, 129.3, 129.1, 128.6, 128.5, 128.3, 128.0, 127.4, 125.6, 116.5, 115.1, 111.8, 79.3, 70.4, 70.0, 69.5, 67.4, 60.7, 51.2, 48.4, 40.4, 28.4, 20.9, 20.7, 11.4; **HRMS** calcd for $C_{42}H_{49}N_2O_8^+$ (M+H $^+$) 709.3483, found 709.3468.

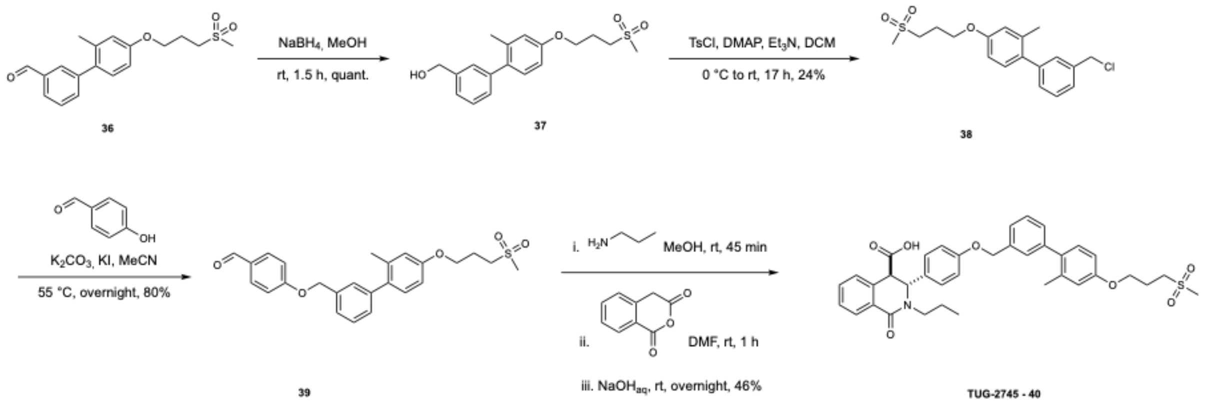


**(2'-Methyl-4'-(3-(methylsulfonyl)propoxy)-[1,1'-biphenyl]-3-yl)methanol (37)**

Compound **36**[25] (69 mg, 0.21 mmol) was charged in a vial flushed with Ar and dissolved in MeOH (2.00 mL). To this solution, $NaBH_4$ (8.00 mg, 0.21 mmol) was added, and evolution of bubbles was observed. The reaction mixture was stirred at rt under Ar for 1.5 h. Upon completion, the reaction was quenched by addition of $H_2O$ and extracted with EtOAc (×3). The combined organic layers were washed with brine, dried over $MgSO_4$, filtered and concentrated *in vacuo* to provide 69 mg (quant. crude yield) of the product as a white solid that was used in the next reaction without further purification: **$^1$H NMR** (600 MHz, $CDCl_3$) δ 7.39 (t, *J* = 7.6 Hz, 1H), 7.33 (dt, *J* = 7.6, 1.5 Hz, 1H), 7.30 – 7.29 (m, 1H), 7.22 (dt, *J* = 7.5, 1.5 Hz, 1H), 7.15 (d, *J* = 8.3 Hz, 1H), 6.80 (d, *J* = 2.7 Hz, 1H), 6.77 (dd, *J* = 8.4, 2.7 Hz, 1H), 4.76 – 4.72 (m, 2H), 4.14 (t, *J* = 5.7 Hz, 2H), 3.30 – 3.25 (m, 2H), 2.97 (s, 3H), 2.40 – 2.32 (m, 2H), 2.25 (s, 3H), 1.70 – 1.66 (m, 1H); **$^{13}$C NMR** (151 MHz, $CDCl_3$) δ 157.7, 141.9, 140.9, 137.1, 135.1, 131.1, 128.9, 128.5, 128.1, 125.4, 116.4, 111.8, 65.61, 65.55, 52.0, 41.1, 22.8, 20.9. Spectra in agreement with literature.[26]

**3'-(Chloromethyl)-2-methyl-4-(3-(methylsulfonyl)propoxy)-1,1'-biphenyl (38)**

To a round bottomed flask charged with compound **37** (102 mg, 0.31 mmol), dry DCM was added (0.70 mL) followed by tosyl chloride (77 mg, 0.40 mmol) and DMAP (3.7 mg, 0.03 mmol). The suspension was then flushed with Ar, cooled to 0 °C in an ice-water bath and dry $Et_3N$ (106 µL, 0.76 mmol) was added dropwise. The resulting yellow solution was stirred at rt for 17 h under Ar atmosphere. The reaction mixture was quenched with sat. aq. $NaHCO_3$ and extracted with DCM (×3). The organic layers were combined, washed with brine, dried over $MgSO_4$ and concentrated *in vacuo.* The crude material was purified by automated flash chromatography (EtOAc:*n*-heptane, 1:1) to give 26 mg (24%) of the product as an off-white oily solid: $R_f$ = 0.28 (EtOAc:*n*-heptane, 1:1); **$^1$H NMR** (400 MHz, $CDCl_3$) δ 7.44 – 7.29 (m, 3H), 7.24 (t, *J* = 1.7 Hz, 1H), 7.15 (d, *J* = 8.3 Hz, 1H), 6.84 – 6.73 (m, 2H), 4.63 (s, 2H), 4.15 (t, *J* = 5.8 Hz, 2H), 3.32 – 3.24 (m, 2H), 2.97 (s, 3H), 2.42 – 2.29 (m, 2H), 2.25 (s, 3H); **$^{13}$C NMR** (151 MHz, $CDCl_3$) δ 157.8, 142.1, 137.4, 137.1, 134.7, 131.0, 129.7, 129.6, 128.6, 126.9, 116.5, 111.8, 65.6, 52.0, 46.4, 41.0, 22.8, 20.8.

**4-((2'-Methyl-4'-(3-(methylsulfonyl)propoxy)-[1,1'-biphenyl]-3-yl)methoxy)-benzaldehyde (39)**

Compound synthesized similarly to Rexen Ulven *et al.*[24] 4-Hydroxybenzaldehyde (6.7 mg, 0.05 mmol) was dissolved in dry MeCN (1 mL) in a flame-dried vial under Ar atmosphere. Compound **38** (25 mg, 0.07 mmol) and $K_2CO_3$ (16 mg, 107 mmol) were added, and the reaction was stirred at 55 °C for 16 h. After completion, the reaction was diluted with water and extracted with EtOAc (×3). The organic phases were combined, dried over $MgSO_4$, and concentrated *in vacuo*. The residue was purified by flash column chromatography (DCM:EtOAc, 100:0 → 1:1) to give 19 mg (80%) of an off-white sticky oil: $R_f$ = 0.20 (EtOAc:*n*-heptane, 1:1); $^1$**H NMR** (400 MHz, $CDCl_3$) δ 9.89 (s, 1H), 7.88 – 7.80 (m, 2H), 7.48 – 7.34 (m, 3H), 7.31 – 7.21 (m, 1H), 7.15 (d, *J* = 8.3 Hz, 1H), 7.13 – 7.05 (m, 2H), 6.82 – 6.75 (m, 2H), 5.19 (s, 2H), 4.15 (t, *J* = 5.8 Hz, 2H), 3.31 – 3.23 (m, 2H), 2.97 (s, 3H), 2.42 – 2.31 (m, 2H), 2.23 (s, 3H); $^{13}$**C NMR** (151 MHz, $CDCl_3$) δ 190.9, 163.8, 157.8, 142.1, 137.1, 135.9, 134.8, 132.1, 131.0, 130.3, 129.5, 128.63, 128.57, 125.8, 116.5, 115.3, 111.8, 70.4, 65.6, 52.0, 41.0, 22.8, 20.8.

***trans*-3-(4-((2'-Methyl-4'-(3-(methylsulfonyl)propoxy)-[1,1'-biphenyl]-3-yl)methoxy)-phenyl)-1-oxo-2-propyl-1,2,3,4-tetrahydroisoquinoline-4-carboxylic acid (TUG-2745 - 40)**

Compound synthesized similarly to Humphries *et al.*[22] Compound **39** (9.0 mg, 21 μmol) was added to a vial and dissolved in MeOH (120 μL), followed by addition of propylamine (2 μL, 21 μmol). The vial was closed with a lid and the resulting solution was stirred at rt for 1 h. After this time, homophthalic anhydride (3.0 mg, 21 μmol) dissolved in anh. DMF (120 μL) was added slowly to the reaction mixture. The reaction was stirred at rt for 1 h before addition of 1 M $NaOH_{aq}$ (120 μL). The vial was closed with a lid again and stirred at rt for 18 h. The reaction was quenched by addition of $H_2O$, acidified to pH 1 by addition of 1 M $HCl_{aq}$ and extracted with EtOAc (×3). The organic layers were combined, washed with brine, dried over $MgSO_4$, filtered and concentrated *in vacuo*. The crude material was purified by flash column chromatography followed by preparative HPLC (20-100% B over 15 min) to give 6 mg (47%) of the product as a white solid: $t_R$ = 9.68 min (purity 99.99% by HPLC); $^1$**H NMR** (600 MHz, $CDCl_3$) δ 8.19 – 8.13 (m, 1H), 7.44 – 7.36 (m, 3H), 7.34 – 7.30 (m, 1H), 7.28 – 7.27 (m, 1H), 7.24 – 7.21 (m, 1H), 7.12 (d, *J* = 8.3 Hz, 2H), 6.96 (d, *J* = 8.7 Hz, 2H), 6.82 (d, *J* = 8.8 Hz, 2H), 6.78 (d, *J* = 2.6 Hz, 1H), 6.75 (dd, *J* = 8.4, 2.7 Hz, 1H), 5.23 (s, 1H), 5.00 (s, 2H), 4.14 (t, *J* = 5.8 Hz, 2H), 4.01 (ddd, *J* = 13.4, 9.0, 7.0 Hz, 1H), 3.90 (s,

1H), 3.90 (d, *J* = 1.5 Hz, 1H), 3.30 – 3.24 (m, 2H), 2.96 (s, 3H), 2.82 (ddt, *J* = 14.0, 9.1, 4.2 Hz, 1H), 2.39 – 2.32 (m, 2H), 2.18 (s, 3H), 1.67 – 1.57 (m, 2H), 0.87 (t, *J* = 7.4 Hz, 3H); **$^{13}$C NMR** (151 MHz, $CDCl_3$) δ 174.1, 163.9, 158.6, 157.7, 141.9, 137.1, 136.7, 134.9, 132.2, 131.7, 131.0, 129.5, 129.4, 129.2, 128.8, 128.6, 128.5, 128.1, 127.6, 125.8, 116.5, 115.2, 111.8, 70.1, 65.6, 60.7, 52.0, 51.3, 48.6, 41.1, 22.8, 21.0, 20.8, 11.5; **HRMS** calcd for $C_{37}H_{40}NO_7S^+$ (M+H$^+$) 642.2520, found 642.2520.

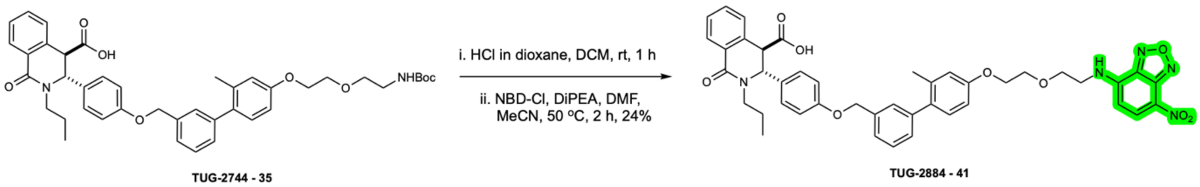


***trans*-3-(4-((2'-Methyl-4'-(2-(2-((7-nitrobenzo[*c*][1,2,5]oxadiazol-4-yl)amino)ethoxy)ethoxy)-[1,1'-biphenyl]-3-yl)methoxy)phenyl)-1-oxo-2-propyl-1,2,3,4-tetrahydroisoquinoline-4-carboxylic acid (TUG-2884)**

To a solution of *trans*-3-(4-((4'-(2-(2-((*tert*-butoxycarbonyl)amino)ethoxy)ethoxy)-2'-methyl-[1,1'-biphenyl]-3-yl)methoxy)phenyl)-1-oxo-2-propyl-1,2,3,4-tetrahydroisoquinoline-4-carboxylic acid (31.3 mg, 44.2 μmol) in DCM (0.1 mL) was added 4 M HCl in dioxane (99 μL, 0.40 mmol) and stirred at room temperature for 1 h. The reaction mixture was concentrated, and the resulting crude amine was dissolved in DMF:MeCN (1:3; 400 μL) and NBD-Cl (9.7 mg, 48.6 μmol) was added followed by DIPEA (84.6 μL, 0.49 mmol). The vial was capped and stirred in dark at 50 °C for 2 h. The crude reaction mixture was purified by preparative HPLC (65% B for 12 min) to afford 8.3 mg (24%) of the product as an orange solid : $\boldsymbol{t_R}$ = 10.87 min (purity >99.99% by HPLC; 40-100% B in 15 min); $\boldsymbol{R_f}$ = 0.21 (MeOH:DCM, 1:9); **$^1$H NMR** (600 MHz, $CDCl_3$) δ 8.47 (d, *J* = 8.6 Hz, 1H), 8.20 – 8.14 (m, 1H), 7.45 – 7.36 (m, 3H), 7.34 – 7.30 (m, 1H), 7.28 – 7.27 (m, 1H), 7.24 – 7.21 (m, 1H), 7.16 – 7.08 (m, 2H), 7.00 – 6.94 (m, 2H), 6.87 – 6.81 (m, 2H), 6.79 (d, *J* = 2.6 Hz, 1H), 6.75 (dd, *J* = 8.4, 2.7 Hz, 1H), 6.68 – 6.61 (m, 1H), 6.21 (d, *J* = 8.6 Hz, 1H), 5.24 (s, 1H), 5.01 (s, 2H), 4.21 – 4.15 (m, 2H), 4.06 – 3.99 (m, 1H), 3.98 – 3.90 (m, 5H), 2.86 – 2.79 (m, 1H), 2.17 (s, 3H), 1.69 – 1.58 (m, 2H), 0.89 (t, *J* = 7.4 Hz, 3H); **HRMS** calcd for $C_{43}H_{42}N_5O_9^+$ (M+H $^+$) 772.2976, found 772.2970.

### Experimental details for spectroscopic measurements

**Absorbance measurements:** the solutions of TUG-2597 and TUG-2884 were prepared by diluting the 10 mM DMSO stock solution of the fluorophore in *n*-octanol (HPLC grade, Sigma-Aldrich) and in 0.01 M $PBS_{7.4}$ (P5368, Sigma-Aldrich). The fluorescence standard fluorescein was dissolved in 0.1 M $NaOH_{aq}$, to make a 10 mM stock solution. The solutions of TUG-2591 and Rhodamine 800 were prepared by diluting 10 mM DMSO stock solutions of the corresponding compounds in DMSO (for spectroscopy Uvasol®, Sigma-Aldrich) and abs. EtOH (for spectroscopy Uvasol®, Sigma-Aldrich), respectively.

For determination of extinction coefficients, absorption spectra at increasing concentrations of TUG-2597, TUG-2884, TUG-2591, fluorescein and Rhodamine 800 were recorded in duplicate using a flat-bottom transparent 96-well plate (Greiner) on a microplate reader Safire 2 (Tecan) with XFLUOR4SAFIREII Version: V 4.62n software.

**Emission measurements:** the solutions of the test compounds and references were prepared using different solvents as described above. For quantum yield determination, emission spectra at increasing concentrations of fluorescein (internal standard, $\Phi$ = 0.93[22]), TUG-2884 and TUG-2597 were recorded by exciting **all** fluorophores at a fixed wavelength of 450 nm. The measurements were performed in duplicate on a LS50B luminescence (PerkinElmer) spectrometer, using 10 mm-path cuvettes, an excitation slit of 5.0 nm and a scan speed of 300 nm $min^{-1}$.

Similarly, the emission spectra at increasing concentrations of Rhodamine 800 (internal standard, $\Phi$ = 0.25[23]) and TUG-2591 were recorded by exciting both fluorophores at a fixed wavelength of 635 nm. The measurements were performed in duplicate on a FluoTime 300 (PicoQuant) spectrometer, using 10 mm-path cuvettes and a bandwidth of 2 nm. All experiments were performed at room temperature. Data analysis was performed using Graphpad Prism v10 and emission spectra were background-subtracted using Spectragryph version v1.2.16.1.[24]

### Spectroscopic characterization of fluorescent tracers

The absorption and emission spectra of TUG-2597 and TUG-2884 were recorded in $PBS_{7.4}$ and in *n*-octanol. TUG-2597 showed a Stoke's shift of 52 nm and an extinction coefficient (ε) of 16340 $M^{-1}$ $cm^{-1}$, TUG-2884 showed a Stoke's shift of 61 nm and an extinction coefficient (ε) of 12363 $M^{-1}$ $cm^{-1}$, values in the range of previously developed

NBD-tracers[1,21,30] (Figure S12, Figure S13, Table S6), with a quantum yield (QY) of 0.22 and 0.27, respectively (Table S6).

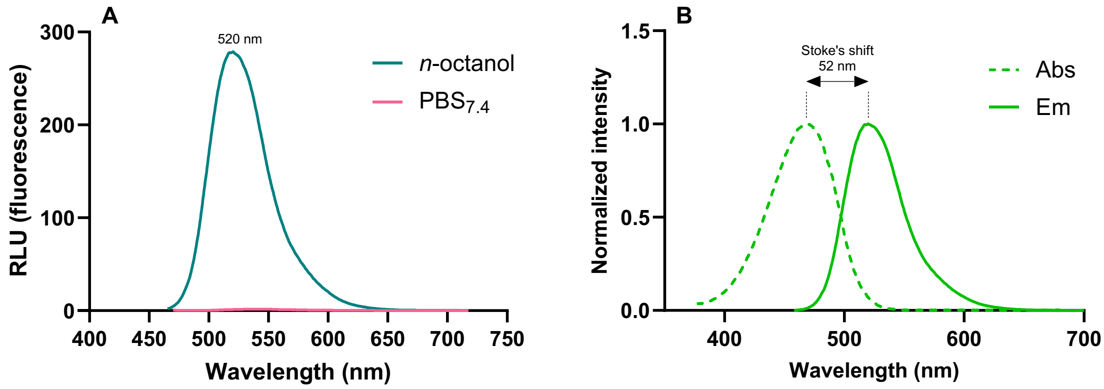


***Figure S12.*** A: Emission of TUG-2597 in $PBS_{7.4}$ *vs* *n*-octanol at 8.50 μM (excitation at 450 nm). B: Normalized absorbance and emission spectra of TUG-2597 in *n*-octanol.

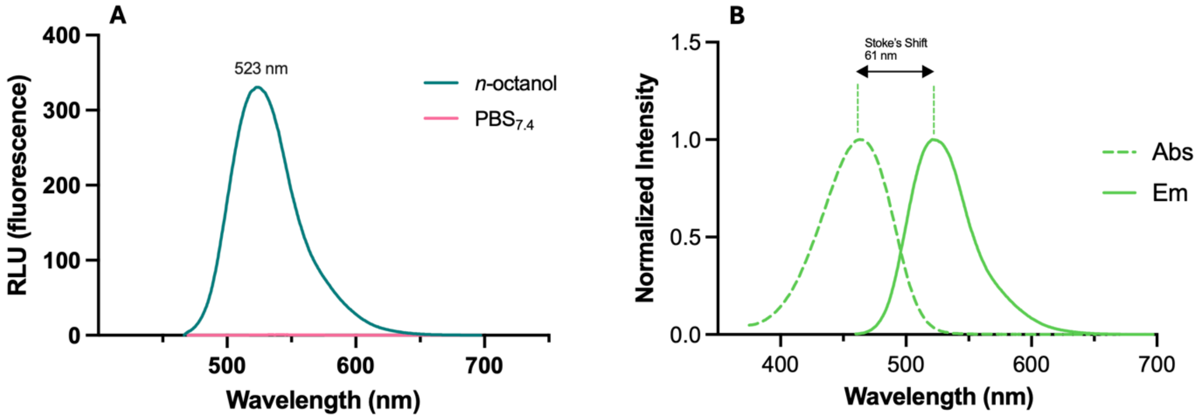


***Figure S13.*** A: Emission of TUG-2884 in $PBS_{7.4}$ *vs* *n*-octanol at 8.57 μM (excitation at 462 nm). B: Normalized absorbance and emission spectra of TUG-2884 in *n*-octanol.

The sulfo-cyanine based fluorescent tracer TUG-2591 was not fully soluble in $PBS_{7.4}$, therefore all measurements were performed in DMSO. TUG-2591 exhibits a Stoke's shift of 22 nm (Figure S14, Table S6), and a quantum yield of 0.35 (Table S6). In summary, the observed spectroscopic properties of the three tracers are summarized in Table S6.

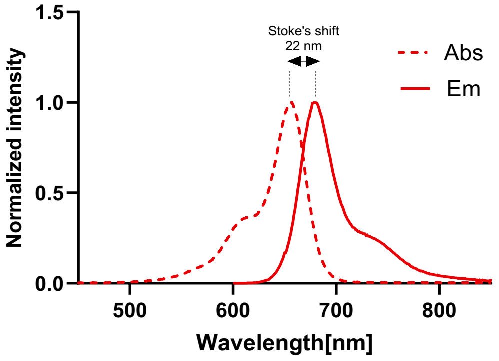


***Figure S14.*** Normalized absorbance and emission spectra of TUG-2591 in DMSO.

***Table S6.*** overview of absorption and fluorescence properties of TUG-2597 and TUG-2591.

| Compound | Solvent | $\lambda_{ex}$ [nm] | $\lambda_{em}$ [nm] | SS[a] [nm] | ε[b] [$M^{-1}$ $cm^{-1}$] | QY[c] |
|---|---|---|---|---|---|---|
| **TUG-2597** | *n*-octanol | 468 | 520 | 52 | 16340 | 0.22 |
| | $PBS_{7.4}$ | 486 | 537 | n.d.[d] | n.d.[d] | n.d.[d] |
| **TUG-2591** | DMSO | 656 | 678 | 22 | 119071 | 0.35 |
| **TUG-2884** | *n*-octanol | 462 | 523 | 61 | 12363 | 0.27 |
| | $PBS_{7.4}$ | 467 | 539 | n.d.[d] | n.d.[d] | n.d.[d] |

[a]SS, Stokes shift; [b]ε, molar extinction coefficient; [c]QY, relative fluorescence quantum yield (fluorescein in 0.1 M $NaOH_{aq}$ used as an internal standard for TUG-2597 and TUG-2884, Rhodamine 800 in EtOH used for TUG-2591); [d]n.d.: not determined.

***Supplemental References***


1 Christiansen, E. *et al.* Free fatty acid receptor 1 (FFA1/GPR40) agonists:mesylpropoxy appendage lowers lipophilicity and improves ADME properties. *Journal Medicinal Chemistry* **55**, 6624-6628 (2012).

2 Brown, S. P., Dranfield, P., Fu, Z., Houze, J., Jiao, X. Y., Kohn, T. J., Pattaropong, V., Vimolratana, M., & Schmitt, M. J. Substituted biphenyl phenoxy-, thiophenyl- and aminophenylpropanoic acid gpr40 modulators. WO2008/130514 (2008).

2 Brown, S. P. *et al.* Discovery of AM-1638: A Potent and Orally Bioavailable GPR40/FFA1 Full Agonist. *ACS Medicinal Chemistry Letters* **3**, 726-730 (2012).

3 Capitta, F., De Luca, L. & Porcheddu, A. “Quick and click” assembly of functionalised indole rings via metal-promoted cyclative tandem reactions. *RSC Advances* **4**, 59297-59301 (2014).

4 Furukawa, H. *et al.* Design and Identification of a GPR40 Full Agonist (SCO-267) Possessing a 2-Carbamoylphenyl Piperidine Moiety. *Journal of Medicinal Chemistry* **63**, 10352-10379 (2020).

5 Swain, N. A., Brown, A. D., Jones, L., H., Marron, B. E., Rawson, D. J., Ryckmans, T., Storer, R. I., & West, C. W. Benzenesulfonamides useful as sodium channel inhibitors. WO2015/18179 (2015).

6 Hauel, N., Ceci, A., Doods, H., Kauffmann-Hefner, I., Konetztki, I., Schuler-Metz, A., & Walter, R. Novel compounds. US2011/0294775 (2011).

7 Nitz, T. J., Wild, C. T., Martin, D. E., & Freed, E. O. Triterpene amine derivatives WO2020/006510 (2020).

8 Bartole, E. *et al.* [3H]UR-DEBa176: A 2,4-Diaminopyrimidine-Type Radioligand Enabling Binding Studies at the Human, Mouse, and Rat Histamine H4 Receptors. *Journal of Medicinal Chemistry* **62**, 8338-8356 (2019).

9 Aida, J., Yoshitomi, Y., Hitomi, Y., Noguchi, N., Hirata, Y., Furukawa, H., Shibuya, A., Watanabe, K., Miyamoto, Y., Okawa, T., Takakura, N., & Miwatashi, S. Aromatic compound. US2016/0115128 (2016).

10 Chadwick, J. *et al.* Design, synthesis and antimalarial/anticancer evaluation of spermidine linked artemisinin conjugates designed to exploit polyamine transporters in Plasmodium falciparum and HL-60 cancer cell lines. *Bioorganic & Medicinal Chemistry* **18**, 2586-2597 (2010).

11 Maudoux, N. *et al.* $C_{sp3}$-H Bond Activation with Triel Metals: Indium and Gallium Zwitterions through Internal Hydride Abstraction in Rigid Salan Ligands. *Chemistry – A European Journal* **20**, 7706-7717 (2014).

12 Vantourout, J. C., Law, R. P., Isidro-Llobet, A., Atkinson, S. J. & Watson, A. J. B. Chan–Evans–Lam Amination of Boronic Acid Pinacol (BPin) Esters: Overcoming the Aryl Amine Problem. *The Journal of Organic Chemistry* **81**, 3942-3950 (2016).

13 Due-Hansen, M. E. *et al.* A protocol for amide bond formation with electron deficient amines and sterically hindered substrates. *Organic & Biomolecular Chemistry* **14**, 430-433 (2016).

14 Holmes, D. L., Smith, E. M. & Nowick, J. S. Solid-Phase Synthesis of Artificial β-Sheets. *Journal of the American Chemical Society* **119**, 7665-7669 (1997).

15 Taliani, S. *et al.* New Fluorescent 2-Phenylindolglyoxylamide Derivatives as Probes Targeting the Peripheral-Type Benzodiazepine Receptor: Design, Synthesis, and Biological Evaluation. *Journal of Medicinal Chemistry* **50**, 404-407 (2007).

16 Robertson, N. S. *et al.* Macrocyclisation and functionalisation of unprotected peptides via divinyltriazine cysteine stapling. *Chemical Communications* **55**, 9499-9502 (2019).

17 Davis, G. J., Sofka, H. A. & Jewett, J. C. Highly Stable Meldrum's Acid Derivatives for Irreversible Aqueous Covalent Modification of Amines. *Organic Letters* **22**, 2626-2629 (2020).

18 Christiansen, E., Hudson, B. D., Hansen, A. H., Milligan, G. & Ulven, T. Development and Characterization of a Potent Free Fatty Acid Receptor 1 (FFA1) Fluorescent Tracer. *Journal of Medicinal Chemistry* **59**, 4849-4858 (2016).

19 Bunschoten, A. *et al.* Tailoring Fluorescent Dyes To Optimize a Hybrid RGD-Tracer. *Bioconjugate Chemistry* **27**, 1253-1258 (2016).

20 Shulov, I. *et al.* Protein-Sized Bright Fluorogenic Nanoparticles Based on Cross-Linked Calixarene Micelles with Cyanine Corona. *Angewandte Chemie International Edition* **55**, 15884-15888 (2016).

21 Ciba, M., Dibnah, B., Hudson, B. D. & Rexen Ulven, E. Development and Characterization of Potent Succinate Receptor Fluorescent Tracers. *Journal of Medicinal Chemistry* **66**, 8951-8974 (2023).

22 Humphries, P. S. *et al.* Chemically-enabled synthesis of 1,2,3,4-tetrahydroisoquinolin-1-ones. *Tetrahedron Letters* **50**, 2140-2143 (2009).

23 Ju, E. J. *et al.* Screening, Synthesis, and In Vitro Evaluation of Vinyl Sulfones as Inhibitors of Complement-Dependent Cytotoxicity in Neuromyelitis Optica. *ChemMedChem* **11**, 377-381 (2016).

24 Rexen Ulven, E. *et al.* Structure-Activity Investigations and Optimisations of Non-metabolite Agonists for the Succinate Receptor 1. *Scintific Reports* **8**, 10010 (2018).

26 Kumar, S., Sharma, R., Mahajan, V. A. & Sawargave, S. P. Phenyl alkanoic acid derivatives as gpr agonists. WO2013/128378 (2013).

27 Brouwer, A. M. Standards for photoluminescence quantum yield measurements in solution (IUPAC Technical Report). **83**, 2213-2228 (2011).

28 Alessi, A., Salvalaggio, M. & Ruzzon, G. Rhodamine 800 as reference substance for fluorescence quantum yield measurements in deep red emission range. Journal of Luminescence **134**, 385-389 (2013).

29 Menges, F. "Spectragryph - optical spectroscopy software" Version 1.2.16.1, 2023, http://www.effemm2.de/spectragryph/.

30 Hansen, A. H. et al. Development and Characterization of a Fluorescent Tracer for the Free Fatty Acid Receptor 2 (FFA2/GPR43). Journal of Medicinal Chemistry **60**, 5638-5645 (2017).

### ***NMR Spectra and Analytical HPLC Traces***

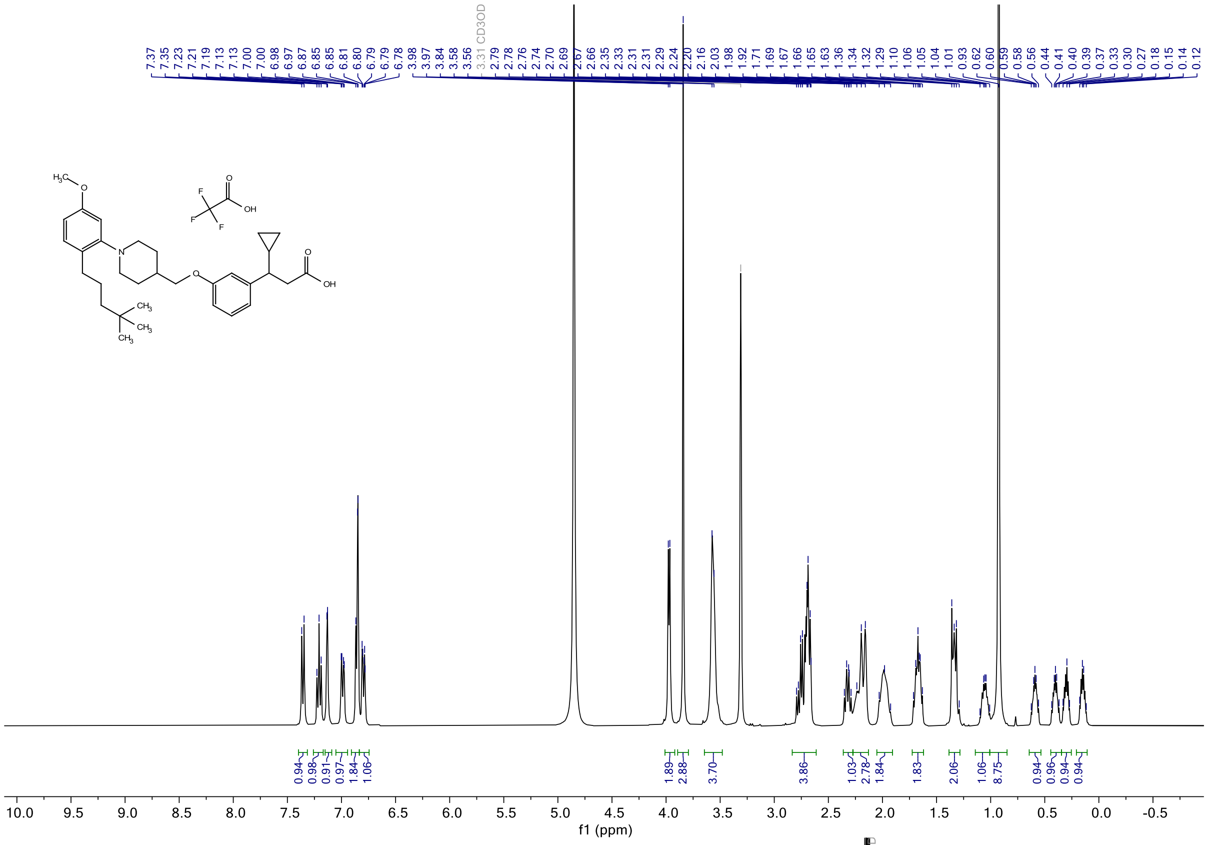

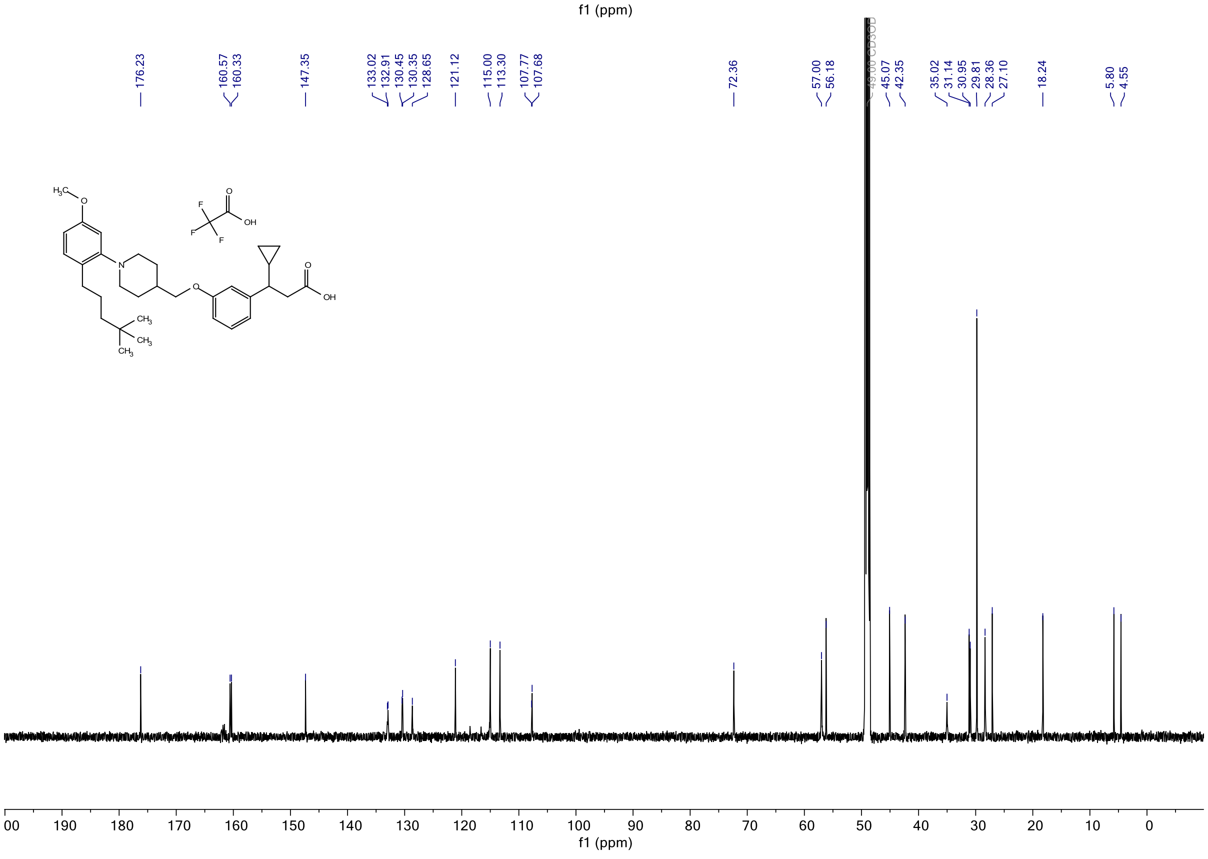

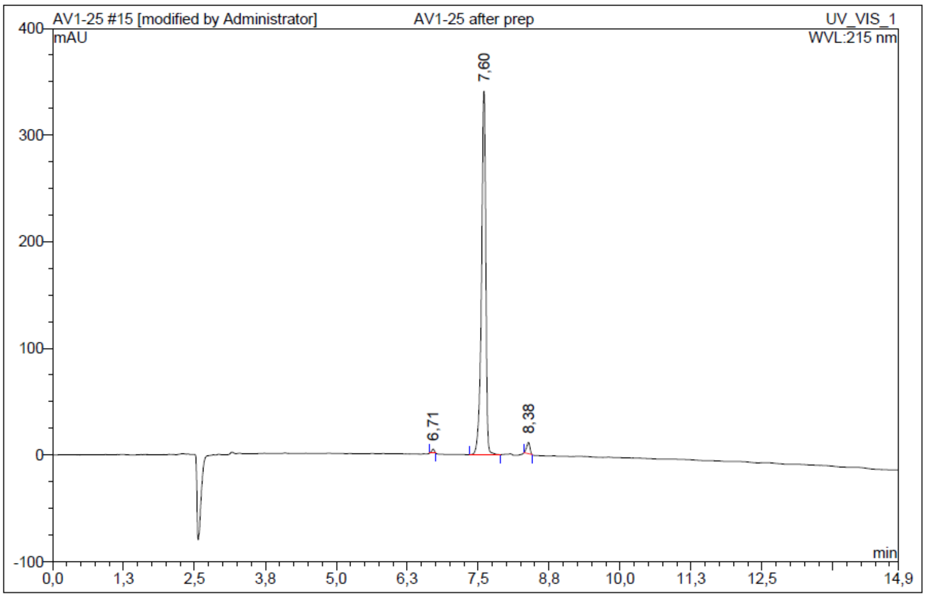


| No. | Ret.Time min | Peak Name | Height mAU | Area mAU*min | Rel.Area % | Amount | Resolution(EP) |
|---|---|---|---|---|---|---|---|
| 1 | 6,71 | n.a. | 3,588 | 0,190 | 0,63 | n.a. | 8,33 |
| 2 | 7,60 | n.a. | 340,937 | 29,384 | 97,03 | n.a. | 6,61 |
| 3 | 8,38 | n.a. | 10,610 | 0,710 | 2,35 | n.a. | n.a. |
| Total: | | | 355,135 | 30,284 | 100,00 | 0,000 | |

***Figure S15.*** $^1$H NMR, $^{13}$C NMR and HPLC chromatogram of **T360 - 8**.

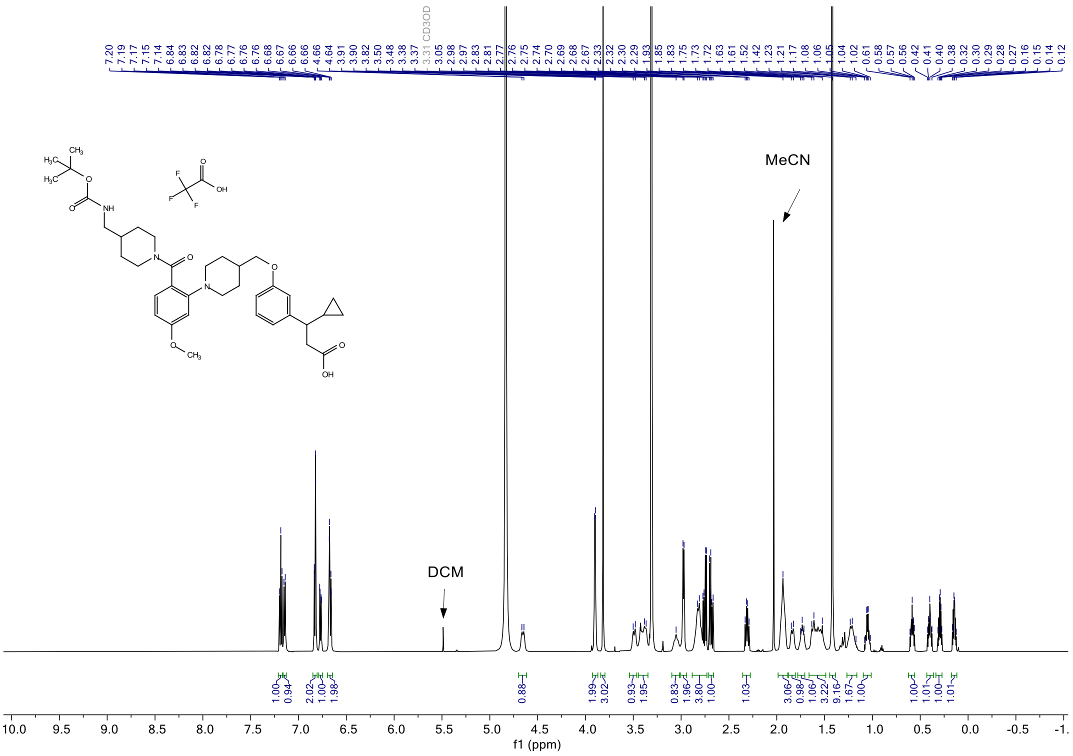

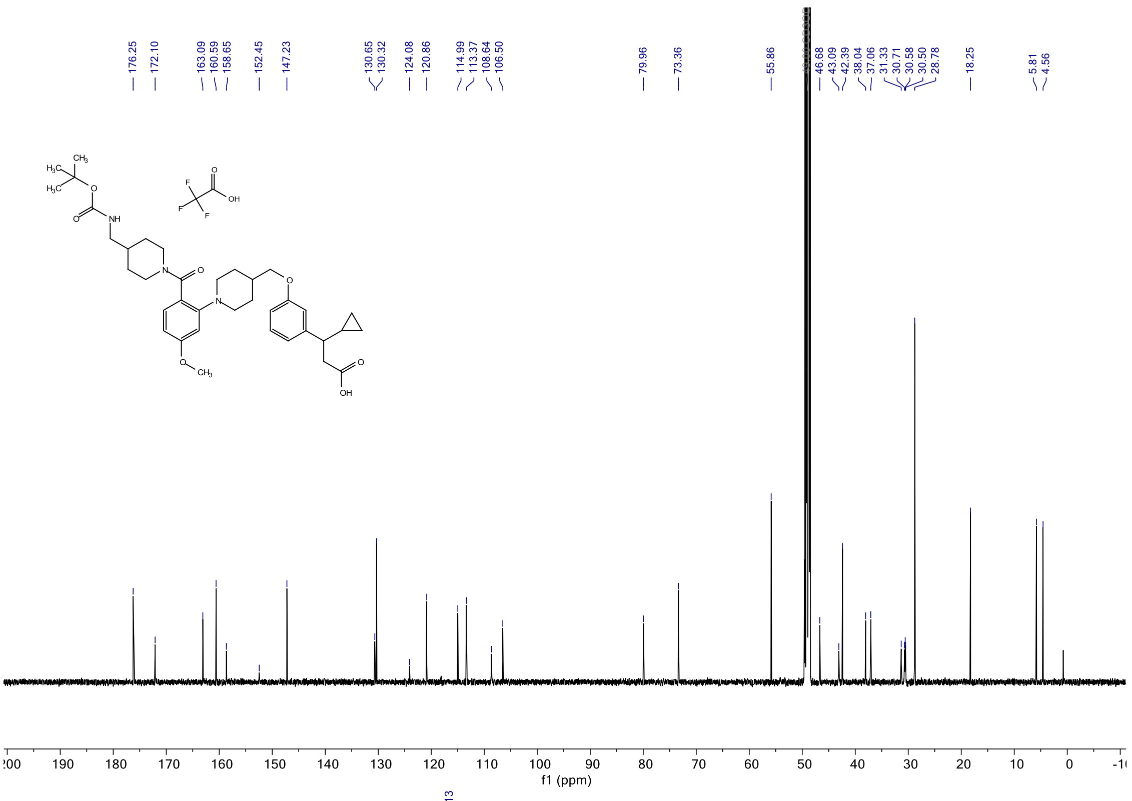
176.25
172.10
163.09
160.59
158.65
152.45
147.23
130.65
130.32
124.08
120.86
114.99
113.37
108.64
106.50
79.96
73.36
55.86
46.68
43.09
42.39
38.04
37.06
31.33
30.71
30.58
30.50
28.78
18.25
5.81
4.56
f1 (ppm)

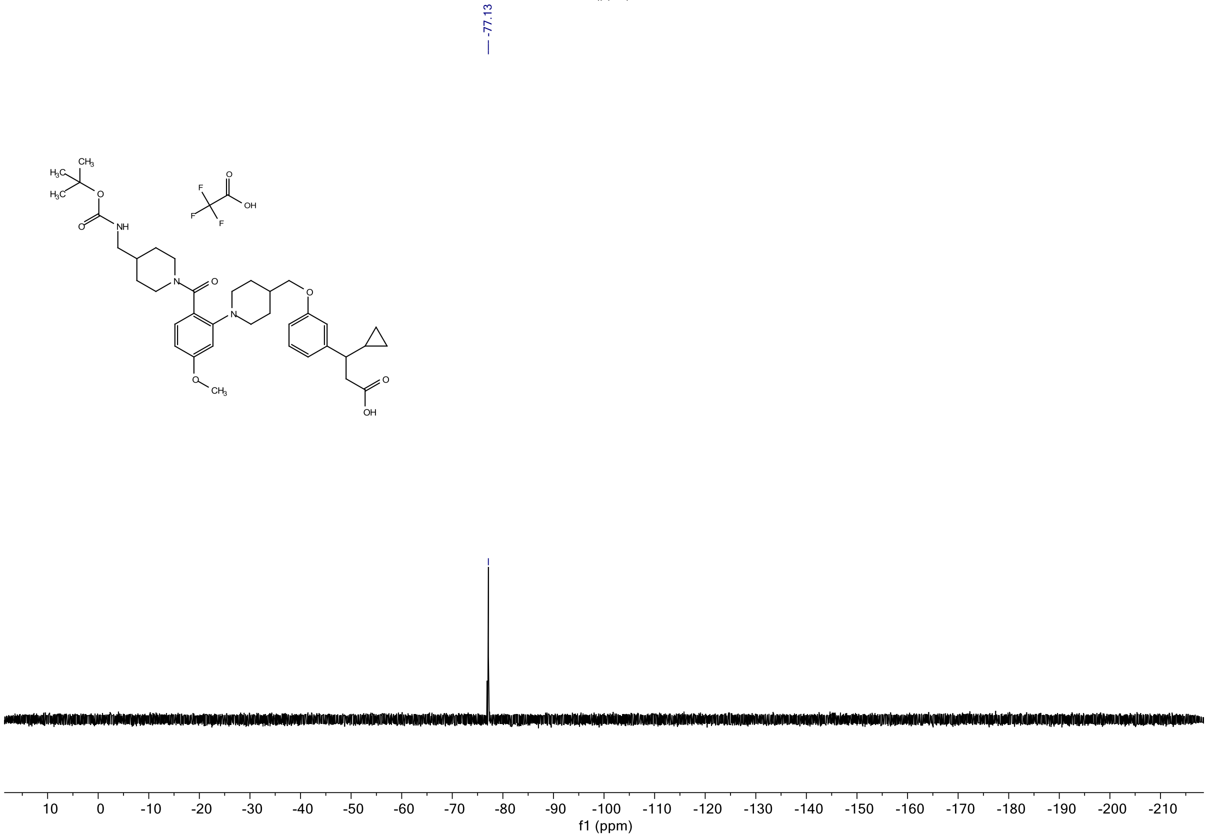
-77.13
f1 (ppm)

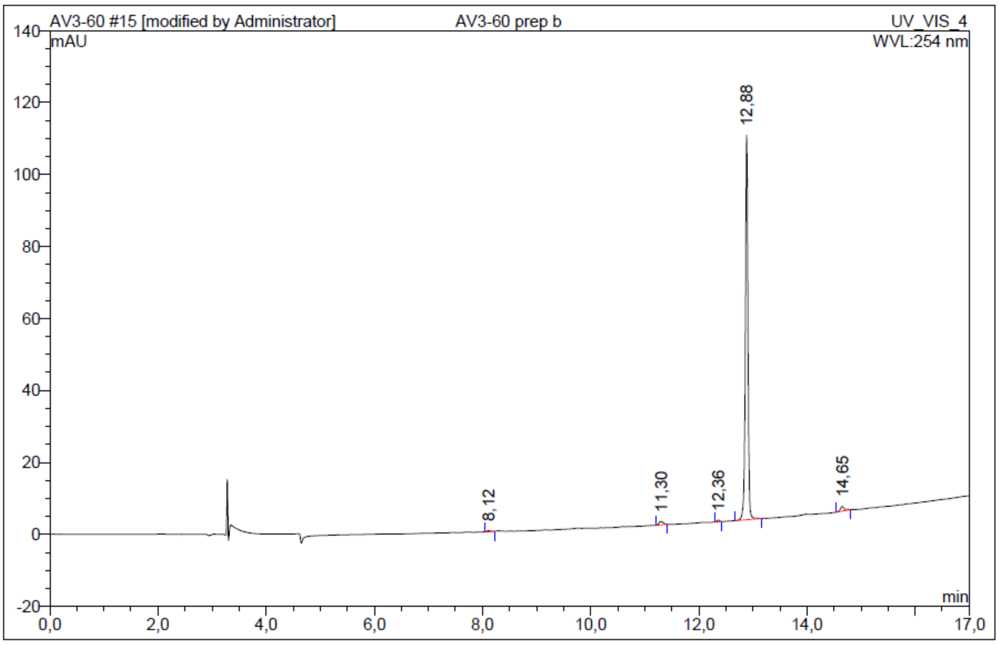


| No. | Ret.Time min | Peak Name | Height mAU | Area mAU*min | Rel.Area % | Amount | Resolution(EP) |
|---|---|---|---|---|---|---|---|
| 1 | 8,12 | n.a. | 0,309 | 0,027 | 0,43 | n.a. | 22,01 |
| 2 | 11,30 | n.a. | 0,950 | 0,088 | 1,38 | n.a. | 9,10 |
| 3 | 12,36 | n.a. | 0,411 | 0,022 | 0,35 | n.a. | 6,11 |
| 4 | 12,88 | n.a. | 106,913 | 6,092 | 96,10 | n.a. | 17,70 |
| 5 | 14,65 | n.a. | 1,265 | 0,110 | 1,74 | n.a. | n.a. |
| Total: | | | 109,848 | 6,339 | 100,00 | 0,000 | |

***Figure S16***. $^{1}$H NMR, $^{13}$C NMR, $^{19}$F and HPLC chromatogram of **TUG-2450 - 14a**.

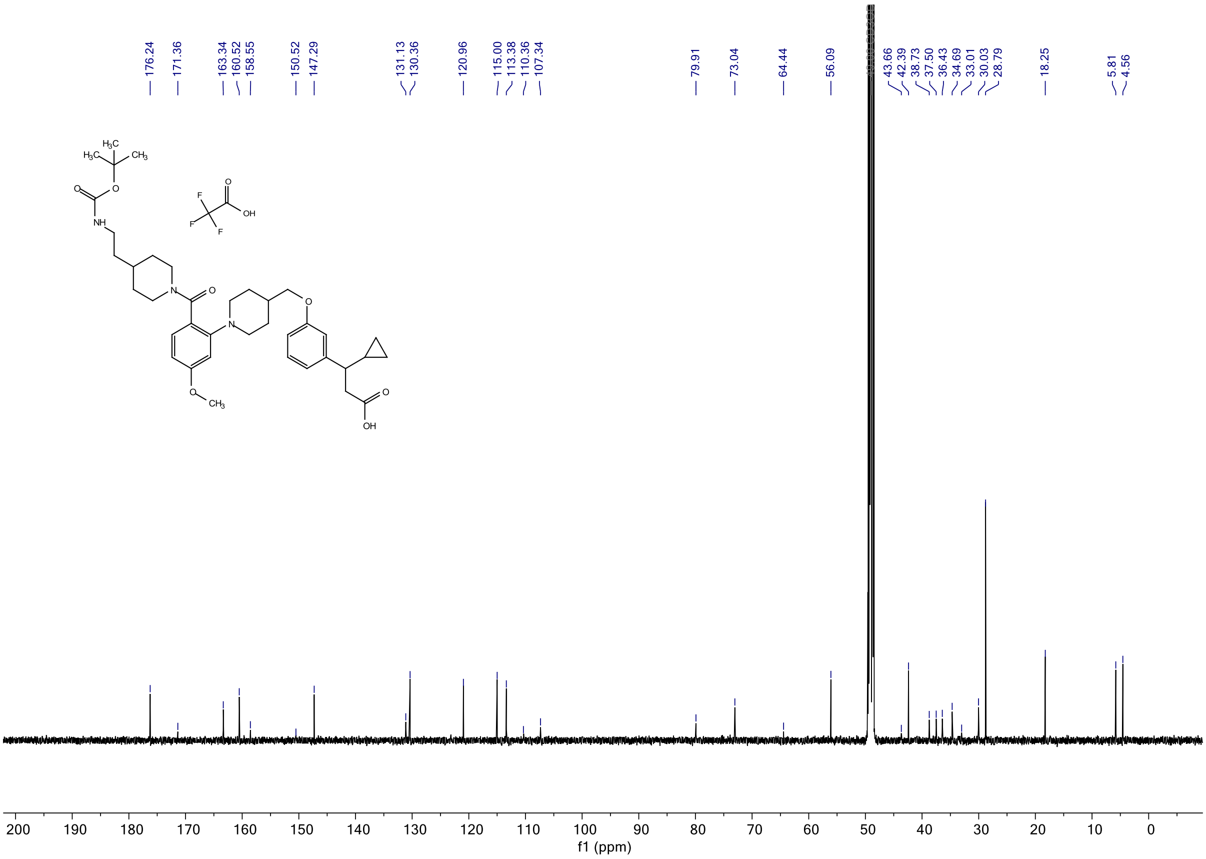

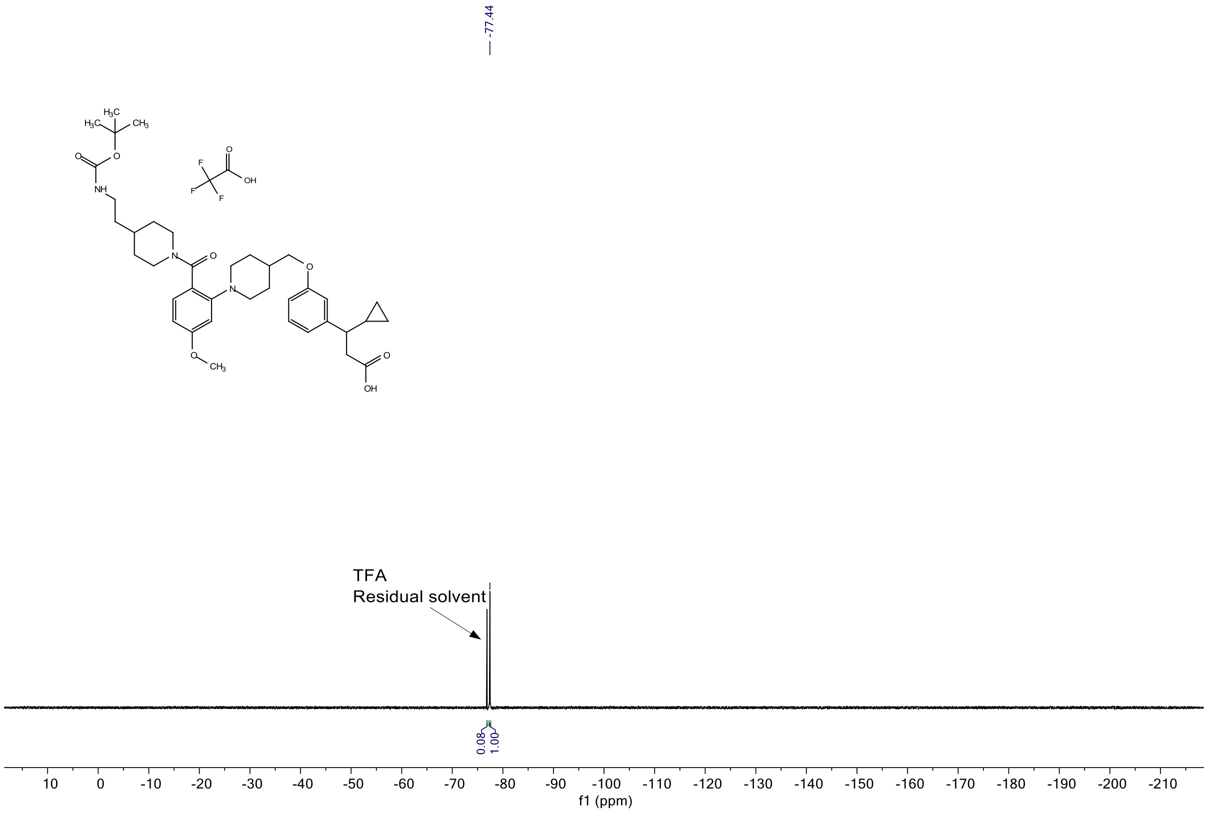


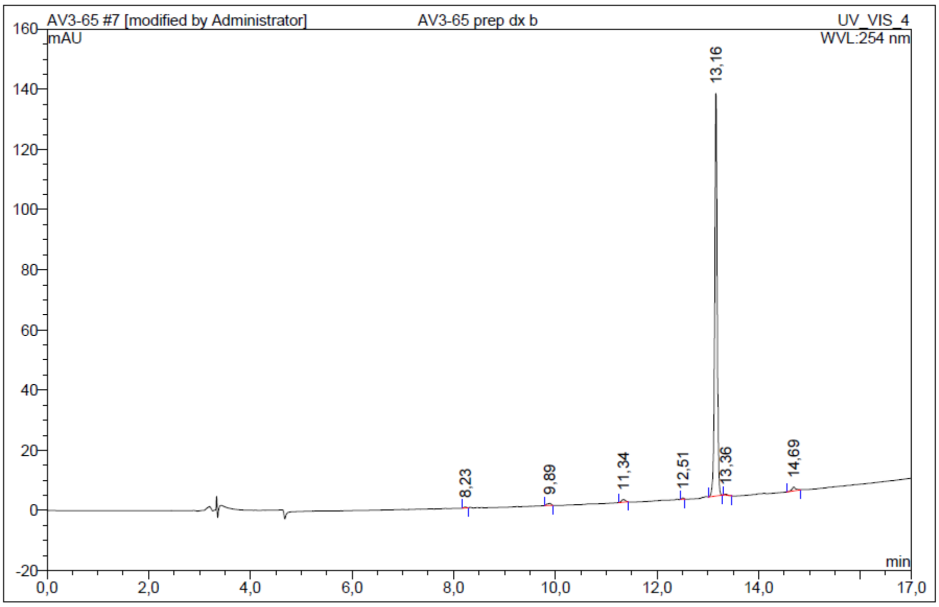


| No. | Ret.Time min | Peak Name | Height mAU | Area mAU*min | Rel.Area % | Amount | Resolution(EP) |
|---|---|---|---|---|---|---|---|
| 1 | 8,23 | n.a. | 0,339 | 0,019 | 0,24 | n.a. | 13,52 |
| 2 | 9,89 | n.a. | 0,669 | 0,056 | 0,71 | n.a. | 10,09 |
| 3 | 11,34 | n.a. | 0,925 | 0,077 | 0,97 | n.a. | 11,03 |
| 4 | 12,51 | n.a. | 0,384 | 0,018 | 0,22 | n.a. | 7,96 |
| 5 | 13,16 | n.a. | 133,830 | 7,595 | 96,13 | n.a. | 1,97 |
| 6 | 13,36 | n.a. | 0,392 | 0,028 | 0,35 | n.a. | 11,80 |
| 7 | 14,69 | n.a. | 1,277 | 0,109 | 1,38 | n.a. | n.a. |
| Total: | | | 137,815 | 7,902 | 100,00 | 0,000 | |

***Figure S17.*** $^{1}H$ NMR, $^{13}C$ NMR, $^{19}F$ NMR and HPLC chromatogram of **TUG-2451 - 14b**.

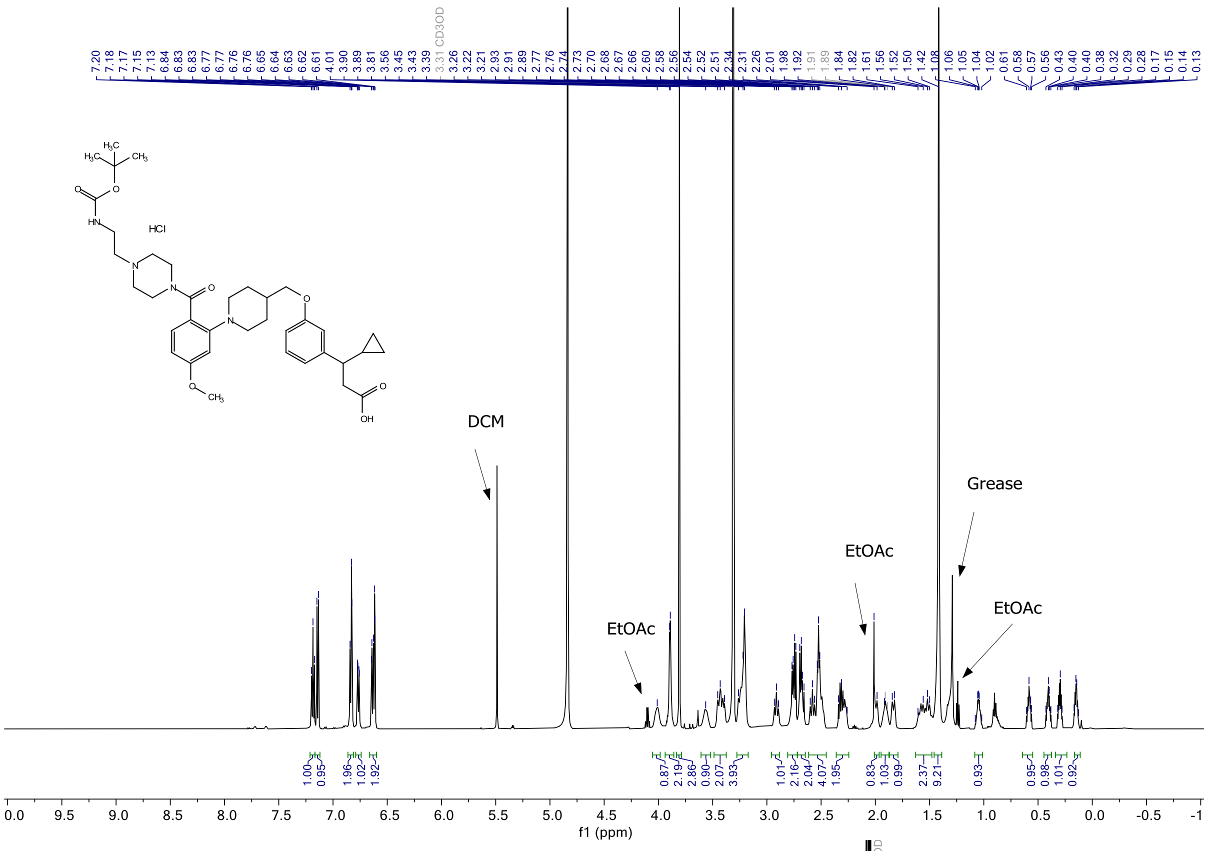

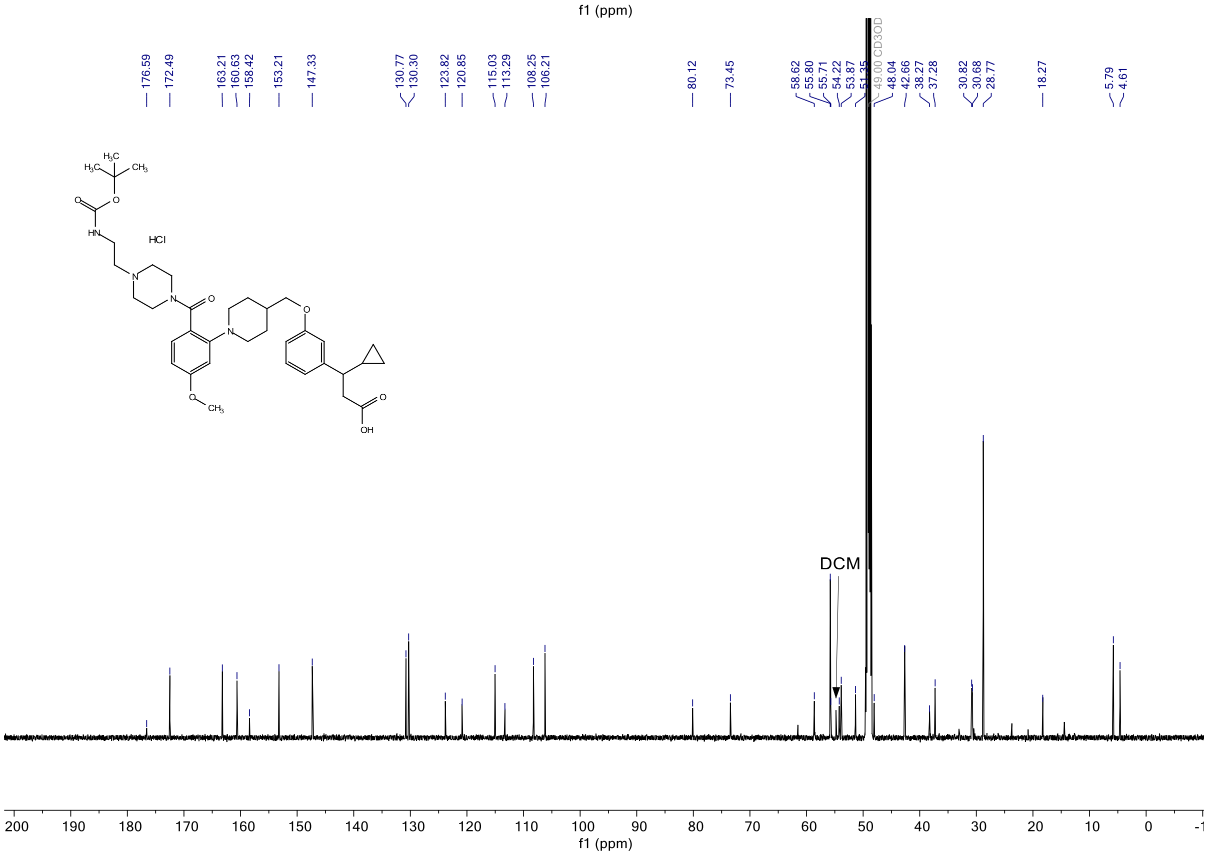

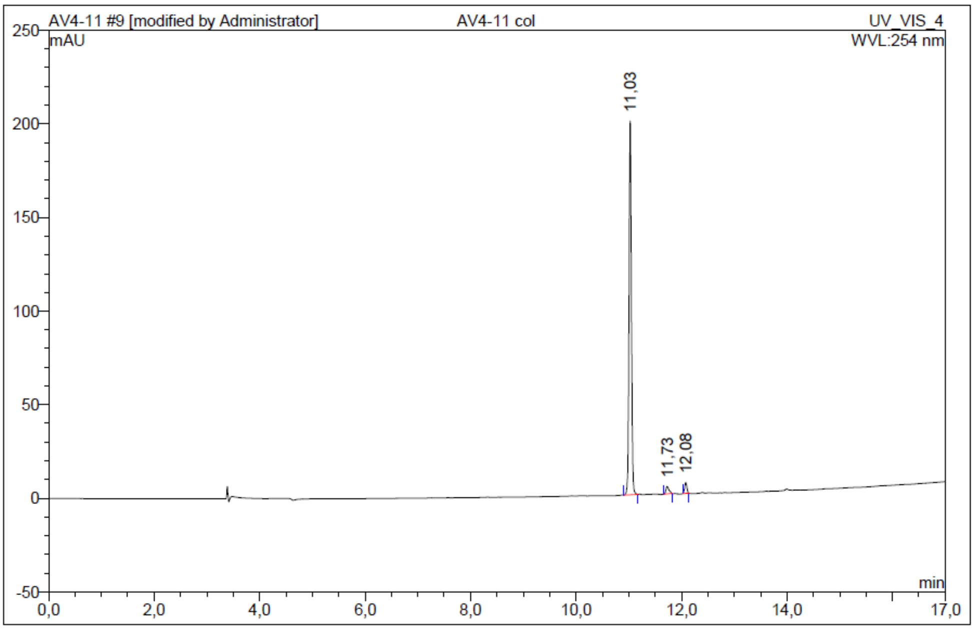


| No. | Ret.Time min | Peak Name | Height mAU | Area mAU*min | Rel.Area % | Amount | Resolution(EP) |
|---|---|---|---|---|---|---|---|
| 1 | 11,03 | n.a. | 199,525 | 10,777 | 95,11 | n.a. | 7,77 |
| 2 | 11,73 | n.a. | 3,950 | 0,258 | 2,28 | n.a. | 3,96 |
| 3 | 12,08 | n.a. | 5,827 | 0,296 | 2,61 | n.a. | n.a. |
| Total: | | | 209,302 | 11,331 | 100,00 | 0,000 | |

***Figure S18.*** $^{1}$H NMR, $^{13}$C NMR and HPLC chromatogram of **TUG-2458 - 14c**.

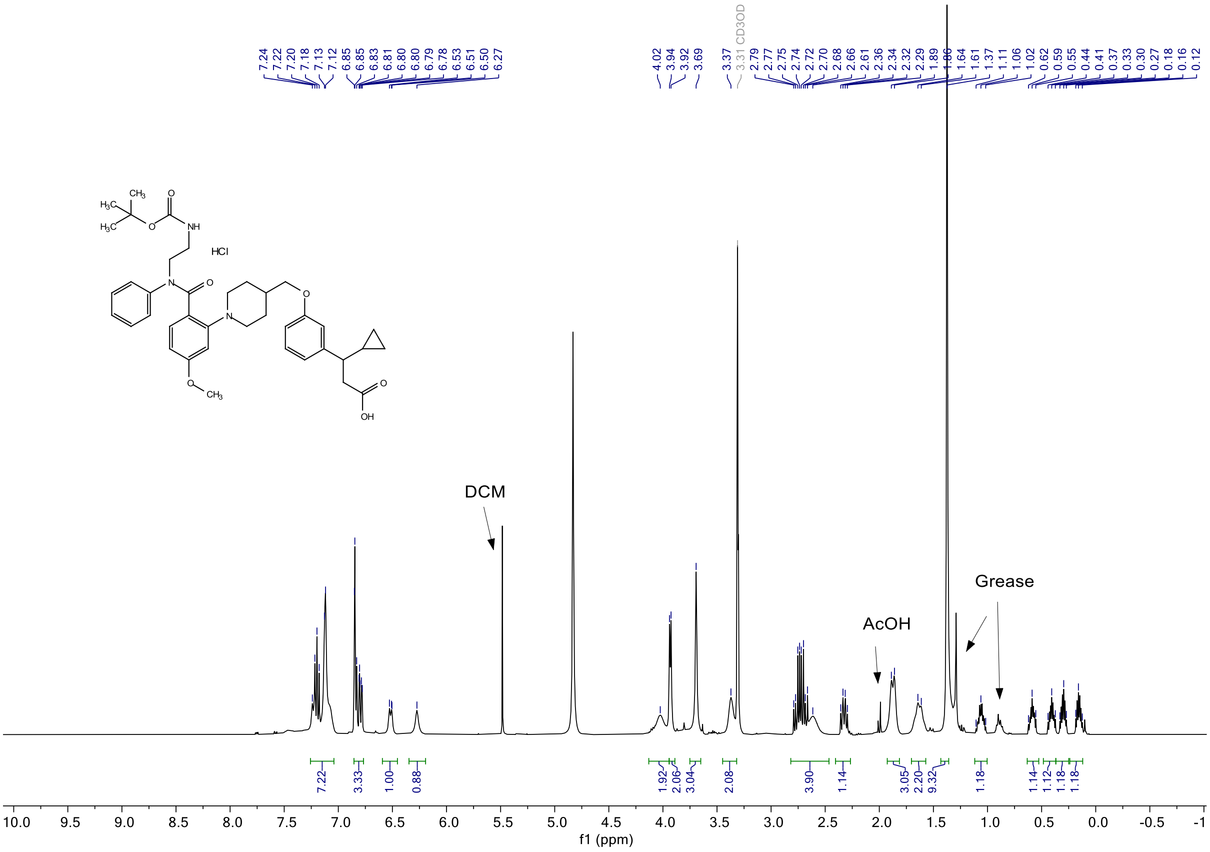

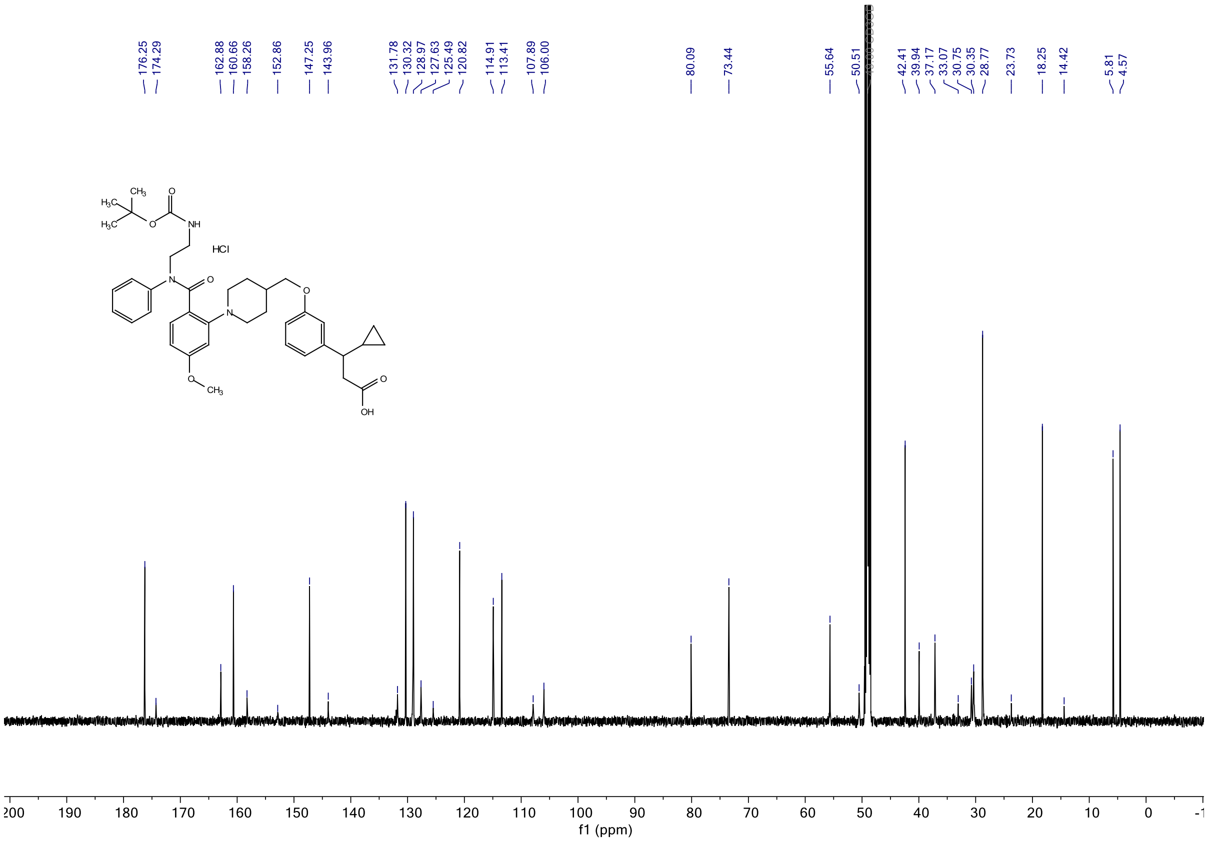


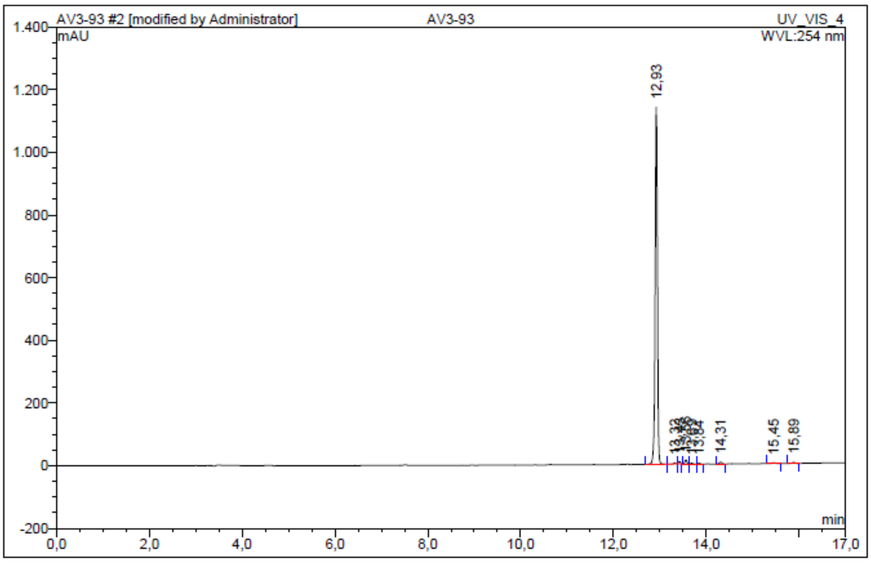


| No. | Ret.Time min | Peak Name | Height mAU | Area mAU*min | Rel.Area % | Amount | Resolution(EP) |
|---|---|---|---|---|---|---|---|
| 1 | 12,93 | n.a. | 1140,295 | 61,555 | 96,30 | n.a. | 5,03 |
| 2 | 13,32 | n.a. | 3,236 | 0,159 | 0,25 | n.a. | 1,33 |
| 3 | 13,43 | n.a. | 7,865 | 0,382 | 0,60 | n.a. | 1,70 |
| 4 | 13,56 | n.a. | 12,789 | 0,653 | 1,02 | n.a. | 1,55 |
| 5 | 13,69 | n.a. | 3,155 | 0,166 | 0,26 | n.a. | 1,80 |
| 6 | 13,84 | n.a. | 2,963 | 0,162 | 0,25 | n.a. | 5,23 |
| 7 | 14,31 | n.a. | 6,325 | 0,409 | 0,64 | n.a. | 9,27 |
| 8 | 15,45 | n.a. | 2,464 | 0,228 | 0,36 | n.a. | 3,58 |
| 9 | 15,89 | n.a. | 3,134 | 0,206 | 0,32 | n.a. | n.a. |
| Total: | | | 1182,227 | 63,919 | 100,00 | 0,000 | |

***Figure S19.*** $^{1}H$ NMR, $^{13}C$ NMR and HPLC chromatogram of **TUG-2455 - 14d**.

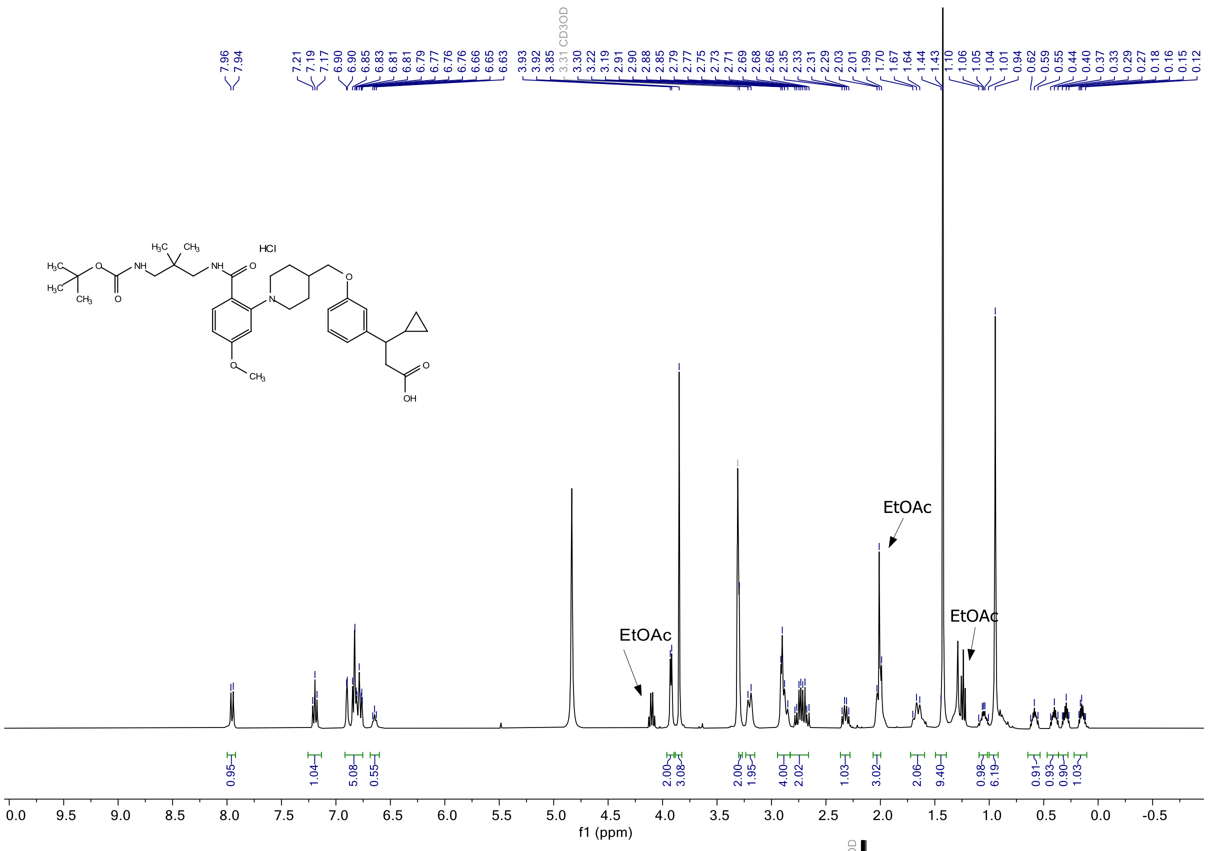
7.96
7.94
7.21
7.19
7.17
6.90
6.90
6.85
6.83
6.81
6.81
6.79
6.77
6.76
6.76
6.66
6.65
6.63
3.93
3.92
3.85
3.31 CD3OD
3.30
3.22
3.19
2.91
2.90
2.88
2.85
2.79
2.77
2.75
2.73
2.71
2.69
2.68
2.66
2.35
2.33
2.31
2.29
2.03
2.01
1.99
1.70
1.67
1.64
1.44
1.43
1.10
1.06
1.05
1.04
1.01
0.94
0.62
0.59
0.55
0.44
0.40
0.37
0.33
0.29
0.27
0.18
0.16
0.15
0.12
HCl
EtOAc
EtOAc
EtOAc
0.95
1.04
5.08
0.55
2.00
3.08
2.00
1.95
4.00
2.02
1.03
3.02
2.06
9.40
0.98
6.19
0.91
0.93
0.90
1.03
f1 (ppm)

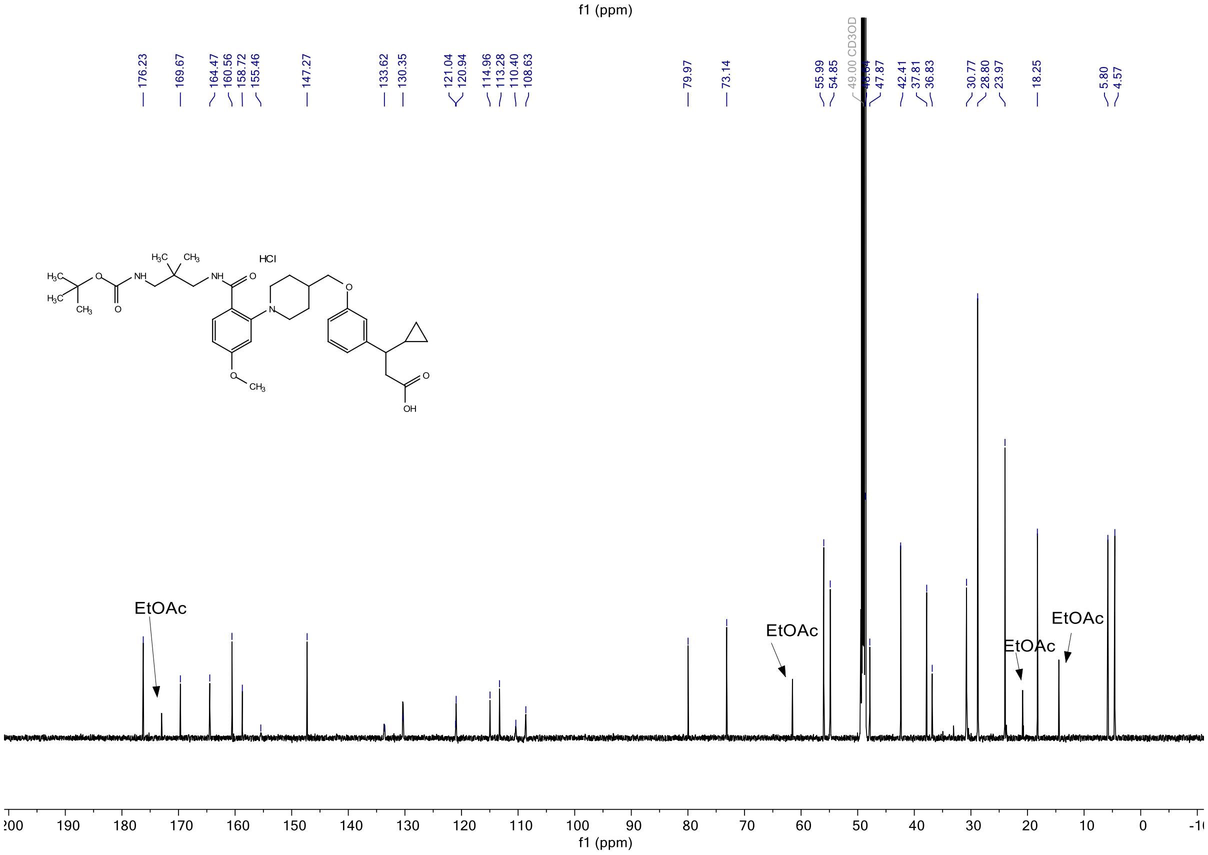
176.23
169.67
164.47
160.56
158.72
155.46
147.27
133.62
130.35
121.04
120.94
114.96
113.28
110.40
108.63
79.97
73.14
55.99
54.85
49.00 CD3OD
47.87
42.41
37.81
36.83
30.77
28.80
23.97
18.25
5.80
4.57
HCl
EtOAc
EtOAc
EtOAc
EtOAc
f1 (ppm)

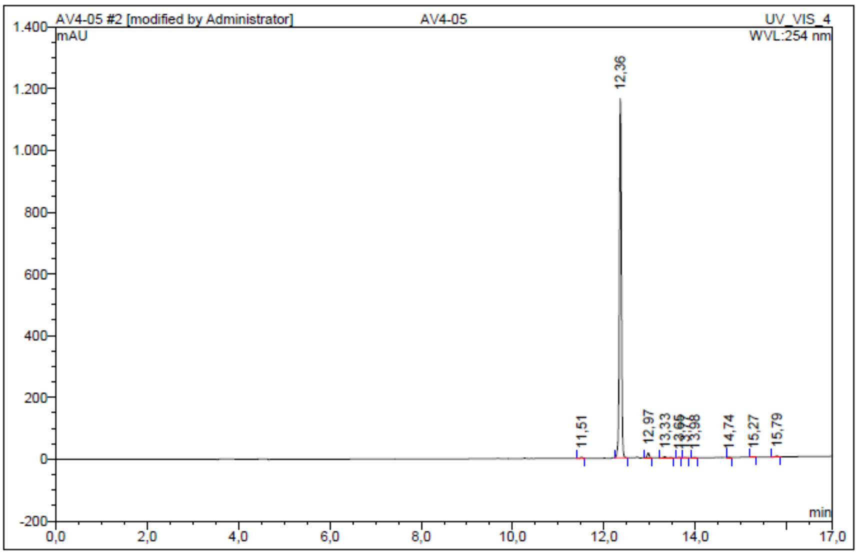


| No. | Ret.Time min | Peak Name | Height mAU | Area mAU*min | Rel.Area % | Amount | Resolution(EP) |
|---|---|---|---|---|---|---|---|
| 1 | 11,51 | n.a. | 4,062 | 0,216 | 0,34 | n.a. | 10,48 |
| 2 | 12,36 | n.a. | 1163,883 | 60,954 | 97,06 | n.a. | 7,64 |
| 3 | 12,97 | n.a. | 16,364 | 0,837 | 1,33 | n.a. | 4,37 |
| 4 | 13,33 | n.a. | 3,824 | 0,335 | 0,53 | n.a. | 3,74 |
| 5 | 13,65 | n.a. | 1,322 | 0,066 | 0,11 | n.a. | 1,53 |
| 6 | 13,77 | n.a. | 0,938 | 0,051 | 0,08 | n.a. | 2,25 |
| 7 | 13,98 | n.a. | 0,831 | 0,055 | 0,09 | n.a. | 8,02 |
| 8 | 14,74 | n.a. | 0,599 | 0,032 | 0,05 | n.a. | 5,82 |
| 9 | 15,27 | n.a. | 0,751 | 0,044 | 0,07 | n.a. | 5,57 |
| 10 | 15,79 | n.a. | 3,433 | 0,213 | 0,34 | n.a. | n.a. |
| Total: | | | 1196,007 | 62,801 | 100,00 | 0,000 | |

***Figure S20.*** $^{1}$H NMR, $^{13}$C NMR and HPLC chromatogram of **TUG-2457 - 14e**.

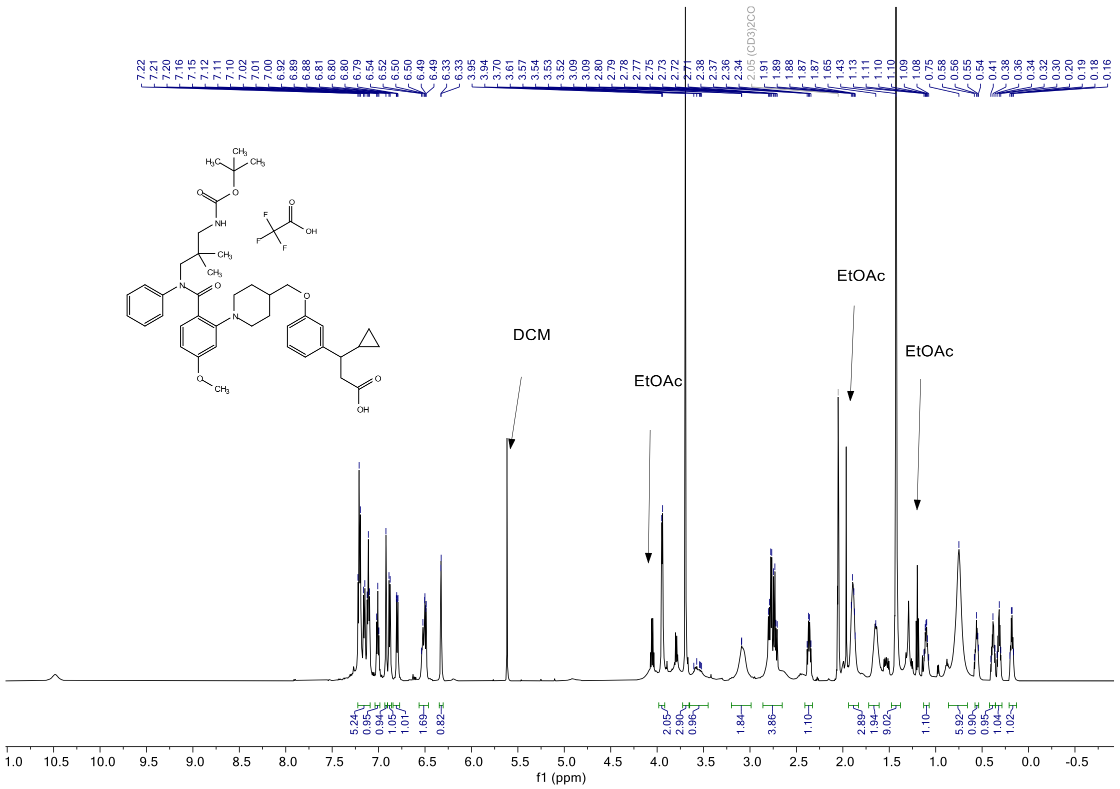

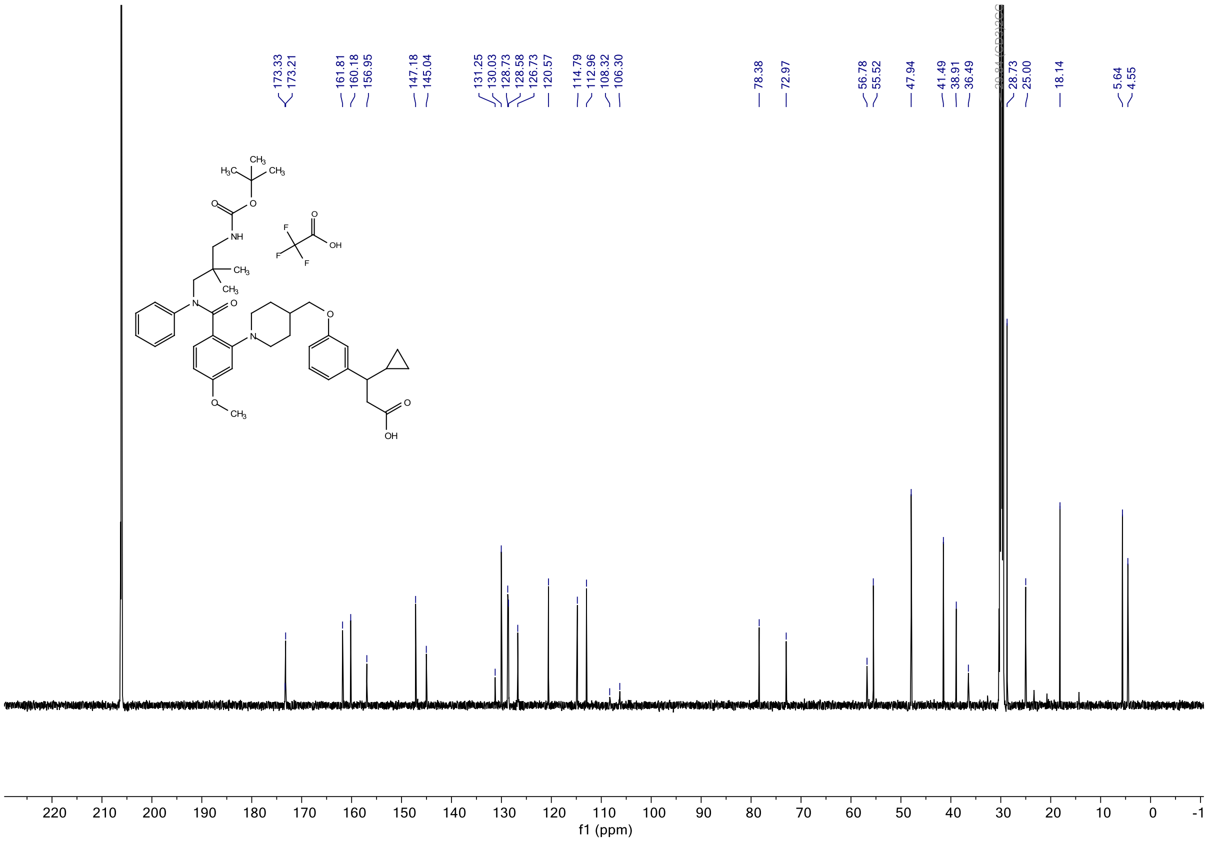


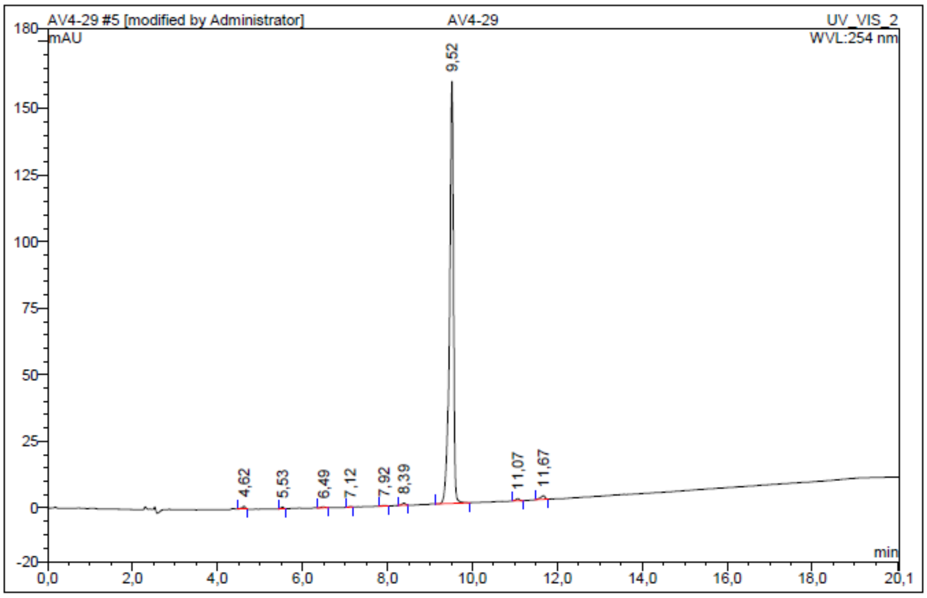


| No. | Ret.Time min | Peak Name | Height mAU | Area mAU*min | Rel.Area % | Amount | Resolution(EP) |
|---|---|---|---|---|---|---|---|
| 1 | 4,62 | n.a. | 1,086 | 0,102 | 0,58 | n.a. | 7,28 |
| 2 | 5,53 | n.a. | 0,604 | 0,042 | 0,24 | n.a. | 5,82 |
| 3 | 6,49 | n.a. | 0,263 | 0,033 | 0,19 | n.a. | 3,78 |
| 4 | 7,12 | n.a. | 0,217 | 0,015 | 0,08 | n.a. | 5,29 |
| 5 | 7,92 | n.a. | 0,175 | 0,020 | 0,11 | n.a. | 2,99 |
| 6 | 8,39 | n.a. | 0,848 | 0,073 | 0,41 | n.a. | 7,87 |
| 7 | 9,52 | n.a. | 158,301 | 17,214 | 97,04 | n.a. | 10,68 |
| 8 | 11,07 | n.a. | 0,858 | 0,075 | 0,42 | n.a. | 3,76 |
| 9 | 11,67 | n.a. | 1,361 | 0,166 | 0,94 | n.a. | n.a. |
| Total: | | | 163,713 | 17,740 | 100,00 | 0,000 | |

***Figure S21.*** $^{1}$H NMR, $^{13}$C NMR and HPLC chromatogram of **TUG-2467 - 14f**.

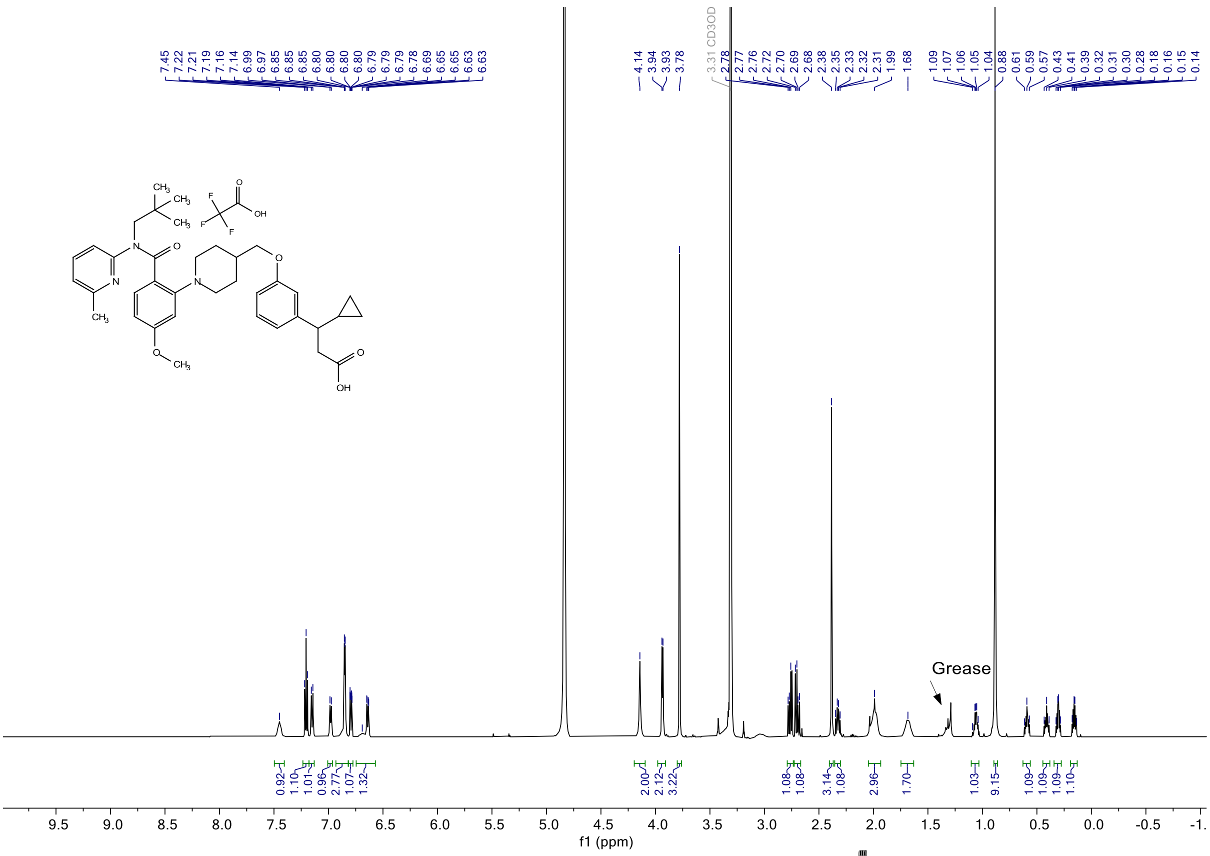
7.45 7.22 7.21 7.19 7.16 7.14 6.99 6.97 6.85 6.85 6.85 6.80 6.80 6.80 6.80 6.80 6.79 6.79 6.78 6.69 6.65 6.65 6.63 6.63
4.14 3.94 3.93 3.78
3.31 CD3OD
2.78 2.77 2.76 2.72 2.70 2.69 2.68 2.38 2.35 2.33 2.32 2.31 1.99 1.68
1.09 1.07 1.06 1.05 1.04 0.88 0.61 0.59 0.57 0.43 0.41 0.39 0.32 0.31 0.30 0.28 0.18 0.16 0.15 0.14
Grease
0.92 1.10 1.01 0.96 2.77 1.07 1.32
2.00 2.12 3.22
1.08 1.08 3.14 1.08 2.96 1.70
1.03 9.15 1.09 1.09 1.09 1.10
f1 (ppm)

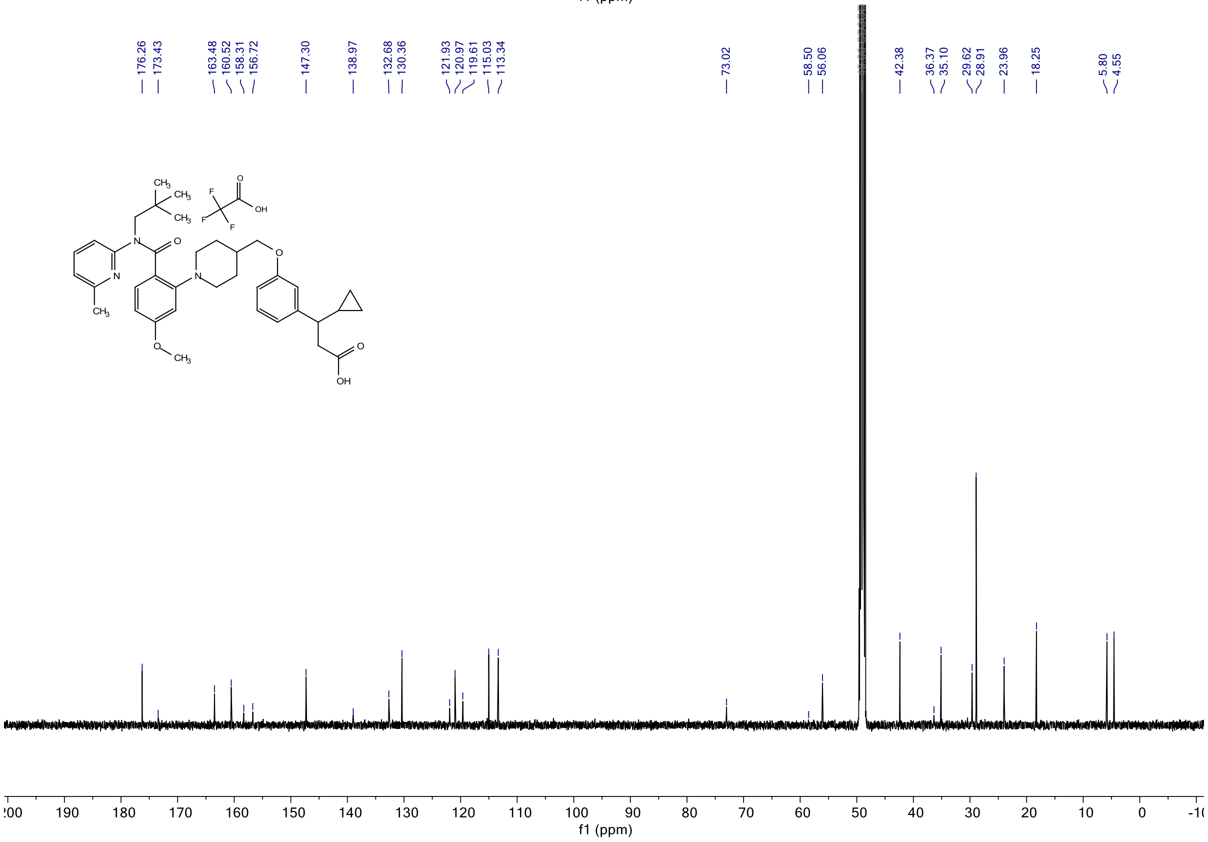
176.26 173.43 163.48 160.52 158.31 156.72 147.30 138.97 132.68 130.36 121.93 120.97 119.61 115.03 113.34
73.02 58.50 56.06 42.38 36.37 35.10 29.62 28.91 23.96 18.25 5.80 4.55
f1 (ppm)

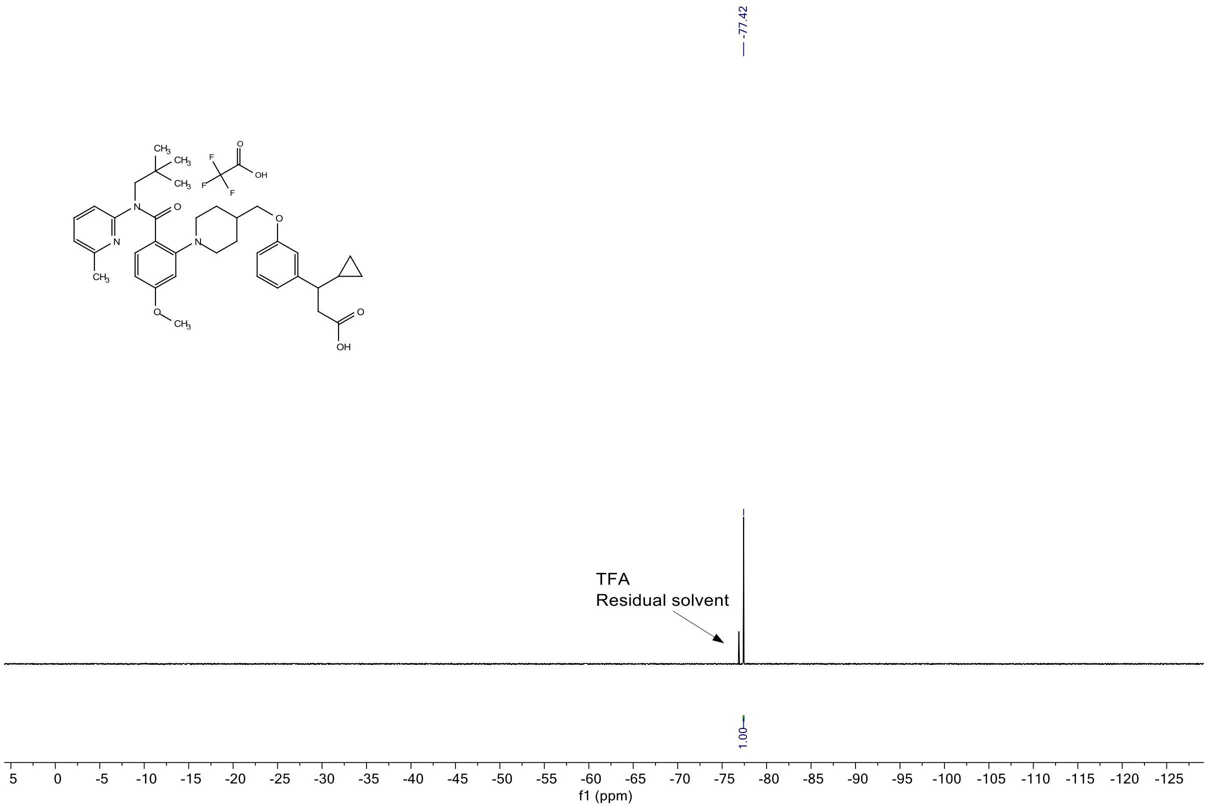


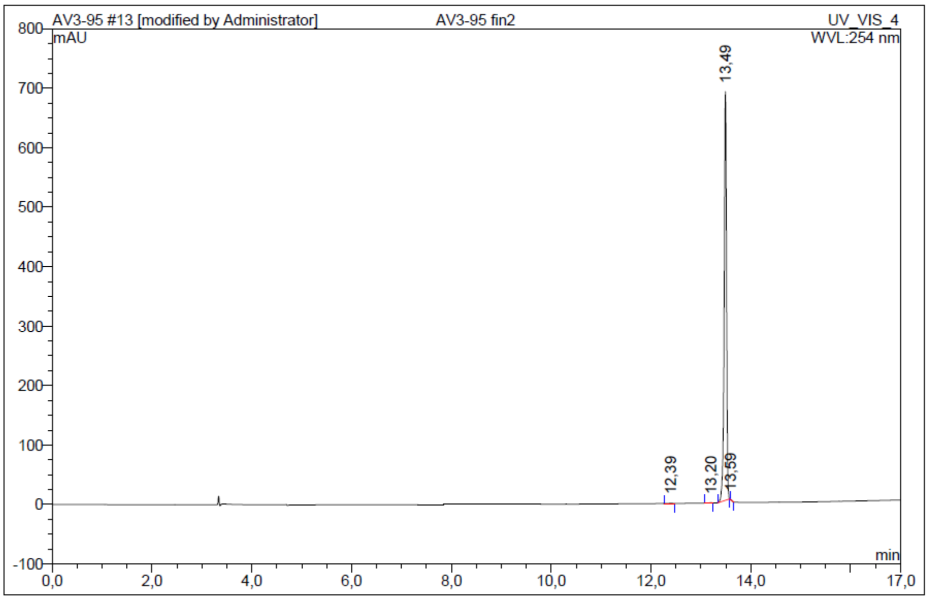


| No. | Ret.Time min | Peak Name | Height mAU | Area mAU*min | Rel.Area % | Amount | Resolution(EP) |
|---|---|---|---|---|---|---|---|
| 1 | 12,39 | n.a. | 0,430 | 0,028 | 0,07 | n.a. | 9,07 |
| 2 | 13,20 | n.a. | 0,349 | 0,023 | 0,06 | n.a. | 3,52 |
| 3 | 13,49 | n.a. | 687,422 | 38,997 | 99,83 | n.a. | n.a. |
| 4 | 13,59 | n.a. | 0,096 | 0,016 | 0,04 | n.a. | n.a. |
| Total: | | | 688,297 | 39,064 | 100,00 | 0,000 | |

***Figure S22.*** $^{1}$H NMR, $^{13}$C NMR and HPLC chromatogram of **Cpd 6h**.

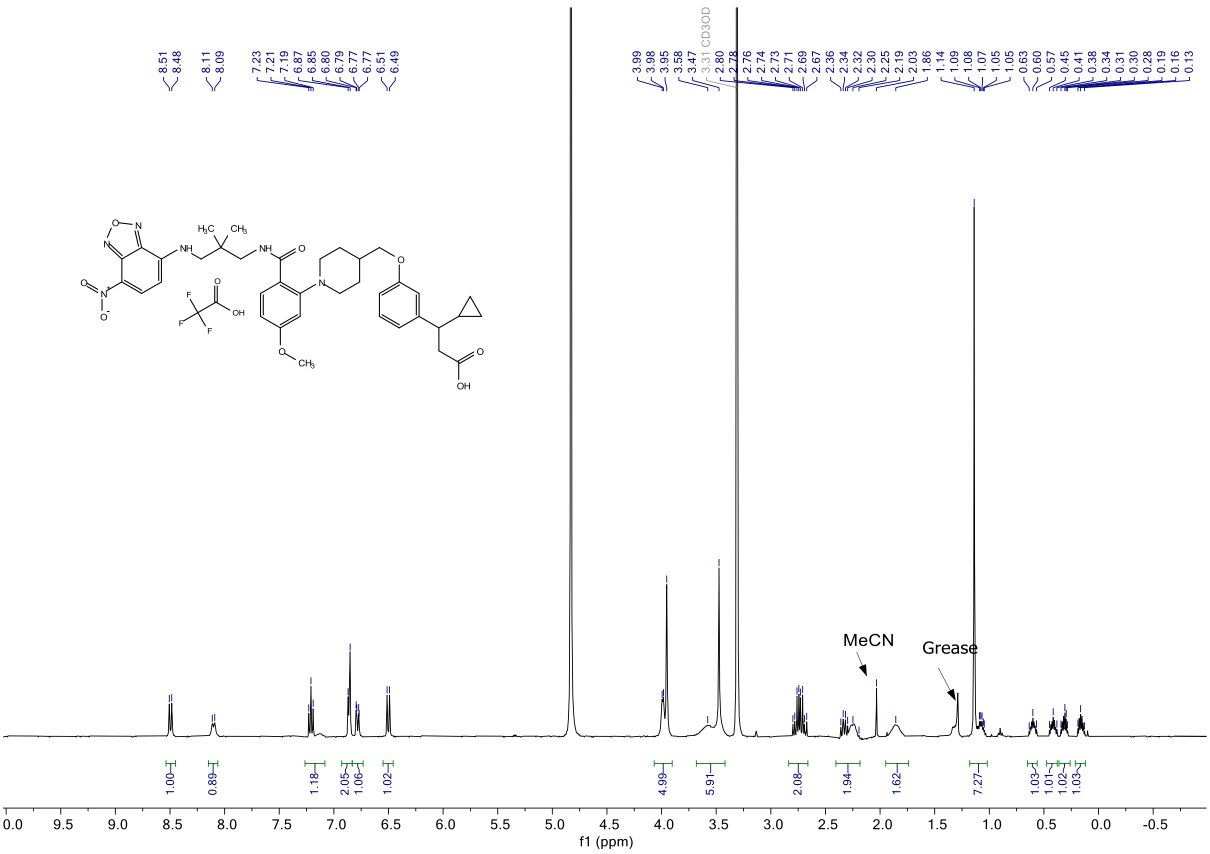
8.51
8.48
8.11
8.09
7.23
7.21
7.19
6.87
6.85
6.80
6.79
6.77
6.77
6.51
6.49
3.99
3.98
3.95
3.58
3.47
3.31 CD3OD
2.80
2.78
2.76
2.74
2.73
2.71
2.69
2.67
2.36
2.34
2.32
2.30
2.25
2.19
2.03
1.86
1.14
1.09
1.08
1.07
1.05
1.05
0.63
0.60
0.57
0.45
0.41
0.38
0.34
0.31
0.30
0.28
0.19
0.16
0.13
MeCN
Grease
1.00
0.89
1.18
2.05
1.06
1.02
4.99
5.91
2.08
1.94
1.62
7.27
1.03
1.01
1.02
1.03
f1 (ppm)

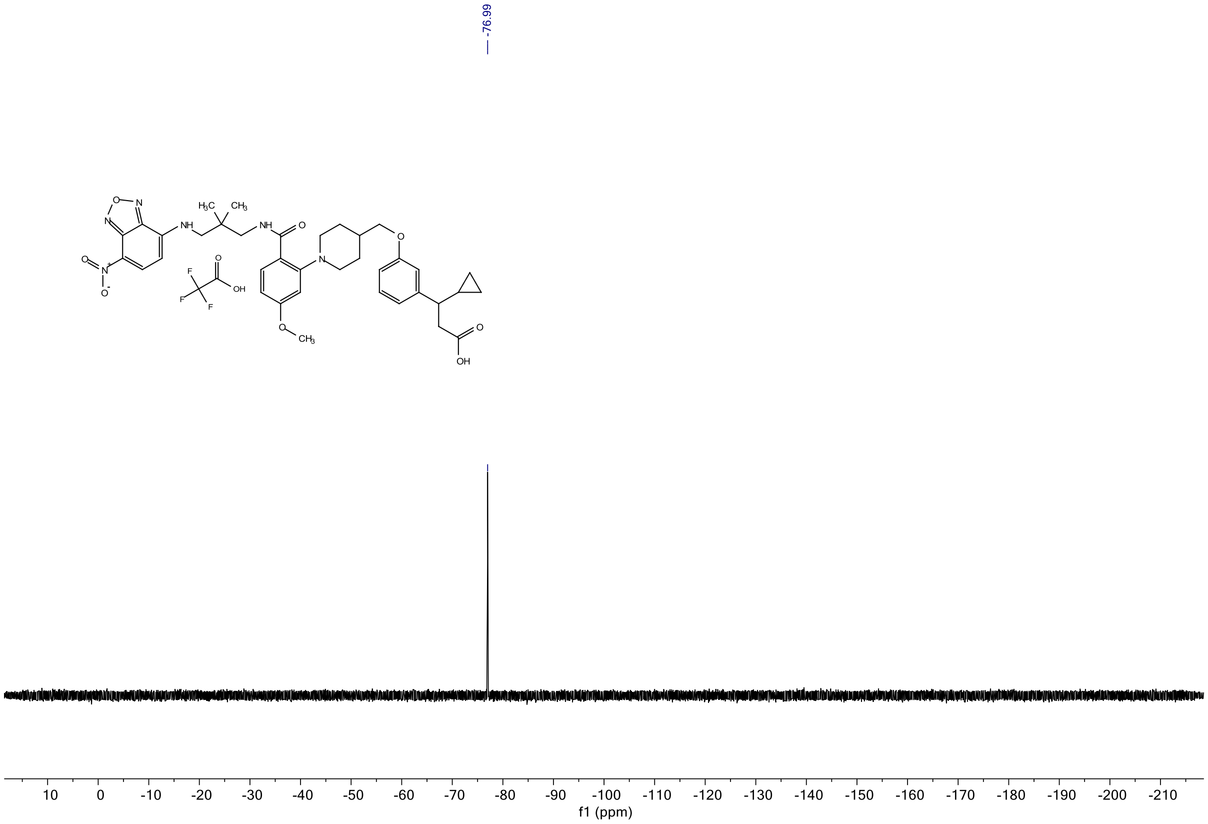
-76.99
f1 (ppm)

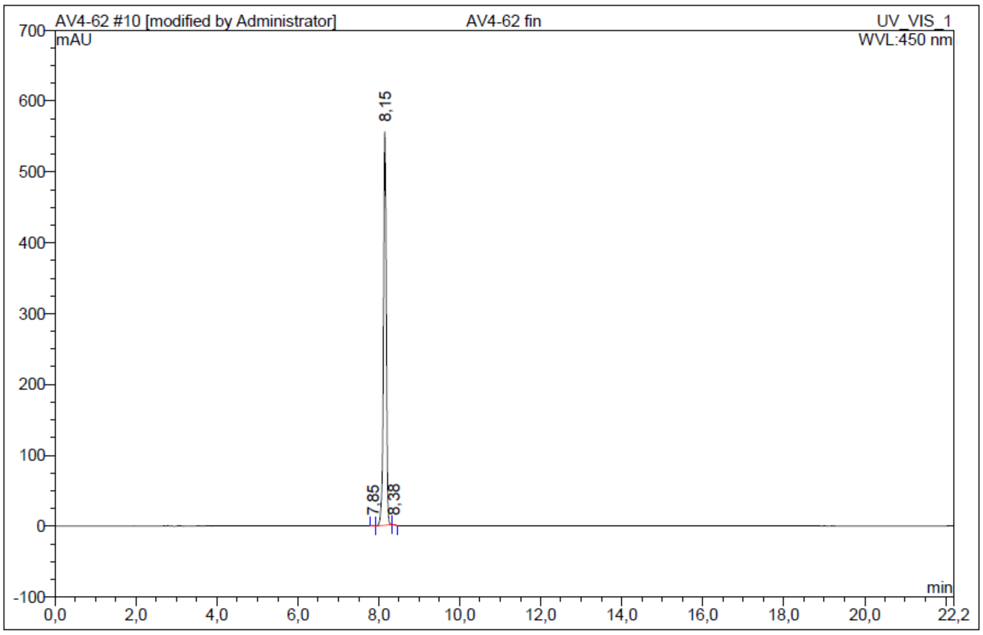

| No. | Ret.Time min | Peak Name | Height mAU | Area mAU*min | Rel.Area % | Amount | Resolution(EP) |
|---|---|---|---|---|---|---|---|
| 1 | 7,85 | n.a. | 0,571 | 0,032 | 0,07 | n.a. | 2,79 |
| 2 | 8,15 | n.a. | 555,724 | 43,288 | 99,82 | n.a. | 2,02 |
| 3 | 8,38 | n.a. | 0,711 | 0,045 | 0,10 | n.a. | n.a. |
| Total: | | | 557,006 | 43,365 | 100,00 | 0,000 | |

***Figure S23***. $^{1}$H NMR, $^{19}$F NMR and HPLC chromatogram of **TUG-2489 - 15**.

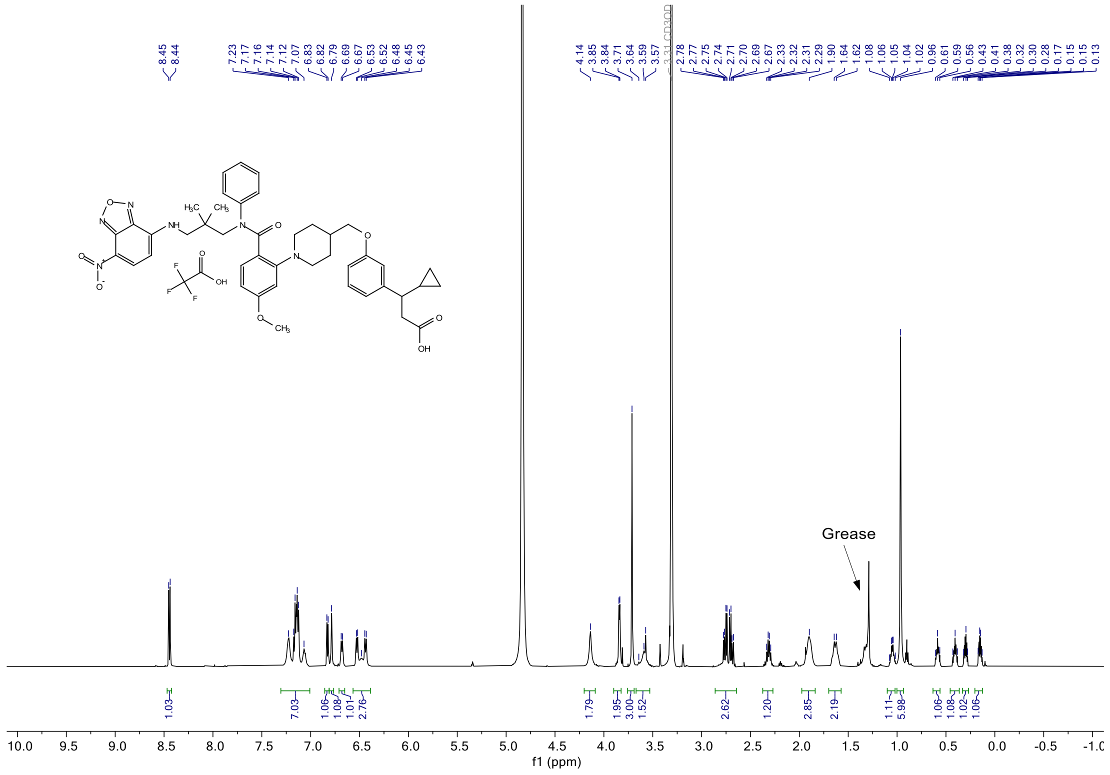

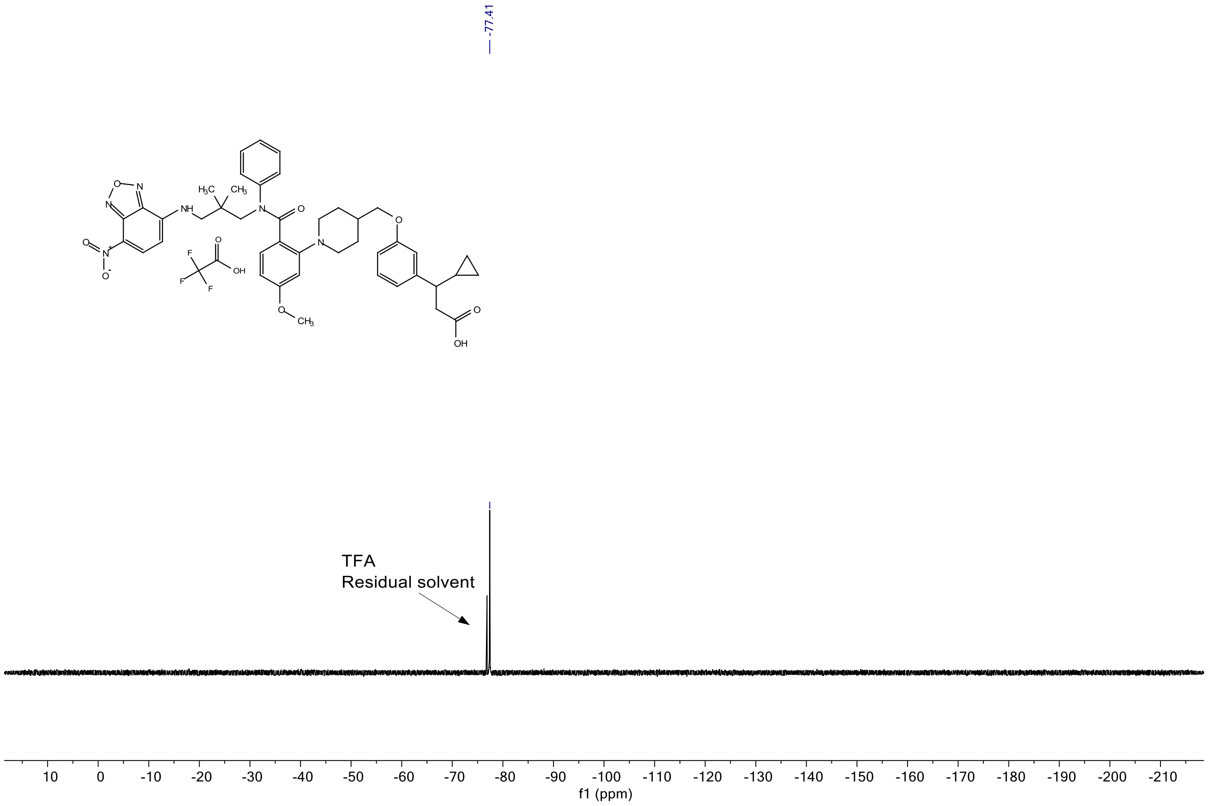


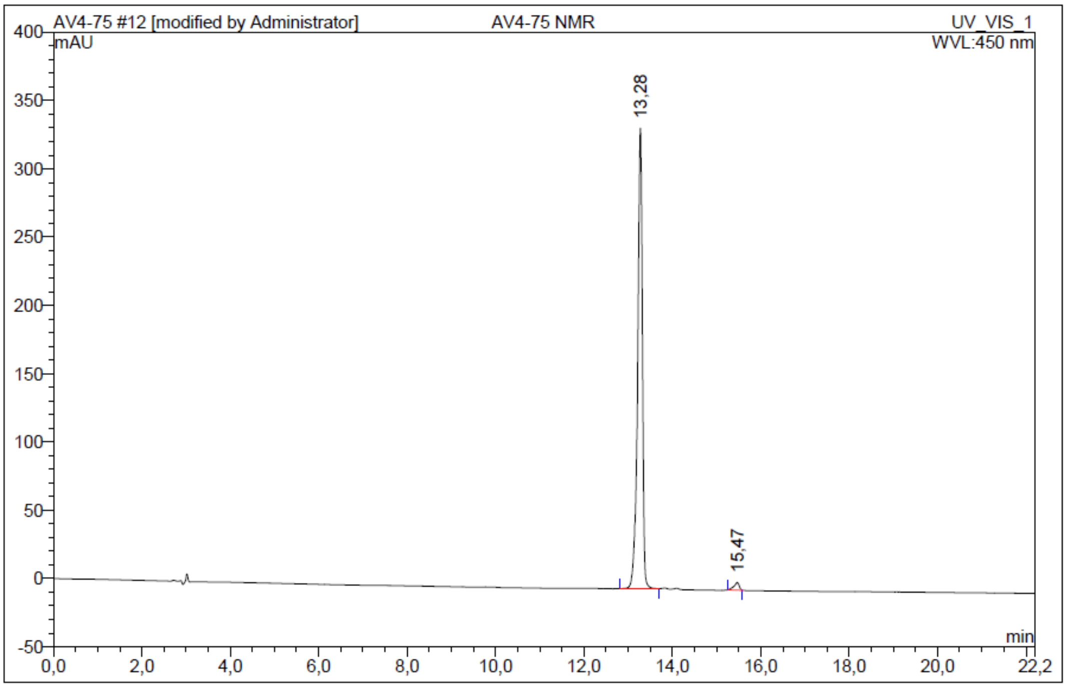


| No. | Ret.Time min | Peak Name | Height mAU | Area mAU*min | Rel.Area % | Amount | Resolution(EP) |
|---|---|---|---|---|---|---|---|
| 1 | 13,28 | n.a. | 337,202 | 42,915 | 98,23 | n.a. | 11,08 |
| 2 | 15,47 | n.a. | 5,595 | 0,773 | 1,77 | n.a. | n.a. |
| Total: | | | 342,796 | 43,688 | 100,00 | 0,000 | |

***Figure S24.*** [1]H NMR, [19]F NMR and HPLC chromatogram of **TUG-2490 - 16**.

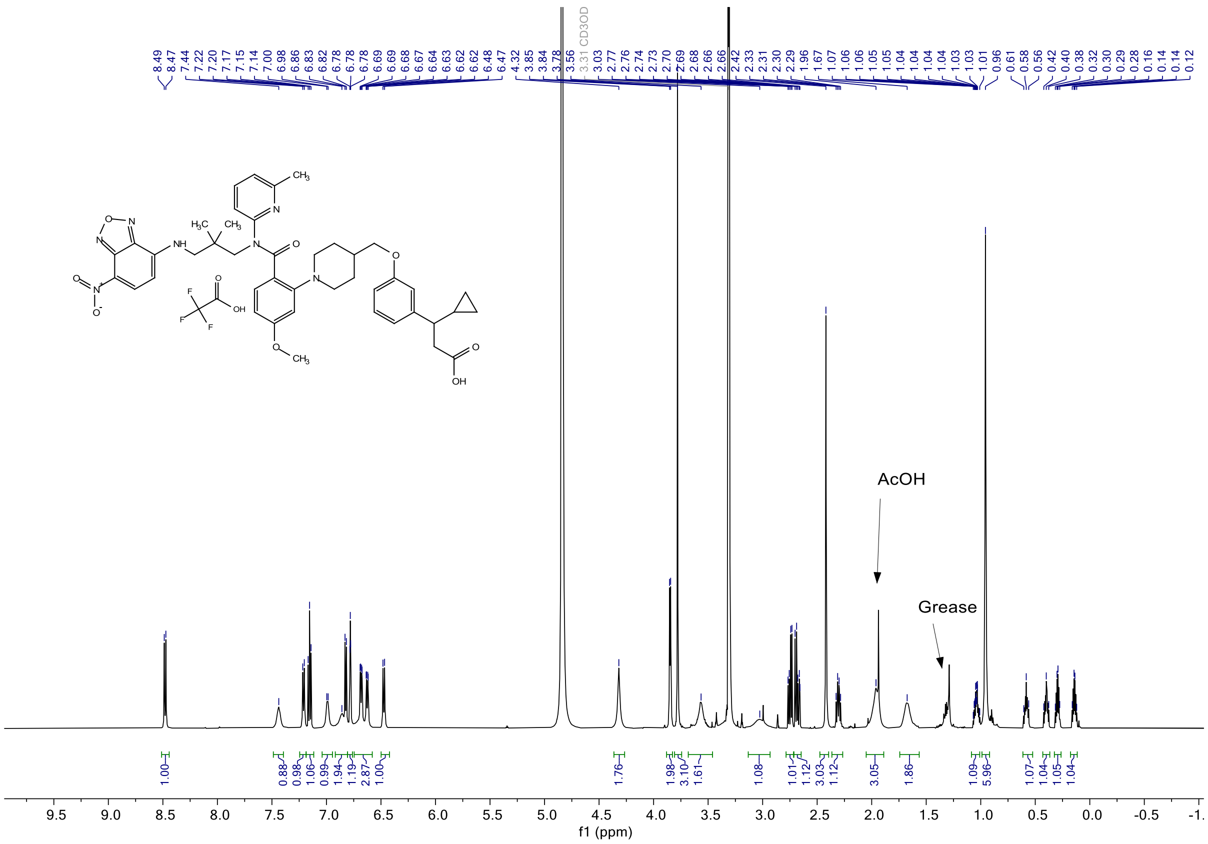
8.49
8.47
7.44
7.22
7.20
7.17
7.15
7.14
7.00
6.98
6.86
6.83
6.82
6.78
6.78
6.78
6.69
6.69
6.68
6.67
6.64
6.63
6.62
6.62
6.48
6.47
4.32
3.85
3.84
3.78
3.56
3.31 CD3OD
3.03
2.77
2.76
2.74
2.73
2.70
2.69
2.68
2.66
2.66
2.42
2.33
2.31
2.30
2.29
1.96
1.67
1.07
1.06
1.06
1.05
1.05
1.04
1.04
1.04
1.04
1.03
1.03
1.01
0.96
0.61
0.58
0.56
0.42
0.40
0.38
0.32
0.30
0.29
0.28
0.16
0.14
0.12
AcOH
Grease
1.00
0.88
0.98
1.06
0.99
1.94
1.19
2.87
1.00
1.76
1.98
3.10
1.61
1.08
1.01
1.12
3.03
1.12
3.05
1.86
1.09
5.96
1.07
1.04
1.05
1.04
f1 (ppm)

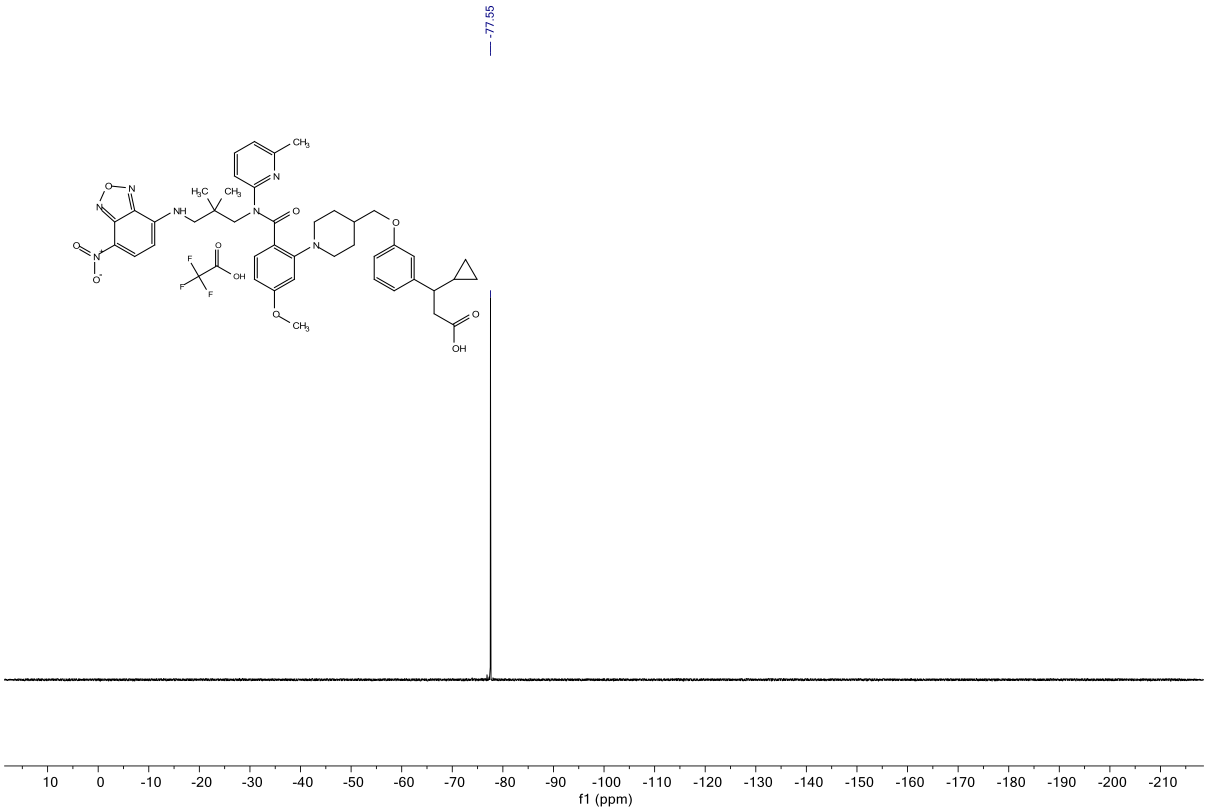
-77.55
f1 (ppm)

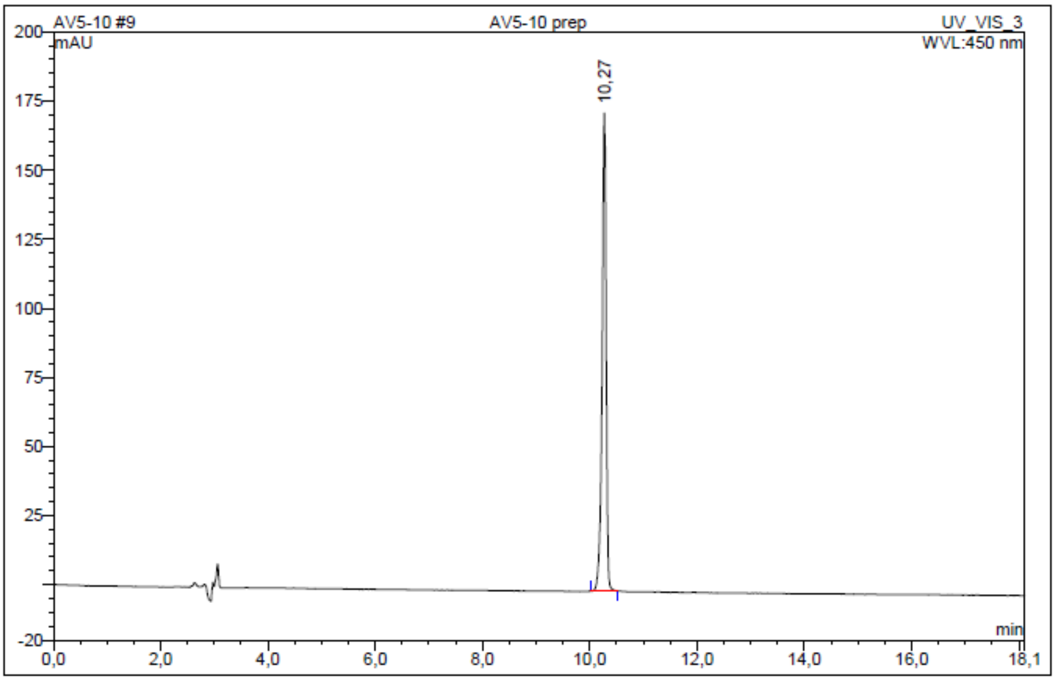

| No. | Ret.Time min | Peak Name | Height mAU | Area mAU*min | Rel.Area % | Amount | Resolution(EP) |
|---|---|---|---|---|---|---|---|
| 1 | 10,27 | n.a. | 172,973 | 14,863 | 100,00 | n.a. | n.a. |
| Total: | | | 172,973 | 14,863 | 100,00 | 0,000 | |

***Figure S25***. $^1$H NMR, $^{19}$F NMR and HPLC chromatogram of **TUG-2597 - 19**.

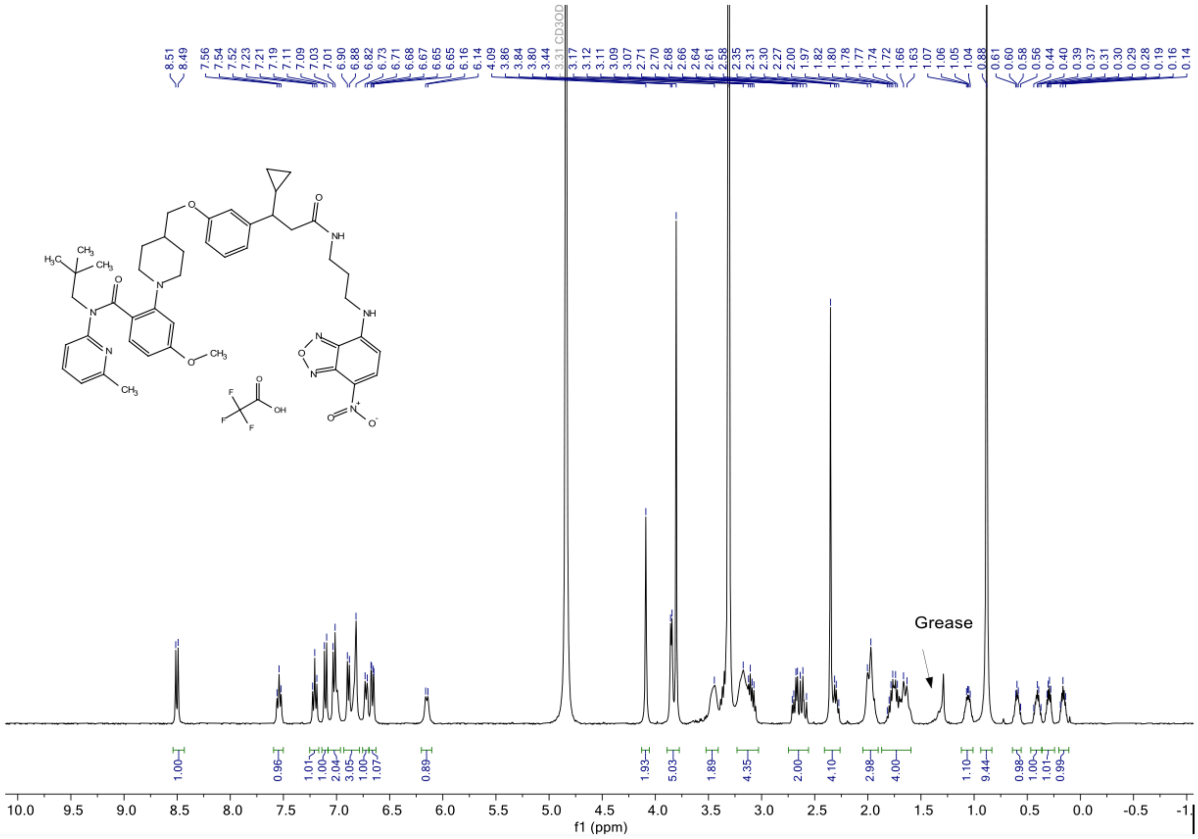

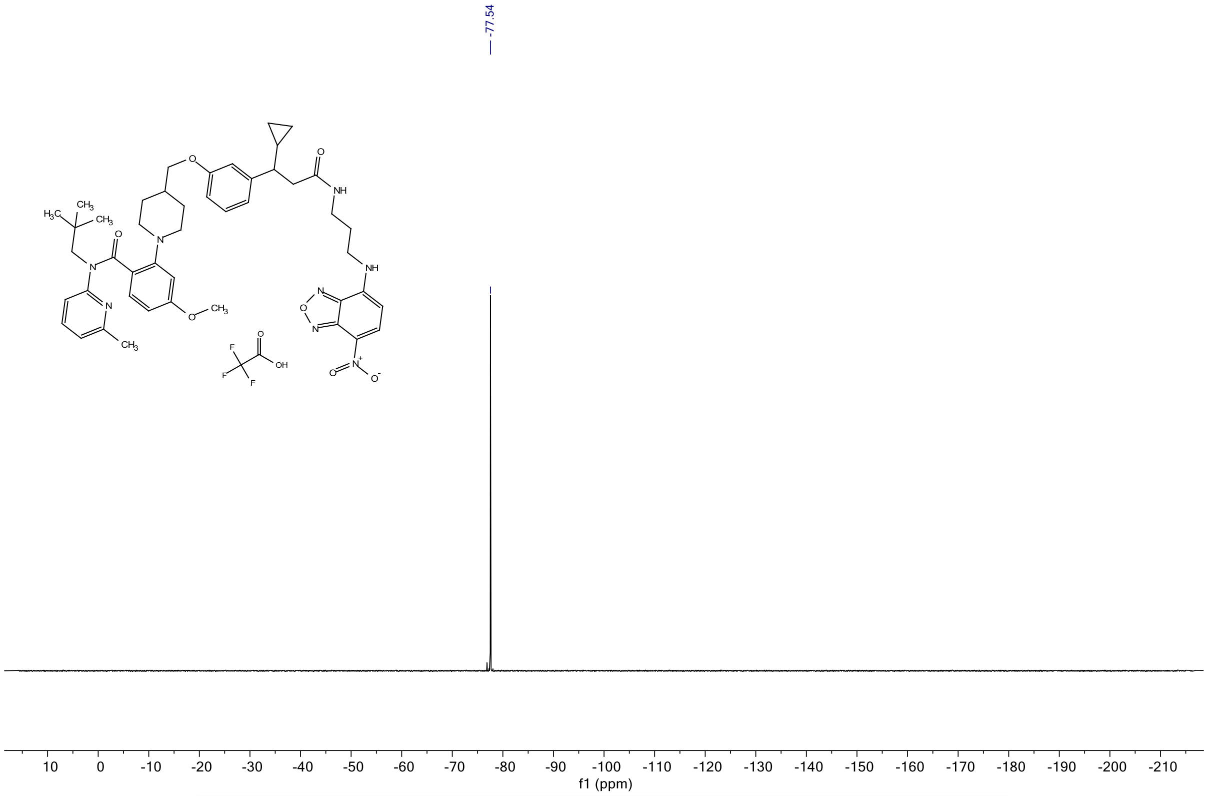


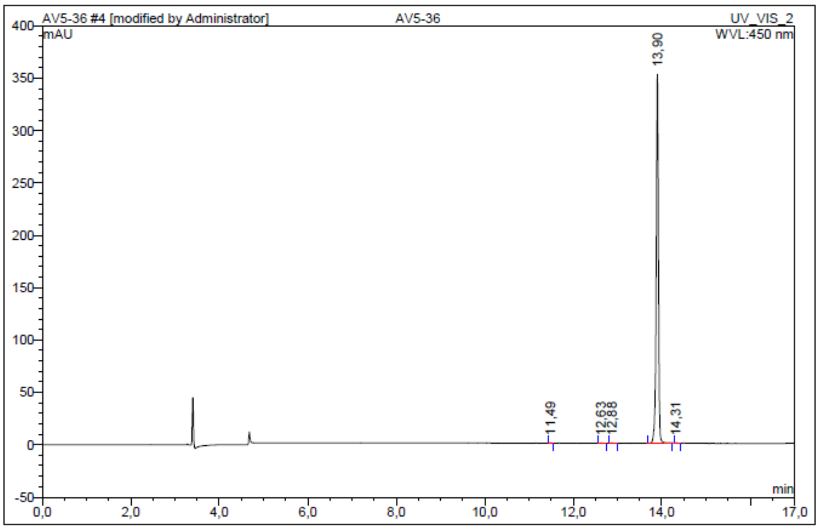


| No. | Ret.Time min | Peak Name | Height mAU | Area mAU*min | Rel.Area % | Amount | Resolution(EP) |
|---|---|---|---|---|---|---|---|
| 1 | 11,49 | n.a. | 0,472 | 0,022 | 0,10 | n.a. | 13,66 |
| 2 | 12,63 | n.a. | 0,239 | 0,015 | 0,07 | n.a. | 2,61 |
| 3 | 12,88 | n.a. | 0,257 | 0,016 | 0,08 | n.a. | 10,69 |
| 4 | 13,90 | n.a. | 352,173 | 21,283 | 99,71 | n.a. | 4,78 |
| 5 | 14,31 | n.a. | 0,194 | 0,009 | 0,04 | n.a. | n.a. |
| Total: | | | 353,336 | 21,345 | 100,00 | 0,000 | |

***Figure S26***. $^{1}$H NMR, $^{19}$F NMR and HPLC chromatogram of **TUG-2598 - 20**.

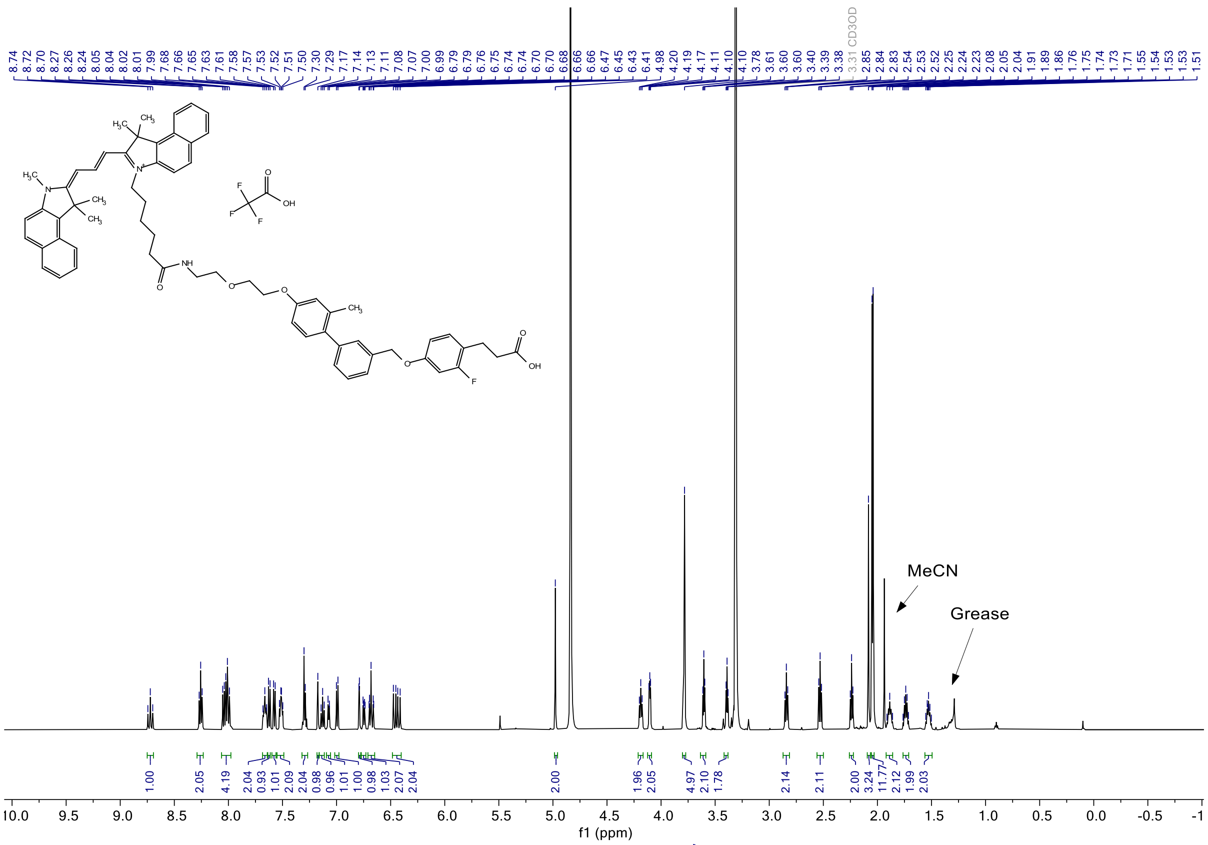
MeCN
Grease
CD3OD
f1 (ppm)

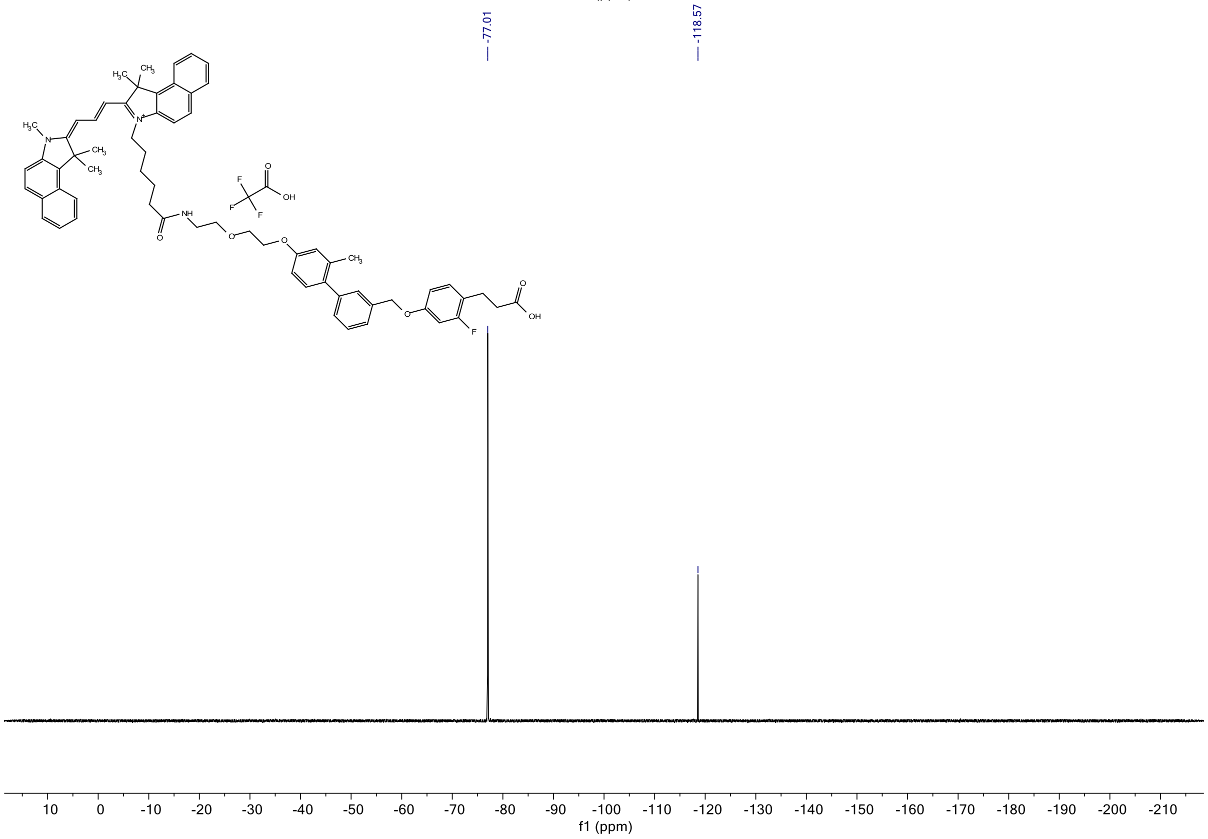
-77.01
-118.57
f1 (ppm)

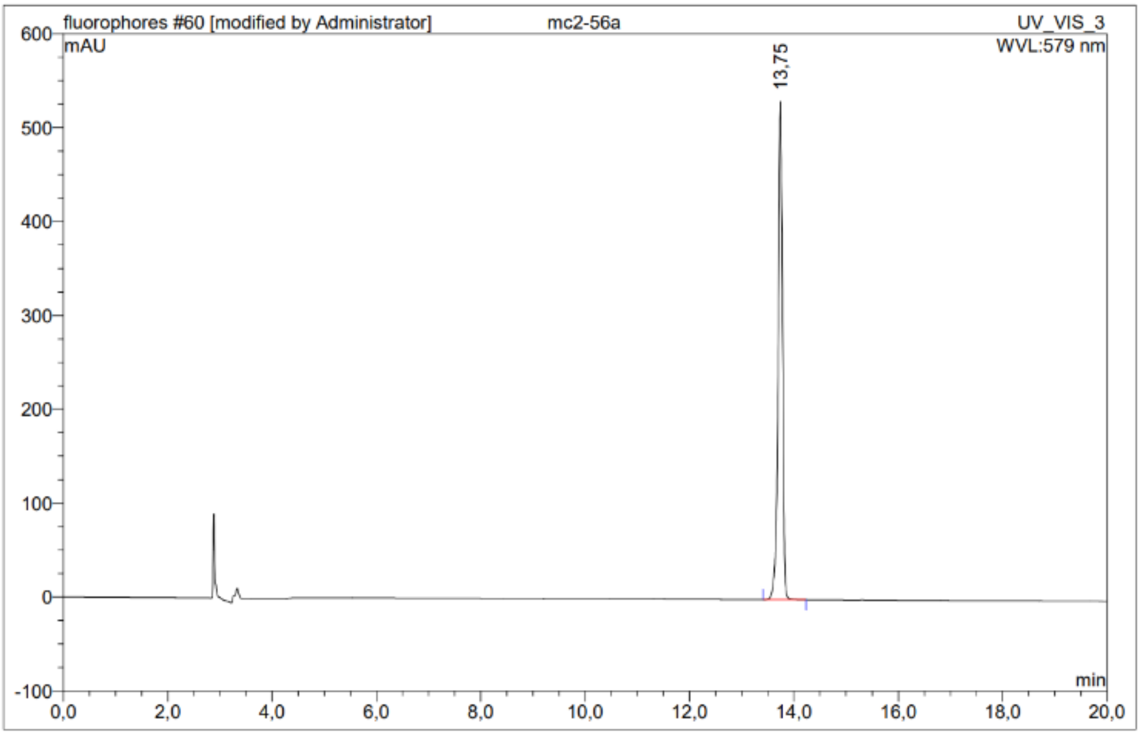


| No. | Ret.Time min | Peak Name | Height mAU | Area mAU*min | Rel.Area % | Amount | Resolution(EP) |
|---|---|---|---|---|---|---|---|
| 1 | 13,75 | n.a. | 530,578 | 50,725 | 100,00 | n.a. | n.a. |
| Total: | | | 530,578 | 50,725 | 100,00 | 0,000 | |

***Figure S27.*** [1]H NMR, [19]F NMR and HPLC chromatogram of **TUG-2287 - 23**.

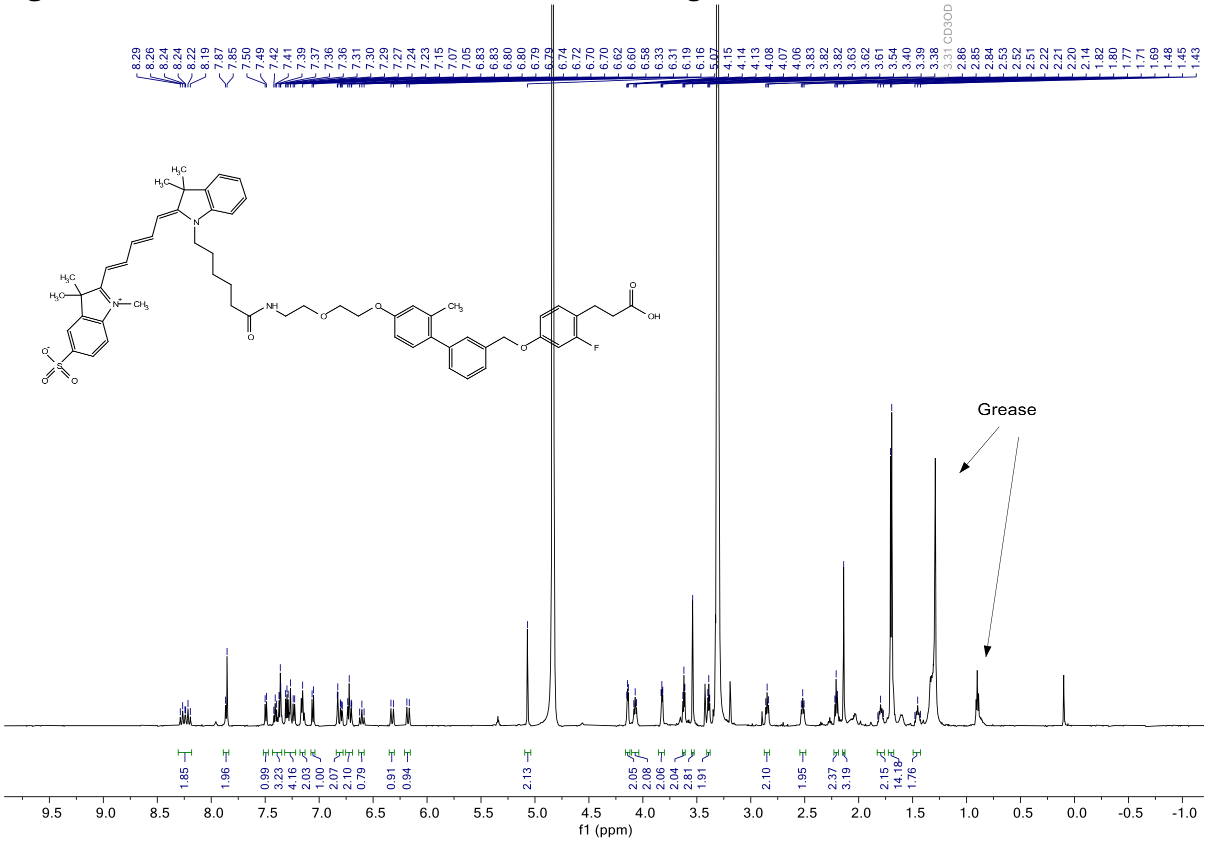

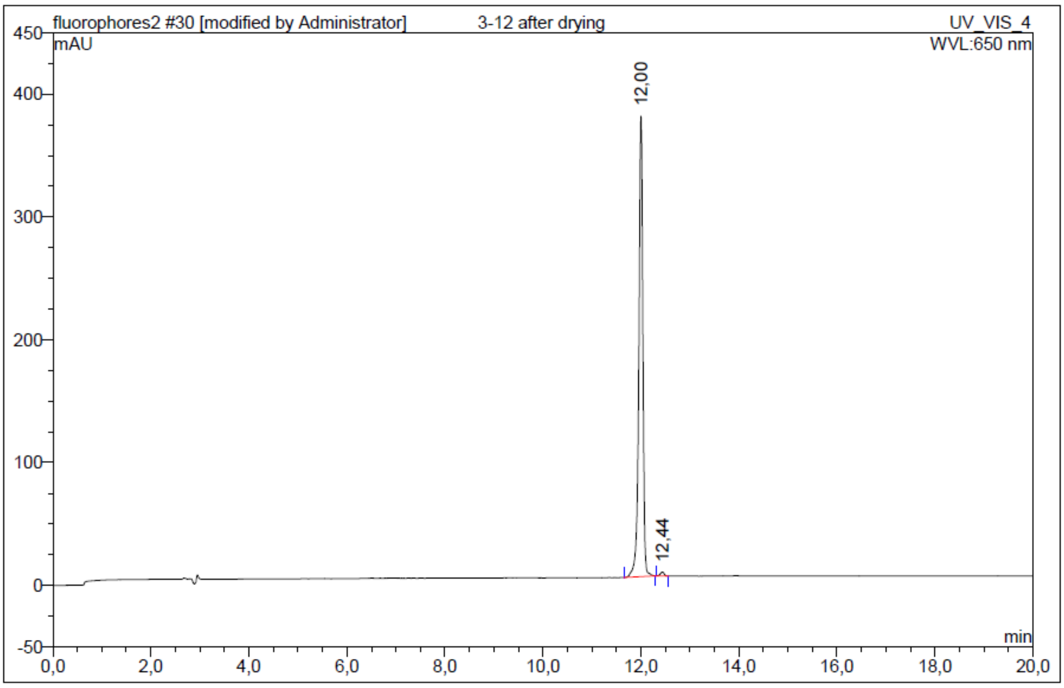


| No. | Ret.Time min | Peak Name | Height mAU | Area mAU*min | Rel.Area % | Amount | Resolution(EP) |
|---|---|---|---|---|---|---|---|
| 1 | 12,00 | n.a. | 374,817 | 36,684 | 99,20 | n.a. | 2,98 |
| 2 | 12,44 | n.a. | 3,226 | 0,296 | 0,80 | n.a. | n.a. |
| Total: | | | 378,043 | 36,980 | 100,00 | 0,000 | |

***Figure S28***. $^1$H NMR and HPLC chromatogram of **TUG-2355 - 24**.

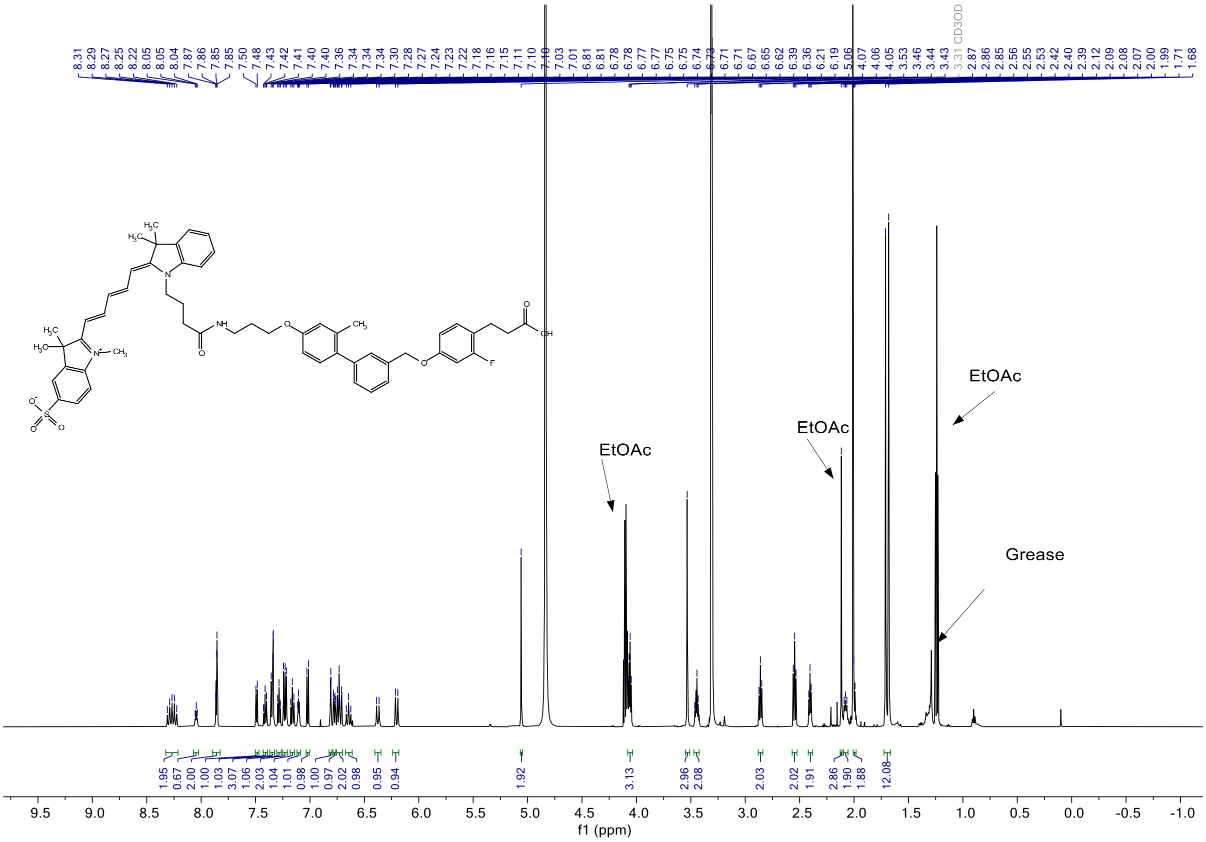

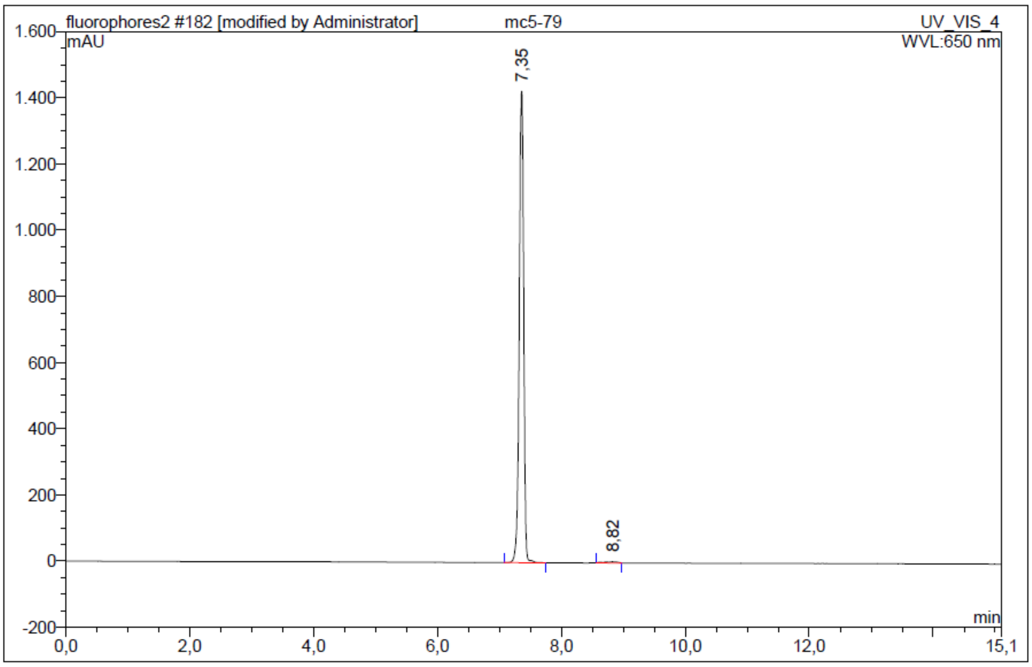


| No. | Ret.Time min | Peak Name | Height mAU | Area mAU*min | Rel.Area % | Amount | Resolution(EP) |
|---|---|---|---|---|---|---|---|
| 1 | 7,35 | n.a. | 1425,063 | 116,383 | 99,33 | n.a. | 4,47 |
| 2 | 8,82 | n.a. | 3,165 | 0,788 | 0,67 | n.a. | n.a. |
| Total: | | | 1428,228 | 117,171 | 100,00 | 0,000 | |

***Figure S29***. $^{1}$H NMR and HPLC chromatogram of **TUG-2591 - 27**.

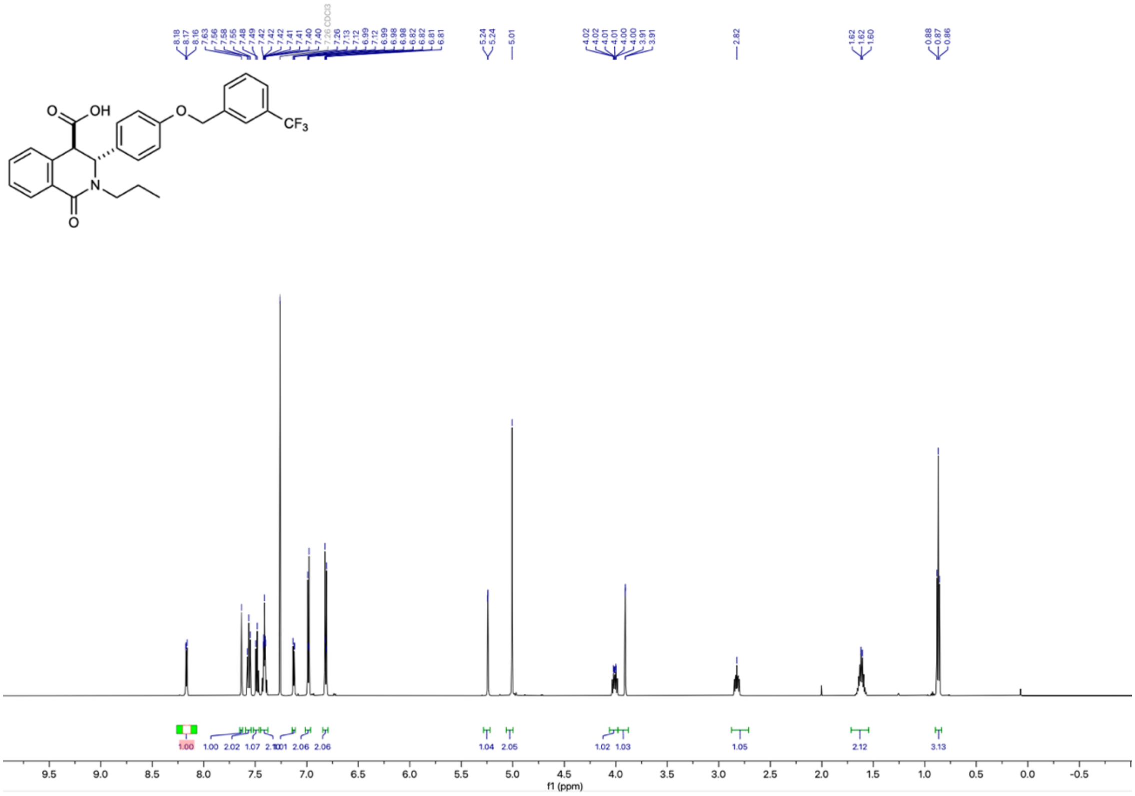

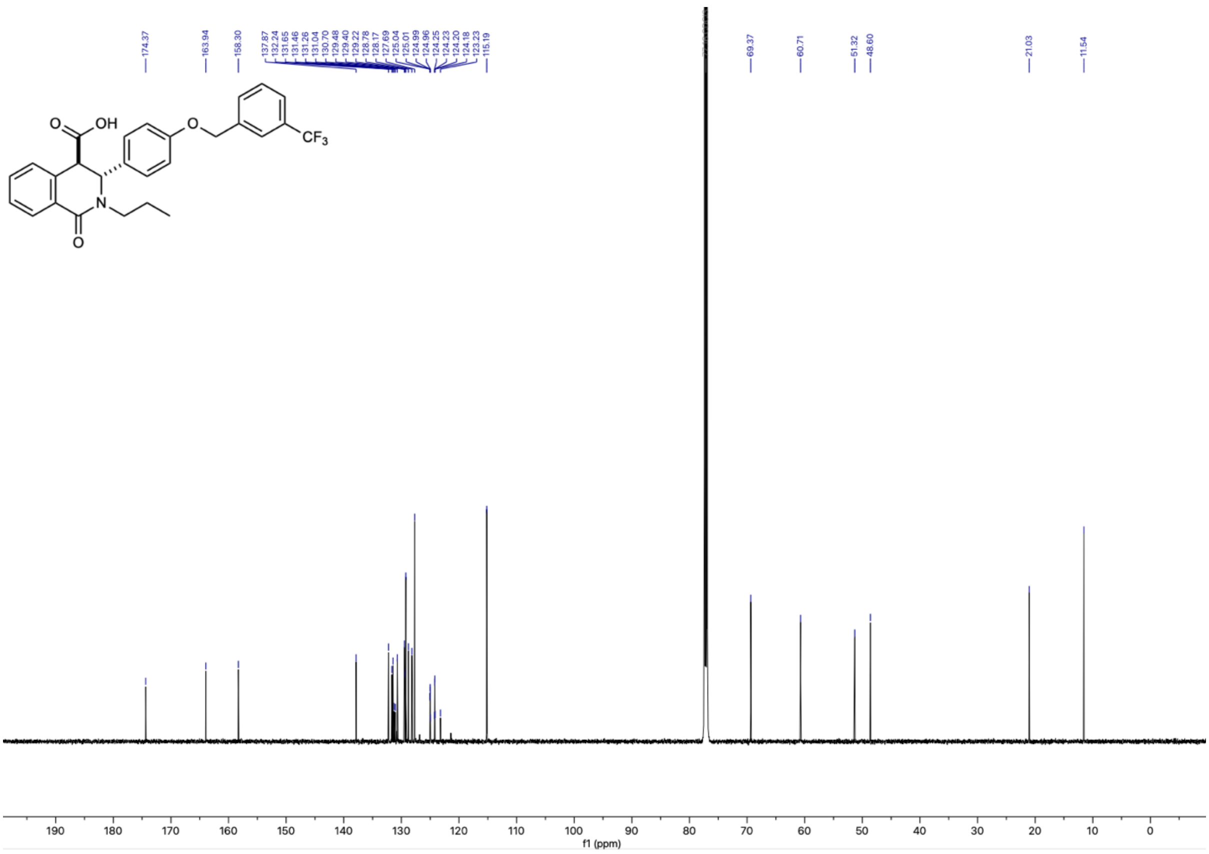
174.37
163.94
158.30
137.87
132.24
131.65
131.46
131.26
131.04
130.70
129.48
129.40
129.22
128.78
128.17
127.69
125.04
125.01
124.99
124.96
124.25
124.23
124.20
124.18
123.23
115.19
69.37
60.71
51.32
48.60
21.03
11.54
OH
O
O
N
O
CF3
f1 (ppm)

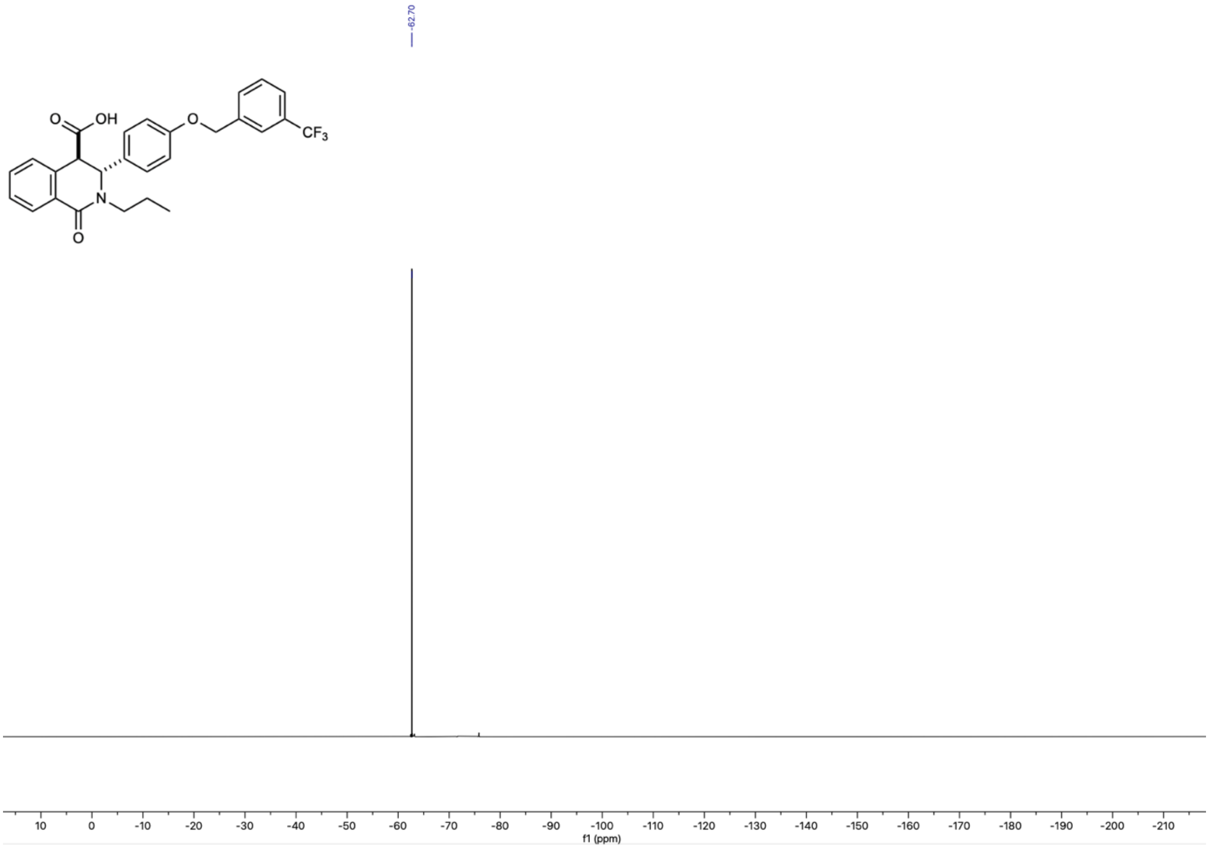
-62.70
OH
O
O
N
O
CF3
f1 (ppm)

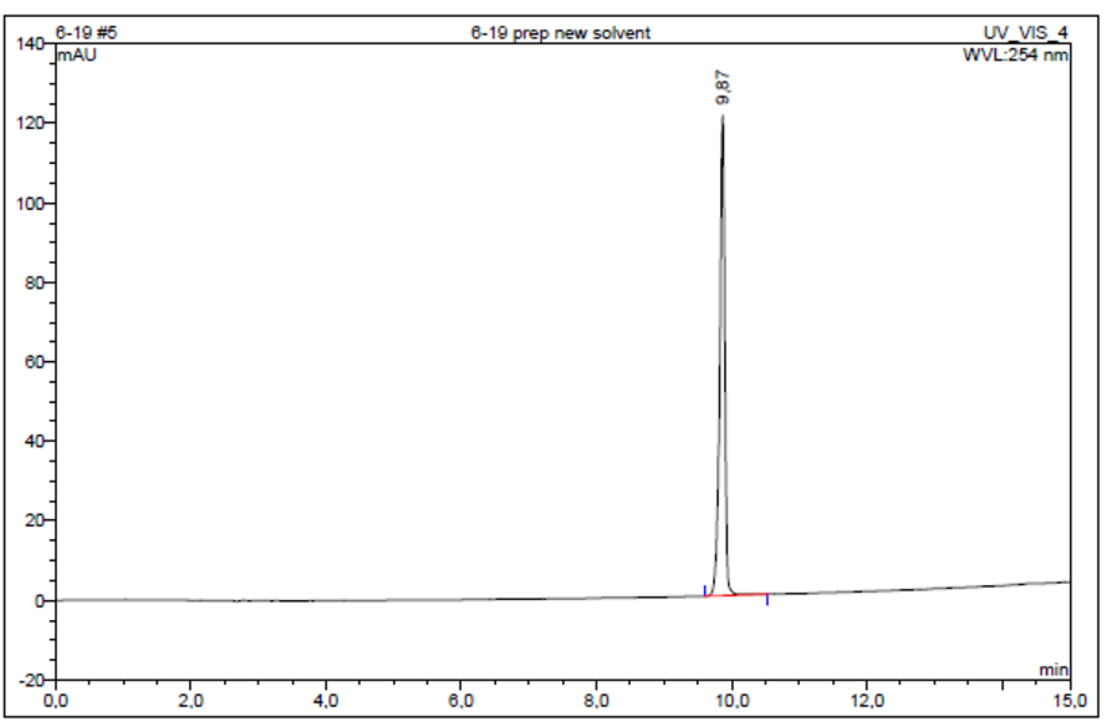


| No. | Ret.Time min | Peak Name | Height mAU | Area mAU*min | Rel.Area % | Amount | Resolution(EP) |
|---|---|---|---|---|---|---|---|
| 1 | 9,87 | n.a. | 120,870 | 10,643 | 100,00 | n.a. | n.a. |
| Total: | | | 120,870 | 10,643 | 100,00 | 0,000 | |

***Figure S30.*** $^{1}$H NMR, $^{13}$C NMR, $^{19}$F NMR and HPLC chromatogram of **TUG-2743**.

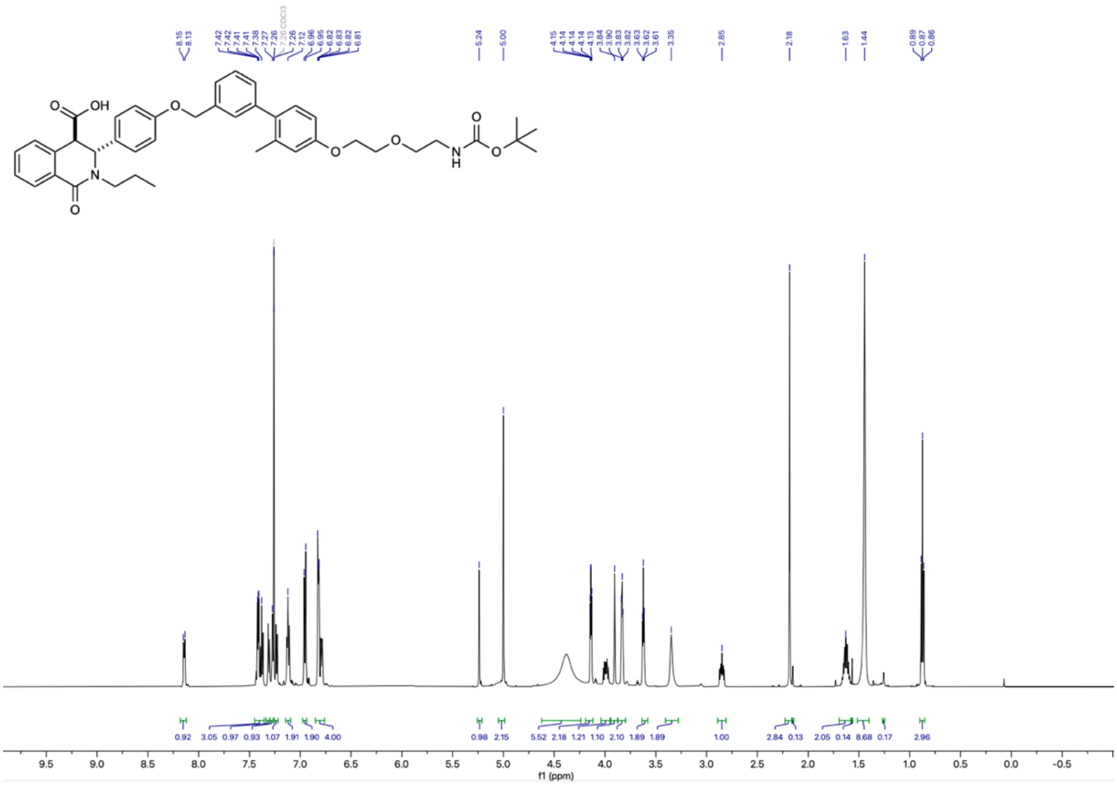

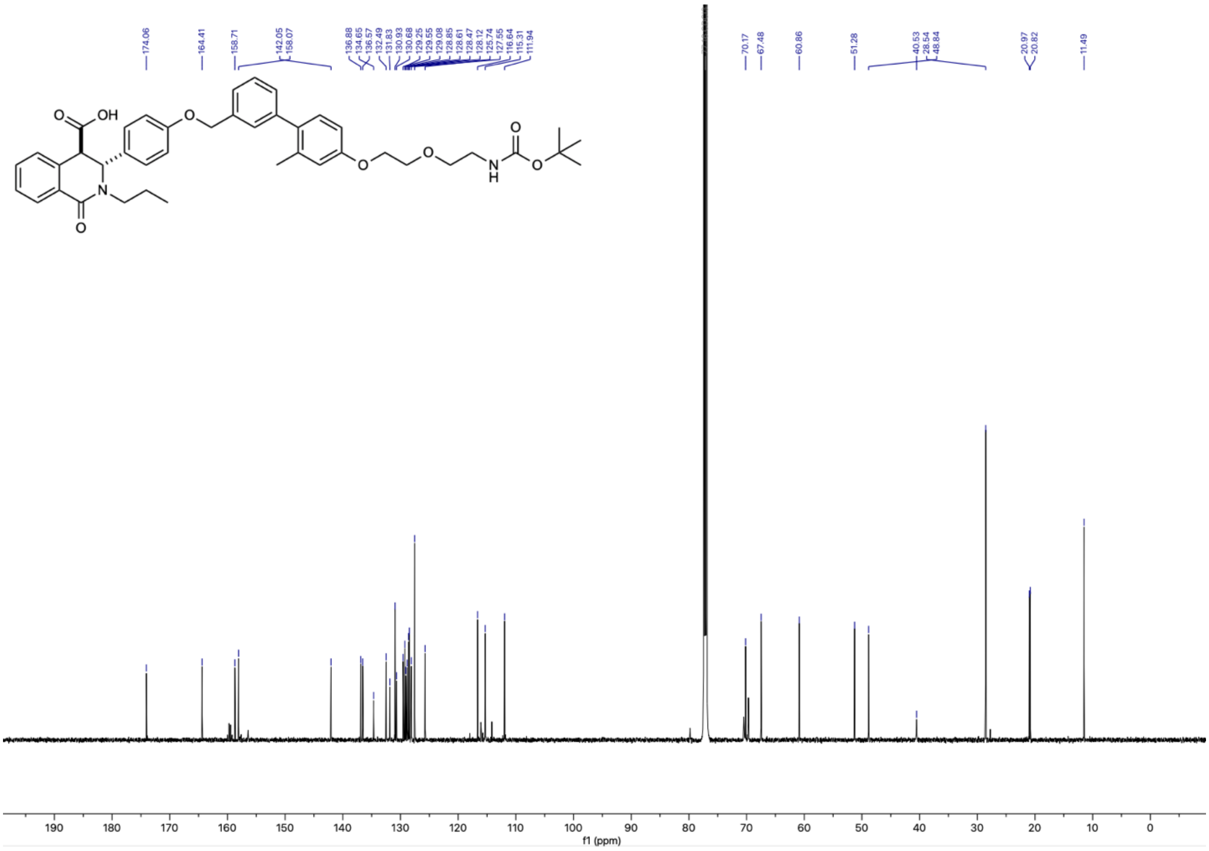


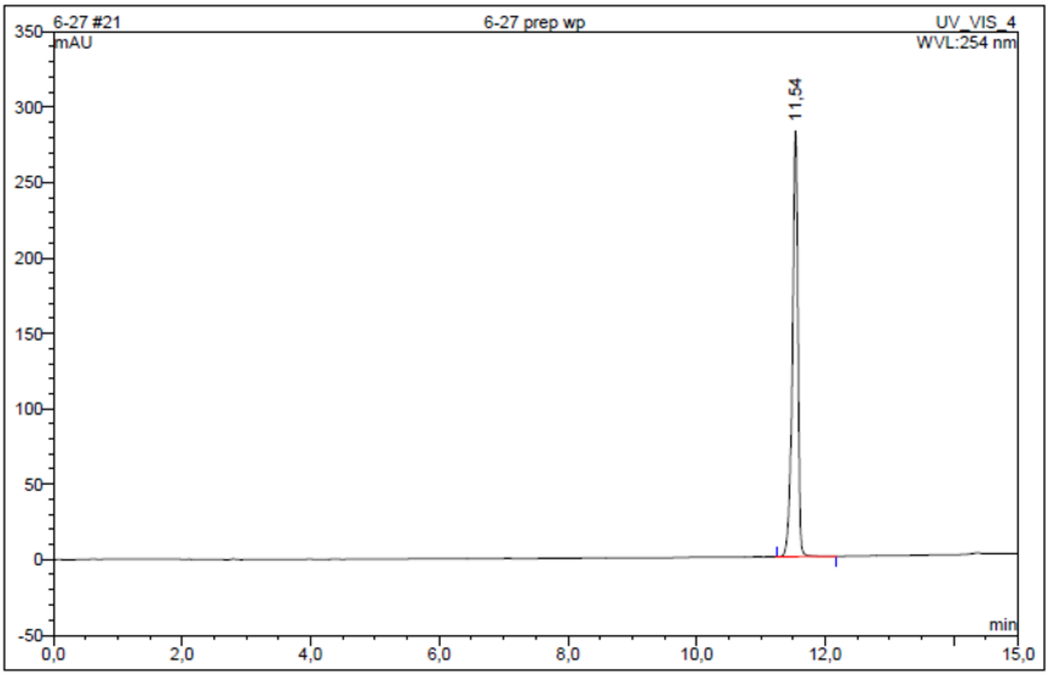


| No. | Ret.Time min | Peak Name | Height mAU | Area mAU*min | Rel.Area % | Amount | Resolution(EP) |
|---|---|---|---|---|---|---|---|
| 1 | 11,54 | n.a. | 282,307 | 28,393 | 100,00 | n.a. | n.a. |
| Total: | | | 282,307 | 28,393 | 100,00 | 0,000 | |

***Figure S31.*** $^{1}H$ NMR, $^{13}C$ NMR and HPLC chromatogram of **TUG-2744**.

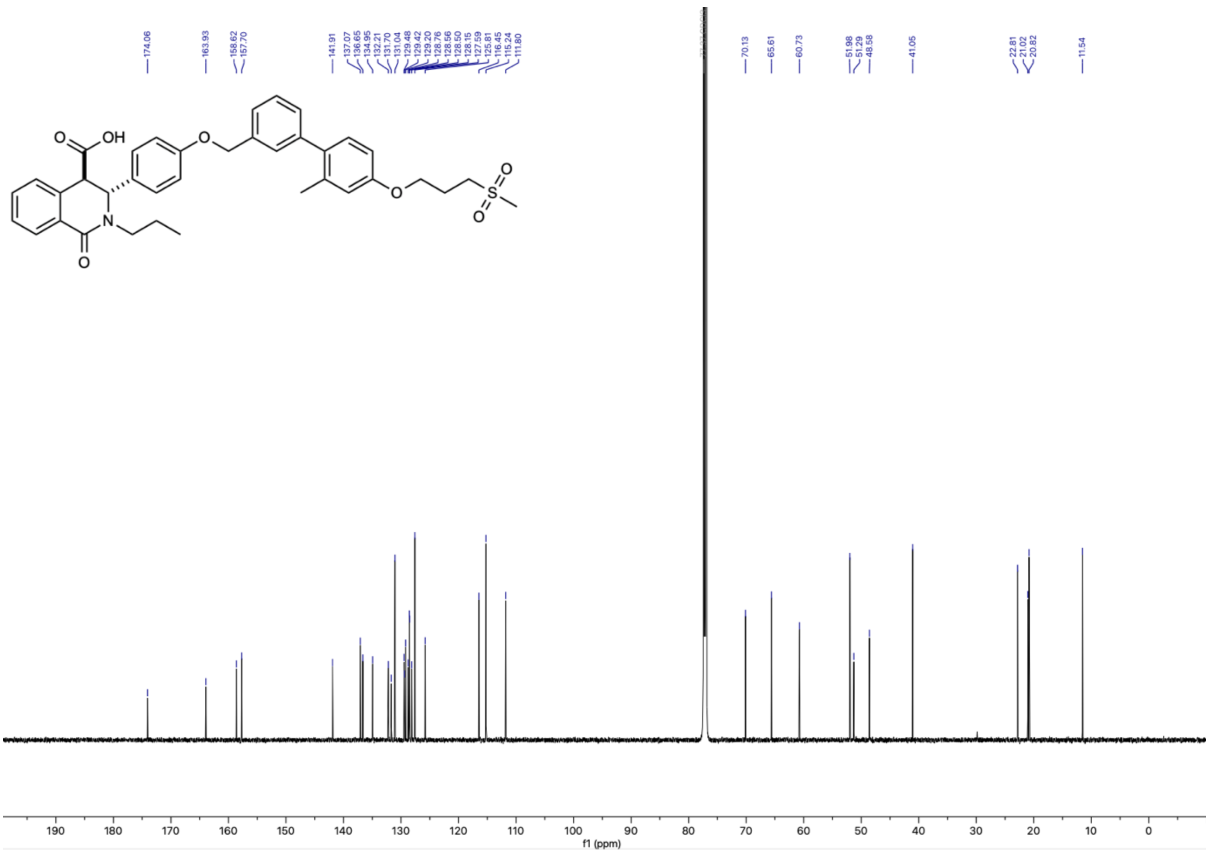

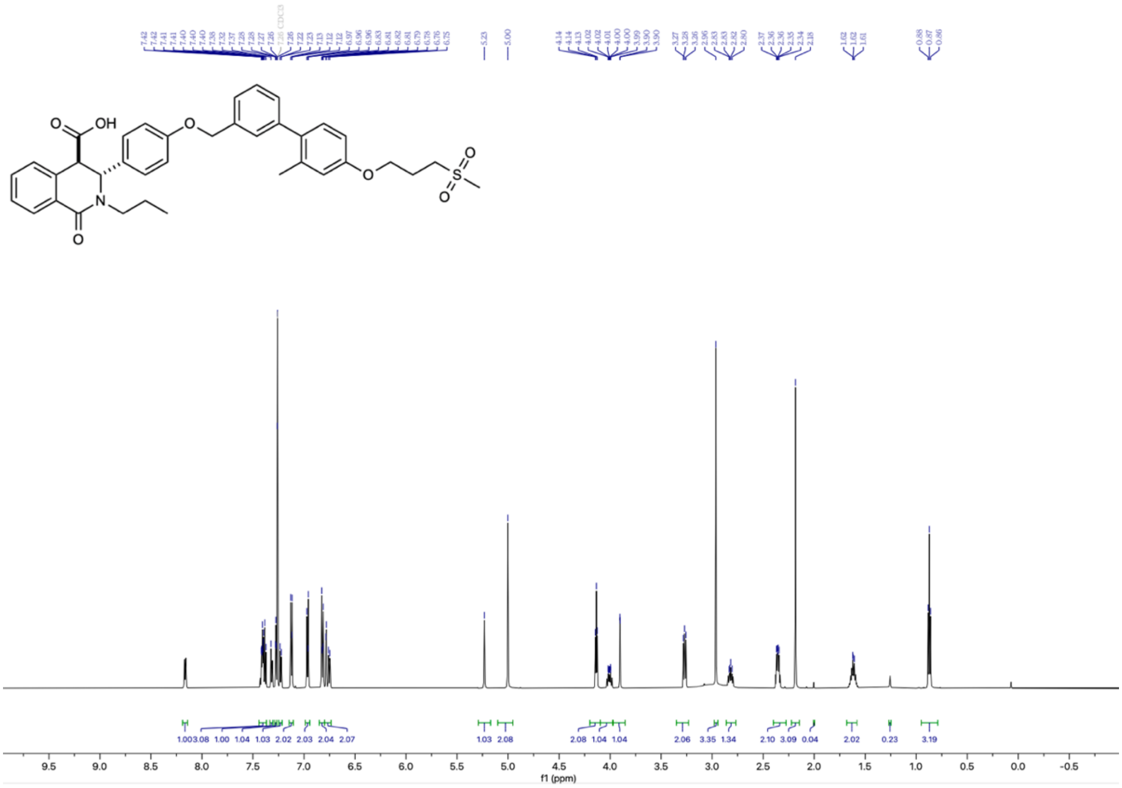

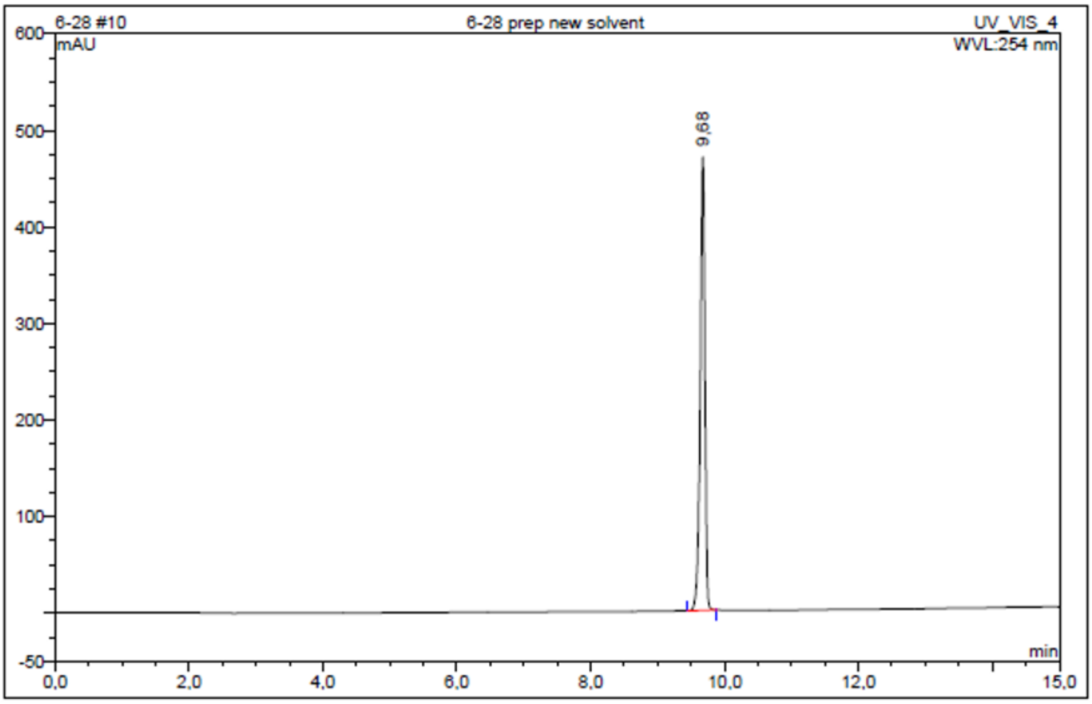


| No. | Ret.Time min | Peak Name | Height mAU | Area mAU*min | Rel.Area % | Amount | Resolution(EP) |
|---|---|---|---|---|---|---|---|
| 1 | 9,68 | n.a. | 469,829 | 38,523 | 100,00 | n.a. | n.a. |
| Total: | | | 469,829 | 38,523 | 100,00 | 0,000 | |

***Figure S32.*** $^{1}H$ NMR, $^{13}C$ NMR and HPLC chromatogram of **TUG-2745**.

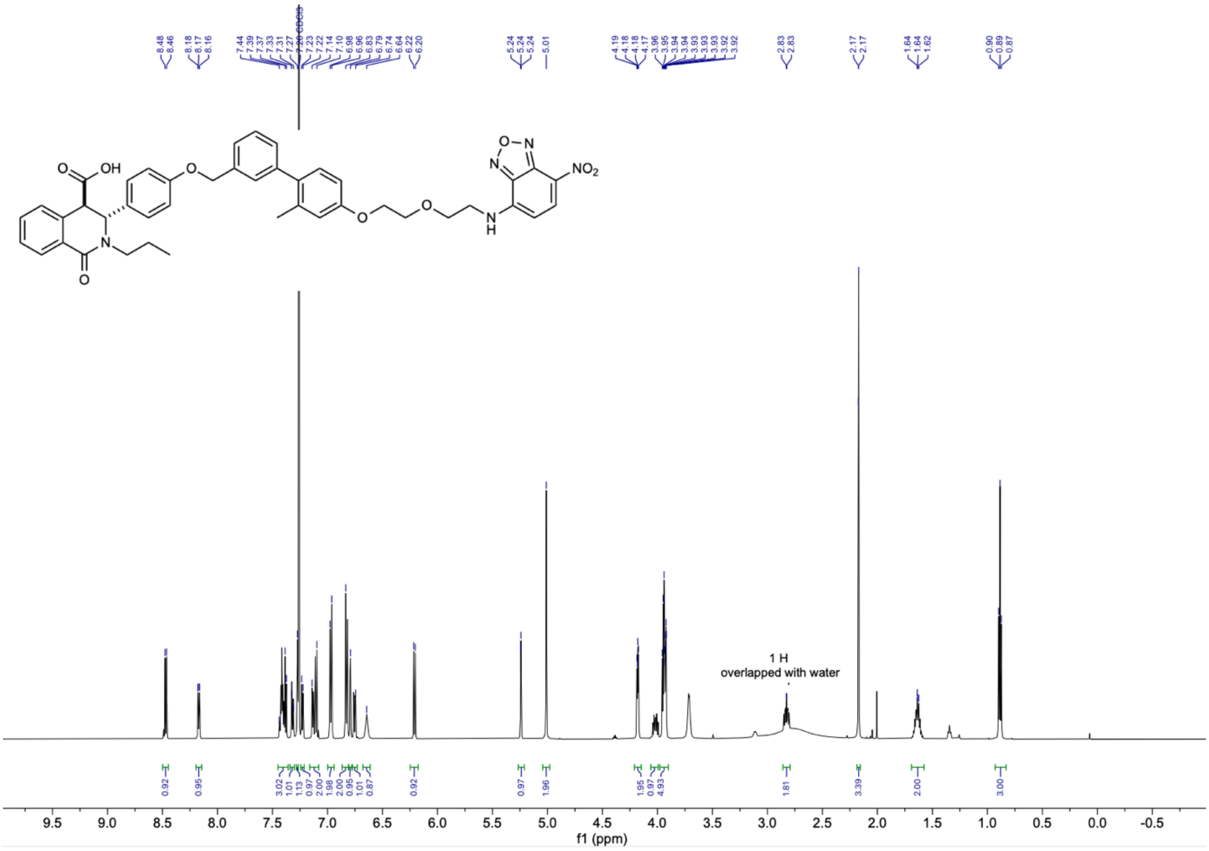

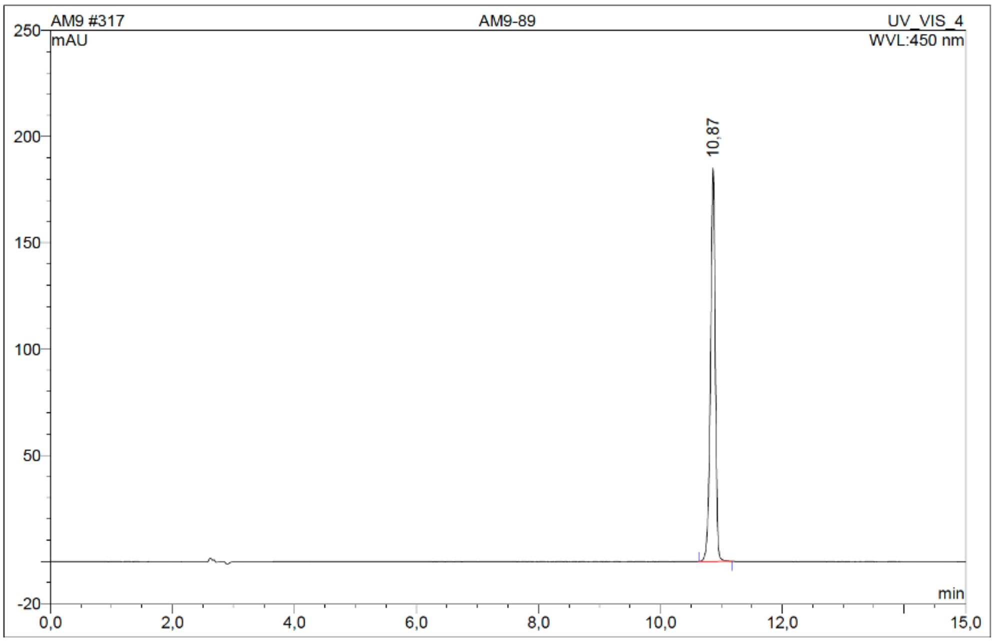


| No. | Ret.Time min | Peak Name | Height mAU | Area mAU*min | Rel.Area % | Amount | Resolution(EP) |
|---|---|---|---|---|---|---|---|
| 1 | 10,87 | n.a. | 185,360 | 16,969 | 100,00 | n.a. | n.a. |
| Total: | | | 185,360 | 16,969 | 100,00 | 0,000 | |

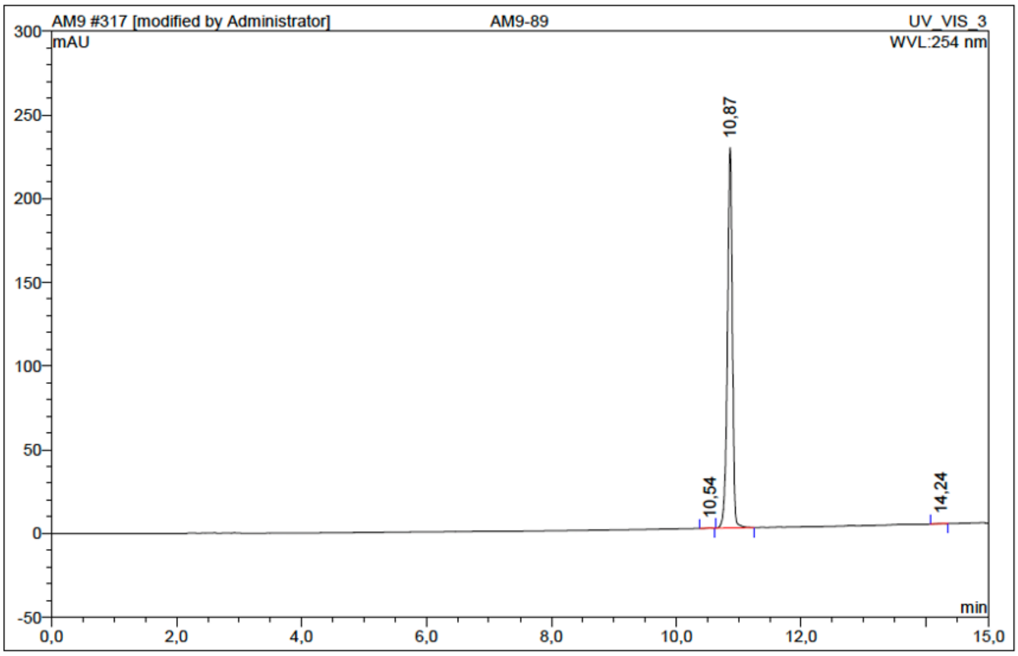


| No. | Ret.Time min | Peak Name | Height mAU | Area mAU*min | Rel.Area % | Amount | Resolution(EP) |
|---|---|---|---|---|---|---|---|
| 1 | 10,54 | n.a. | 0,257 | 0,021 | 0,10 | n.a. | 2,42 |
| 2 | 10,87 | n.a. | 227,347 | 20,844 | 99,78 | n.a. | 19,43 |
| 3 | 14,24 | n.a. | 0,188 | 0,024 | 0,12 | n.a. | n.a. |
| Total: | | | 227,791 | 20,890 | 100,00 | 0,000 | |

***Figure S33.*** [1]H NMR and HPLC chromatograms of **TUG-2884.**
